# Preventing Jacob-induced transcriptional inactivation of CREB protects synapses from β-amyloid in Alzheimer’s Disease

**DOI:** 10.1101/2020.01.08.898304

**Authors:** Katarzyna M. Grochowska, Guilherme M. Gomes, Rajeev Raman, Rahul Kaushik, Liudmila Sosulina, Hiroshi Kaneko, Anja M. Oelschlegel, PingAn Yuanxiang, Irene Reyes-Resina, Gonca Bayraktar, Sebastian Samer, Christina Spilker, Marcel S. Woo, Markus Morawski, Jürgen Goldschmidt, Manuel A. Friese, Steffen Rossner, Gemma Navarro Brugal, Stefan Remy, Carsten Reissner, Anna Karpova, Michael R. Kreutz

**Affiliations:** RG Neuroplasticity, Leibniz Institute for Neurobiology; Magdeburg, Germany; Leibniz Group ‘Dendritic Organelles and Synaptic Function’, Center for Molecular Neurobiology (ZMNH), University Medical Center Hamburg-Eppendorf; Hamburg, Germany; Center for Behavioral Brain Sciences, Otto von Guericke University; Magdeburg, Germany; Department of Cellular Neuroscience, Leibniz Institute for Neurobiology; Magdeburg, Germany; German Center for Neurodegenerative Diseases (DZNE); Magdeburg, Germany; Institute of Neuroimmunology and Multiple Sclerosis, Center for Molecular Neurobiology (ZMNH), University Medical Center Hamburg-Eppendorf; Hamburg, Germany; Molecular Imaging in Neurosciences, Paul Flechsig Institute of Brain Research; Leipzig, Germany; Department of Systems Physiology of Learning and Memory, Leibniz Institute for Neurobiology; Magdeburg, Germany; Department of Biochemistry and Physiology, Faculty of Pharmacy and Food Science, University of Barcelona, Barcelona, Spain and Institut de Neurociències de la Universitat de Barcelona; Barcelona, Spain; Institute of Anatomy and Molecular Neurobiology, Westfälische Wilhelms-University; Münster, Germany

**Keywords:** Alzheimer’s disease, Amyloid pathology, CREB, early synaptic dysfunction, Jacob/NSMF

## Abstract

Synaptic dysfunction caused by soluble β-Amyloid (Aβ) is a hallmark of the early stage of Alzheimer’s disease (AD) and is tightly linked to cognitive decline. Aβ induces by yet unknown mechanisms disruption of transcriptional activity of cAMP– responsive element-binding protein (CREB), a master regulator of cell survival and plasticity-related gene expression. Here, we report that Aβ elicits cytonuclear trafficking of Jacob, a protein serves as a mobile signaling hub that docks a signalosome to CREB, which induces transcriptional inactivation and subsequent synapse impairment and eventually loss in AD. The small chemical compound Nitarsone selectively hinders assembly of this signalosome and thereby restores CREB transcriptional activity. Nitarsone prevents impairment of synaptic plasticity as well as cognitive decline in mouse models of AD. Collectively, the data suggest that targeting Jacob induced CREB shutoff is a therapeutic avenue against early synaptic dysfunction in AD.

## INTRODUCTION

Soluble oligomeric Aβ induces deterioration of synaptic function in AD even before overt signs of dementia and plaque formation(Selkoe, 2002, Selkoe & Hardy, 2016, Forner et al., 2017, Li & Selkoe, 2020, Peng et al., 2022). While a large number of synaptic proteins have been suggested as Aβ receptors (Jarosz-Griffiths et al., 2016), their pathophysiological relevance for synaptic dysfunction *in vivo* is still elusive given that the earliest hallmark of AD in humans and animal models is neuronal hyperexcitability caused by suppression of glutamate reuptake(Zott et al., 2019). In this scenario, glutamate spillover to perisynaptic sites might cause detrimental activation of extrasynaptic N-Methyl-D-Aspartate receptors (NMDAR). NMDAR are heteromeric glutamate-gated ion channels implicated in synaptic plasticity, learning and memory but also in neurodegeneration and excitotoxicity (Hardingham & Bading, 2010, Parsons & Raymond, 2014, Bading, 2017).

Aberrant and synergistic activation of GluN2B-containing NMDAR at extrasynaptic sites by glutamate and Aβ has been shown in AD (Bordji et al., 2010, Malinow, 2012, Bading, 2017, Marcello et al., 2018). Signaling downstream of synaptic and extrasynaptic NMDAR is tightly and antagonistically coupled to the transcription factor CREB. Activation of synaptic NMDAR activates CREB through sustained phosphorylation of a crucial serine at position 133 (S133) and thereby promotes the expression of plasticity-related genes (Hardingham & Bading, 2010) critically involved in learning and memory (Barco et al., 2002, Carlezon et al., 2005). Conversely, predominant activation of extrasynaptic NMDAR leads to sustained dephosphorylation of CREB (CREB shutoff), rendering CREB transcriptionally inactive (Hardingham et al., 2002). Loss of CREB-dependent gene expression after extrasynaptic NMDAR activation precedes cell death and neurodegeneration, however, whether Aβ-induced CREB shutoff plays a role already in early synaptic dysfunction driving cognitive impairment in AD is currently unclear (Espana et al., 2010, Saura & Valero, 2011, Yiu et al., 2011, Teich et al., 2015, Bartolotti et al., 2016). Under the premise that activation of extrasynaptic NMDAR happens before the manifestation of clinical symptoms, it is likely that Aβ will interfere already at this very early stage with transcriptional regulation. Surprisingly little is known, however, on molecular mechanisms of CREB shutoff in general and in AD in particular.

In previous work, we found that the synapto-nuclear protein messenger Jacob, following long-distance transport and nuclear import, differentially transduces NMDAR signals of synaptic and extrasynaptic origin to the nucleus (Dieterich et al., 2008, Karpova et al., 2013, Panayotis et al., 2015, Grochowska et al., 2021). Activation of synaptic NMDARs leads to phosphorylation of Jacob at serine 180 (S180) (pJacob) via MAP-kinase ERK1/2, which is followed by trafficking of a pJacob/pERK1/2 signalosome along microtubules to neuronal nuclei (Karpova et al., 2013). Binding of the intermediate filament α-internexin protects pJacob and pERK against dephosphorylation during transport. This signalosome promotes CREB phosphorylation at S133 and hence CREB-dependent gene expression (Karpova et al., 2013). On the contrary, activation of extrasynaptic NMDAR leads to prominent nuclear translocation of non-phosphorylated Jacob. In this case nuclear import is followed by stripping of synaptic contacts, simplification of dendritic arborization, and cell death (Rönicke et al., 2011, Gomes et al., 2014, Grochowska et al., 2017) and is correlated with CREB shutoff. The molecular identity of the signalosome assembled by non-phosphorylated Jacob is not known and the molecular mechanism of CREB shutoff is at present elusive.

In the present study, we show the involvement of Jacob in Aβ pathology and present a molecular mechanism of CREB shutoff. In addition, we demonstrate the relevance of the identified mechanism for early synaptic failure in AD by targeting a crucial protein interaction responsible for CREB shutoff with the small chemical compound Nitarsone. Taken together, the data support the hypothesis that Jacob operates as a mobile signalling hub that docks NMDAR-derived signalosomes to nuclear target sites andthe results point to the significance of macromolecular protein transport from NMDAR to the nucleus for disease progression at an early stage of AD. Finally, our findings suggest that this pathway provides novel molecular entry points for interventions.

## RESULTS

### CREB shutoff and reduced pJacob levels in AD patient brains

We first examined the levels of pJacob and pan Jacob in post mortem tissue of AD patients (see Table S1 for patient information) to provide evidence for a potential involvement of the protein in human AD pathology. Immunoblotting of a nuclear enriched fraction obtained from the temporal cortex of AD patients did not reveal a significant reduction in pan Jacob levels as compared to controls (Fig EV1A-C). However, the levels of pJacob were significantly reduced by roughly 40% (Fig 1A, B; Fig EV1A), indicating that nuclear import of Jacob following activation of synaptic NMDAR is diminished probably at the expense of activation of extrasynaptic NMDAR (Karpova et al., 2013). Expectedly, we observed significant neuronal loss in AD patients as evidenced by NeuN-immunoblotting (Fig EV1D, E). Since Jacob, unlike CREB, is only detectable in neurons (Mikhaylova et al., 2014), we could compare phosphorylation of nuclear Jacob normalized to total protein levels and corrected these values for NeuN content to adjust for neuronal cell loss (Fig 1C). With this measure we observed a clear reduction of the pJacob/Jacob ratio (Fig 1C) that was correlated with the degree of CREB shutoff, which we determined following FACS sorting of NeuN-positive nuclei (Fig 1D-F Fig EV1F), given that CREB is expressed in both, glia cells and neurons. Taken together, the data indicate CREB shutoff in human AD brains. In addition, lower levels of pJacob suggest a functional link between CREB shutoff and Aβ-pathology that is mediated by Jacob.

**Fig. 1.**
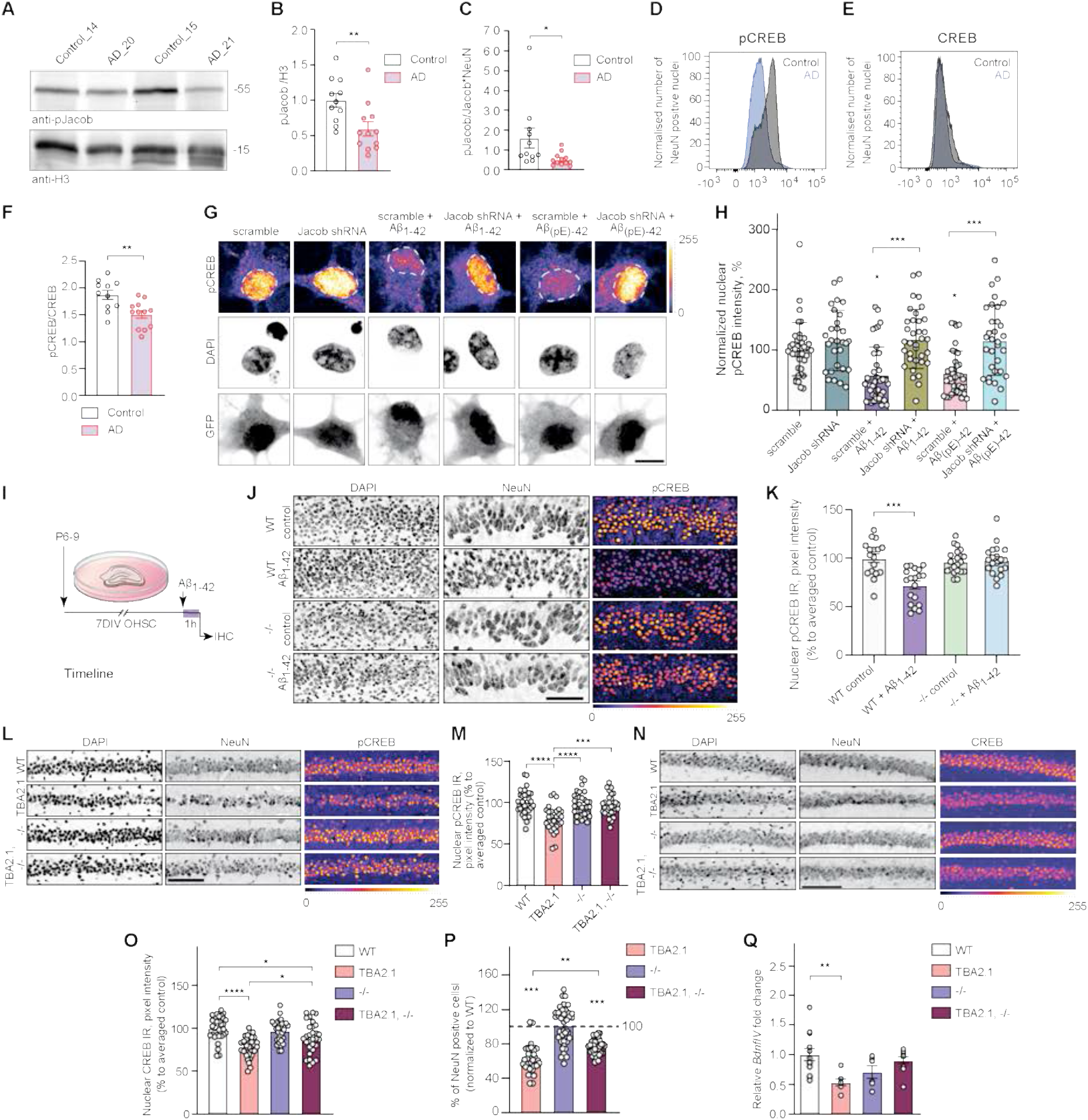
Jacob is associated with CREB shutoff in human AD and in AD mouse models. **(A-C)** Jacob and CREB expression in human AD. **(A, B)** Total pJacob protein levels and (C) pJacob/Jacob ratio corrected by NeuN are significantly reduced in the temporal cortex of AD patients as compared to the age-matched control group. All samples are normalized to Histone3 (H3). N=11-12 different subjects. **(D-F)** FACS of neuronal nuclei revealed significantly decreased pCREB, but not CREB, immunoreactivity in AD patients as compared to the control group. (D, E) Frequency distribution plot of neuronal nuclei immunoreactivity of (d) pCREB and (E) CREB. (F) pCREB/CREB ratio in neuronal nuclei. N=11-12 different subjects. **(G, H)** Jacob shRNA knockdown prevents Aβ-induced CREB shutoff. (G) Representative confocal images of hippocampal neurons transfected with Jacob-shRNA or scrambled (scr) shRNA control (both expressing GFP) and treated with oligomeric preparations of Aβ1-42 or Aβ3(pE)-42. (H) Contrary to the scr shRNA, neurons transfected with a Jacob knockdown construct did not display reduction of pCREB staining intensity after treatment with Aβ1-42 or Aβ3(pE)-42. Bar plot of mean nuclear pCREB intensity normalized to untreated control. Scale bar: 10 µm. N=29-39 nuclei from 2 independent experiments. **(I-K)** Acute (1h) Aβ_1-42_ treatment does not induce CREB shutoff in organotypic hippocampal slices from Jacob (−/−) mice. (J) Representative confocal images of slices immunolabeled against pCREB, co-labeled with NeuN and DAPI. (K) Bar plot of pCREB, N=17-21 slices. **(L, M)** The quantification of pCREB intensity in NeuN-positive cells revealed a statistically significant decrease in pCREB immunoreactivity in TBA2.1 but not in double transgenic animals (TBA2.1, −/−). (L) Representative confocal images of CA1 cryosections from 13 weeks old mice stained for NeuN, DAPI and pCREB. Scale bar: 100 µm. Data represented as cumulative frequency distribution. (M) Bar plot of average hippocampal pCREB nuclear immunoreactivity normalized to WT. N=32-34 hippocampal sections from 7-9 animals. **(N, O)** The quantification of CREB intensity in NeuN positive cells revealed a statistically significant decrease in CREB immunoreactivity in Jacob/*Nsmf* knockout (−/−) and TBA2.1 x Jacob/*Nsmf* knockout (TBA2.1, −/−) mice. (N) Representative confocal images of CA1 cryosections from 13 weeks old mice stained for NeuN, DAPI and CREB. Scale bar: 100 µm. (O) Bar plot of CREB nuclear immunoreactivity normalized to WT. N= 30-32 hippocamal images from 7-9 animals. **(P)** Double transgenic TBA2.1 Jacob/*Nsmf* knockout (TBA2.1, −/−) mice display significantly lower neuronal loss compared to TBA2.1 mice. The number of NeuN positive cells was normalized to WT group. N=31-42 CA1 images analyzed from 9-11 animals per genotype. **(Q)** Jacob knockout rescues decrease in the *BdnfIV* gene transcription. Bar plot of mean *BdnfIV* transcript levels in hippocampal homogenates normalized to *β-actin* as reference transcript. N=5-10 hippocampi. (G, J, L, N) Lookup table indicates the pixel intensities from 0 to 255. *p<0.05, **p<0.01, ***p<0.001, ****p<0.0001 by (b, c, f) two-tailed Student t-test or (H, K, M, O, P, Q) two-way ANOVA followed by Bonferroni’s multiple comparisons test. All data are represented as mean ± SEM.

### Jacob protein knockdown and gene knockout protects against Aβ toxicity

Aβ oligomers can be found in various, post-translationally modified forms, out of which the N-terminally truncated, pyroglutamylated Aβ_3(pE)-42_ species are prominent in the brain of AD patients (Bayer & Wirths, 2014, Kummer & Heneka, 2014). Previous work suggests that Jacob might play a role in Aβ-induced CREB shutoff that is elicited by activation of extrasynaptic GluN2B containing NMDAR by yet unknown mechanisms (Rönicke et al., 2011, Gomes et al., 2014, Grochowska et al., 2017). Knockdown of Jacob by shRNA in hippocampal neurons indeed prevented CREB shutoff induced by treatment of cultures with 500 nM Aβ_1-42_ or Aβ_3(pE)-42_ oligomers (Fig 1G, H). Similar results were obtained in organotypic hippocampal slices from Jacob/*Nsmf* knockout (−/−) mice or wild-type littermates treated with 1 µM oligomeric Aβ_1-42_ (Fig 1I-K). Basal pCREB immunofluorescence levels were not different between both genotypes, however, neurons from knockout mice, unlike the wild-type, did not display Aβ_1-42_-induced CREB shutoff (Fig 1j, K). Total CREB levels remained unchanged (Fig EV1G, H).

### Jacob gene knockout ameliorates neuronal loss in transgenic AD mice

We next reasoned that the lack of Aβ-induced CREB shutoff in Jacob knockout mice could confer neuroprotection in AD. The CA1 subfield of the hippocampus is one of the areas earliest affected in AD, with pronounced neuronal loss and a decreased number of synaptic contacts (Price et al., 2001, Yiu et al., 2011, Padurariu et al., 2012, Wirths & Zampar, 2020). TBA2.1 mice express Aβ_3(pE)-42_ and display very early on severe CA1 neuronal loss, amyloidosis, LTP impairment, and neuroinflammation (Alexandru et al., 2011). We chose TBA2.1 mice because they show probably the most aggressive and prominent amyloid pathology of all transgenic AD mouse models. Western blot analysis of protein extracts from TBA2.1 mouse brain revealed that, while Jacob protein levels remained unchanged (Fig EV1I, J), pJacob levels are decreased resulting in a reduced pJacob/Jacob ratio like in human brain (Fig EV1I-L). In accordance with reports from other AD transgenic mouse lines (Caccamo et al., 2010, Yiu et al., 2011, Bartolotti et al., 2016), we found that TBA2.1 mice exhibit significantly reduced nuclear pCREB levels (Fig 1l, M). To directly study whether the loss of Jacob expression in neurons confers neuroprotection in TBA2.1 mice, we next crossed both lines to obtain homozygous TBA2.1 and Jacob/*Nsmf* (−/−) mice. Interestingly, the double transgenic animals (TBA2.1 x Jacob/*Nsmf* −/−) did not display CREB shutoff as evidenced by no reduction in pCREB levels (Fig 1l-O). Although we observed in all three genotypes (TBA2.1, Jacob/*Nsmf* −/−, and double TBA2.1 x Jacob/*Nsmf* −/−) slightly decreased nuclear CREB levels (Fig 1N, O), the absence of CREB shutoff in Jacob/*Nsmf* −/− and double transgenic animals points to a key role of the protein for transcriptional inactivation at an early stage of Aβ-amyloidosis. Accordingly, cell-loss in the dorsal CA1 region was less pronounced in double transgenic mice (on average 23%) (Fig 1P). The rescue mediated by Jacob gene knockout was also visible at the level of brain-wide network activation patterns when we imaged cerebral blood flow (CBF) in unrestrained behaving mice of all four genotypes (TBA2.1, Jacob/*Nsmf* −/−, TBA2.1 x Jacob/*Nsmf* −/−, and wild-type (WT)) using SPECT (Kolodziej et al., 2014, Oelschlegel & Goldschmidt, 2020) (Fig EV1M). Decreases in CBF, found in dorsal CA1 (arrow) of TBA2.1 mice when compared to wild type animals, were partially rescued in double transgenic mice (Fig EV1M). This rescue was also apparent in the lateral septum and the diagonal band, regions connected to the hippocampus (Fig EV1N). In addition, levels of BDNF mRNA transcribed from promoter IV of the *Bdnf* gene (*BdnfIV*), a synaptic plasticity related neurotrophic factor (Spilker et al., 2016), whose expression is regulated by CREB in an activity-dependent manner, were decreased in TBA2.1 mice, but not in Jacob/*Nsmf* (−/−) and double transgenic animals (Fig 1Q).

Jacob gene deletion did not influence the number of astrocytes (Fig EV1O, P) and activated microglia (Fig EV1O, Q). Moreover, amyloid load, evidenced by the number of Aβ-positive deposits (Fig EV1R, S), was not affected as well, indicating that indirect effects of neuroinflammation or amyloid deposition do not account for the neuroprotection conferred by Jacob gene deletion.

### Jacob is a direct binding partner of CREB and LMO4

These data collectively suggest that <u>Ja</u>cob-induced <u>C</u>REB <u>s</u>hutoff that we termed JaCS contributes to transcriptional inactivation of CREB in AD and we therefore next aimed to decipher underlying molecular mechanism. We first tested for a possible direct interaction between both proteins. A pull-down assay with bacterially expressed proteins revealed a direct association of both N-terminal 117-172 amino acid (aa) and C-terminal (262-532 aa) regions of Jacob to the bZIP domain of CREB (Fig 2A, Fig EV2A-C). Accordingly, super-resolution stimulated emission depletion (STED) imaging showed nuclear Jacob in close proximity to CREB in cultured hippocampal neurons (Fig EV2D, E). We could co-immunoprecipitate endogenous CREB from HEK293T cells following heterologous expression of Jacob (Fig EV2F) and in support of these data we found prominent *in vivo* FRET efficiency when we co-expressed either full-length or the N-terminal half of Jacob and a C-terminal fragment of CREB (Fig 2B-E). Of note, the N-terminal fragment of Jacob yielded significantly stronger fluorescence resonance energy transfer (FRET) signals than the C-terminal fragment (Fig 2B-F).

**Fig. 2.**
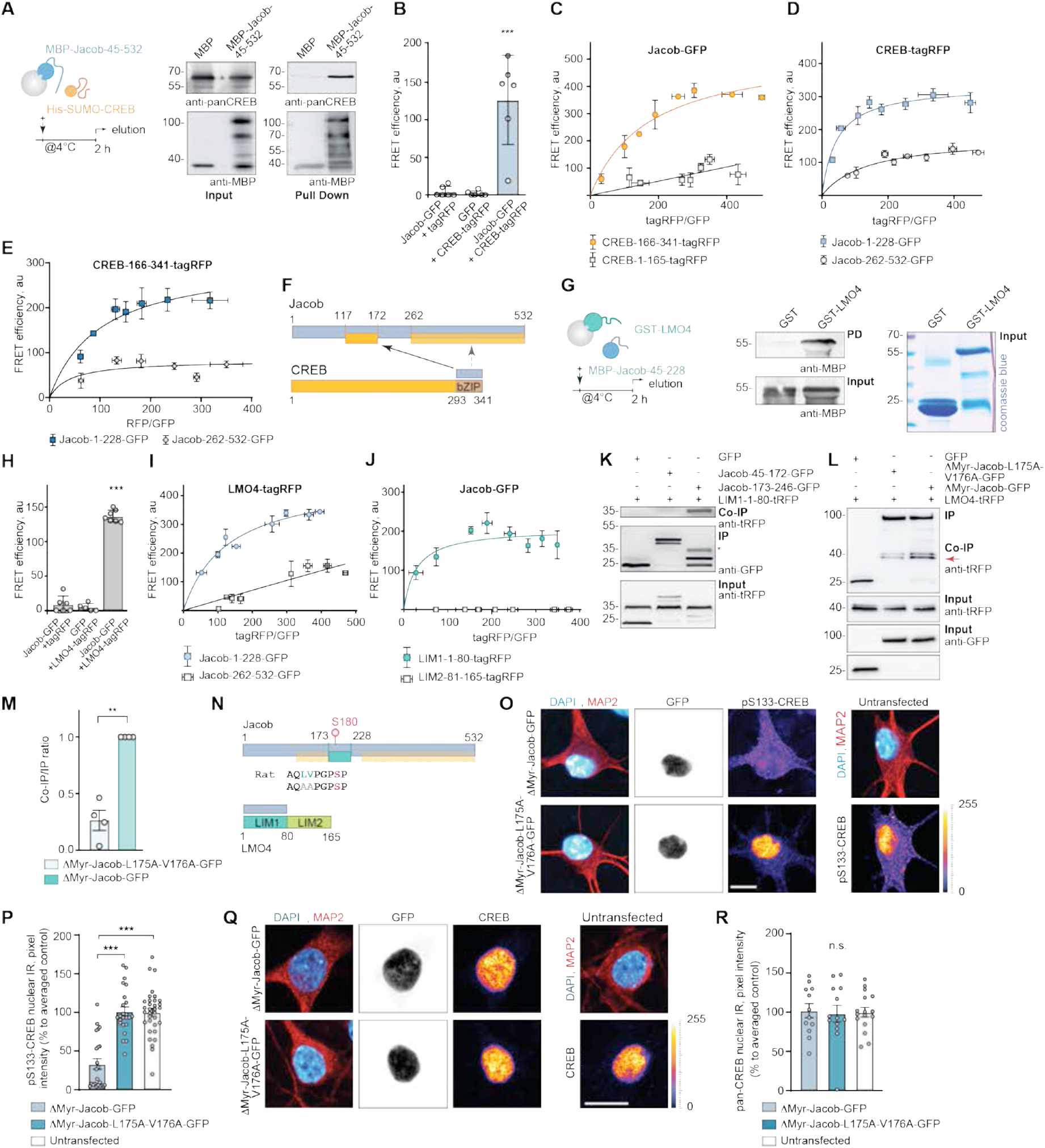
Jacob directly associates with CREB and LMO4. **(A)** Pull-down assays confirm a direct interaction of MBP-Jacob-45-532 and His-SUMO-CREB. **(B)** FRET measurements show an association between CREB-tagRFP and Jacob-GFP. N=5 independent experiments measured in triplicates. **(C)** The C-terminus (CREB-166-341-tagRFP) but not the N-terminus (CREB-1-165-tagRFP) of CREB closely associates with Jacob-GFP in FRET saturation experiments. **(D)** Both the N-(Jacob-1-228-GFP) and the C-terminus (Jacob-262-532-GFP) of Jacob are in close proximity to CREB-tagRFP, however, the Jacob-1-228-GFP association with CREB is significantly stronger. **(E)** The C-terminus of CREB (CREB-166-341-tagRFP) associates prominently with the N-terminus (Jacob-1-228-GFP) and less strong with the C-terminus (Jacob-262-532-GFP) of Jacob. (C-E) FRET efficiency is presented in arbitrary units from 5-6 independent experiments. **(F)** Binding interfaces between CREB and Jacob. **(G)** GST-LMO4 but not GST alone directly binds to MBP-Jacob-45-228. **(H)** FRET experiments revealed that Jacob-GFP interacts with LMO4-tagRFP. **(I, J)** FRET saturation experiments indicate the association of Jacob-1-228-GFP with LIM-1-80-tagRFP. FRET efficiency is presented in arbitrary units as a mean of 6 independent experiments measured in triplicates. **(K)** Co-immunoprecipitation experiments to map the binding region of Jacob to the LIM1 domain of LMO4 revealed the association with Jacob-179-246-GFP, but not with Jacob-45-172-GFP (CREB-binding region). **(L, M)** Heterologous co-immunoprecipitation experiments between LMO4-tagRFP and nuclear ΔMyr-Jacob-GFP, ΔMyr-Jacob-L175A-V176A-GFP or GFP point to a decreased association of ΔMyr-Jacob-L175A-V176A-GFP with LMO4 as compared to ΔMyr-Jacob-GFP. N=4 independent experiments. **(N)** Binding interfaces between Jacob and LMO4. **(O-R)** A Jacob-LMO4-binding mutant expressed in the nucleus does not induce CREB shutoff. (O, Q) Representative confocal images of hippocampal neurons transfected with ΔMyr-Jacob-GFP (Jacob targeted to the nucleus) or ΔMyr-Jacob-L175A-V176A-GFP. Scale bar: 10 µm. Lookup table indicates the pixel intensities from 0 to 255. (P, R) The mean of nuclear (P) pCREB or (R) immunoreactivity in Jacob-expressing neurons was normalized to non-transfected controls. N =23-33 neuronal nuclei from two independent cell cultures. **p<0.01, ***p<0.001, ****p<0.0001 by (M) one-sample t-test or one-way ANOVA followed by (B, H) Bonferroni’s or (P, R) Tukey’s multiple comparisons test. All data are represented as mean ± SEM.

In a yeast two-hybrid (YTH) screen performed with the N-terminus of Jacob as bait we identified LMO4 as a binding partner (Fig EV3A). LMO4 is a transcriptional co-activator of CREB (Kashani et al., 2006) and we therefore wondered whether the Jacob-LMO4 interaction has a role in JaCS. A region encompassing aa 117-228 of Jacob directly interacted with the LIM1 domain of LMO4 (Fig 2G, Fig EV3B, C). In addition, both proteins were in close proximity to each other in neuronal nuclei (Fig EV3D, E) and the LIM1 domain co-localized with cytosolic Jacob clusters following heterologous expression (Fig EV3F). The association of both proteins was further confirmed by heterologous co-immunoprecipitation with tag-specific antibodies (Fig EV3G) and the direct interaction was corroborated by *in vivo* FRET analysis (Fig 2H-J). Subsequent heterologous co-immunoprecipitation of Jacob fragments with the LIM1 domain indicated that the binding regions for LMO4 and CREB do not overlap (Fig 2F, K). Interestingly, like other LIM1 domain binding proteins, Jacob contains a leucine- and valine-rich stretch (Joseph et al., 2014). Point mutations (L175A-V176A) within this region resulted in much weaker binding (Fig 2 L-N). In previous work we found that nuclear overexpression of non-phosphorylated Jacob leads to dephosphorylation of CREB (Dieterich et al., 2008). Nuclear accumulation of the LMO4 binding mutant of Jacob, however, did not induce CREB shutoff (Fig 2N-R), indicating that the association with LMO4 is instrumental for JaCS.

### Jacob competes with CREB for LMO4 binding

We next asked why the association with LMO4 is crucial for JaCS. *In vivo* FRET assays and heterologous co-immunoprecipitation revealed that the LIM1 domain, which is the binding interface for Jacob (Fig 2I, J, N, Fig EV3A, F, G), also interacts with a N-terminal fragment of CREB (Fig 3A-C). LMO4 directly binds to this region, but not to the isolated bZIP domain of CREB (Fig 3D-F, Fig EV3B, C), which binds Jacob. The direct interaction with LMO4 promotes phosphorylation of S133 and thereby CREB transcriptional activity as evidenced by increased CRE-driven luciferase activity following heterologous expression of LMO4 (Fig 3G). shRNA knockdown of LMO4 (Fig EV3H-J) in neurons resulted in reduced pCREB (Fig 3H, I) but not total CREB immunofluorescence (Fig 3J, K).

**Fig. 3.**
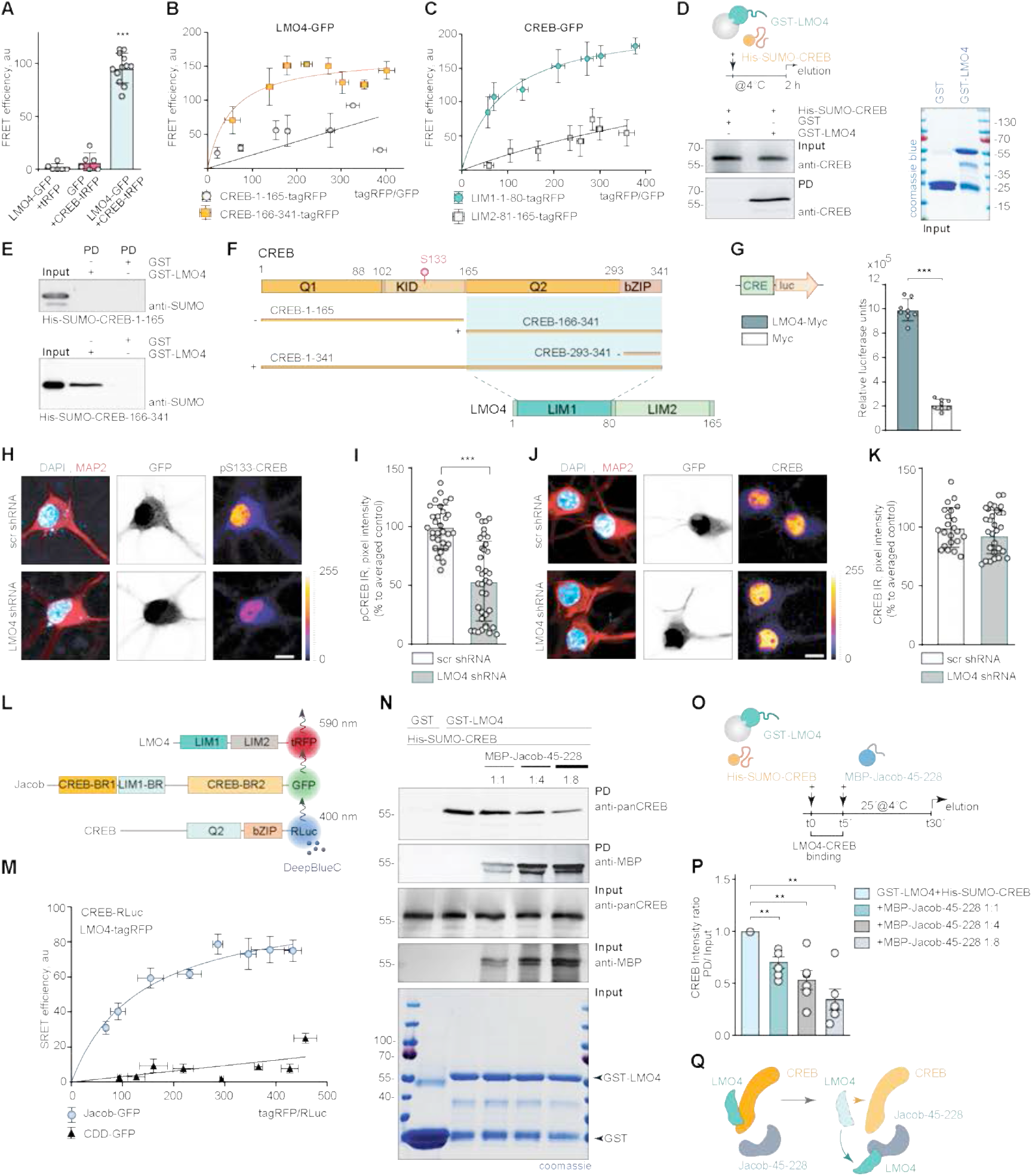
Jacob can displace LMO4 from CREB. **(A)** FRET measurements indicate a tight association between LMO4-GFP and CREB-tagRFP. N=6 independent experiments measured in triplicates. **(B, C)** FRET saturation experiments with (B) LMO4-GFP and CREB-1-165-tagRFP or (C) CREB-166-341-tagRFP and CREB-GFP and LIM1-1-80-tagRFP or LIM2-81- 165-tagRFP revealed an association between the LIM1 domain of LMO4 and the C-terminus of CREB. N=8 independent experiments. **(D)** Recombinant His-SUMO-CREB directly binds to GST-LMO4 in pull-down experiments. **(E)** The interaction between CREB with LMO4 is mediated by the C-terminus of CREB (166-341 aa), but not by its N-terminus (1-165 aa), as evidenced by pull-down experiments between recombinant His-SUMO-1-165-CREB, His-SUMO-166-341-CREB, and GST-LMO4. **(F)** Schematic representation of CREB and LMO4 domain structure and fusion constructs used for the experiments. Light green boxes represent the interaction interface. **(G)** Myc-LMO4 overexpression increases CREB dependent luciferase expression in HEK293T cells expressing luciferase under the CRE promoter. Relative luciferase units in cells overexpressing Myc-LMO4 as compared to Myc-transfected controls. N=8 from two independent experiments. **(H-K)** Knockdown of LMO4 reduces nuclear pCREB immunoreactivity. (H, J) Representative confocal images of hippocampal neurons transfected with LMO4 shRNA construct or scrambled control (both expressing GFP under CMV promoter as a transfection control). Scale bar: 10µm. Dot plots representing the mean of nuclear (I) pCREB or (K) CREB staining intensity normalized to scrambled control. N=30-37 nuclei analyzed from at least 3 independent cell cultures. **(l. M)** SRET saturation experiments reveal that Jacob-GFP forms a triple complex with CREB-RLuc and LMO4-tagRFP. The caldendrin (CDD-GFP) was used as negative control. N=8 independent experiments. (M) Schematic representation of constructs used in SRET experiments. “BR” stands for binding region. **(N-Q)** The N-terminus of Jacob displaces LMO4 from CREB. (N) GST-LMO4 coupled to beads was preincubated with His-SUMO-CREB and subsequently incubated with an increasing amount of MBP-Jacob-45-228. (O) Schematic depicts the timeline of the competition pull-down experiment. (P) Bar graphs represent six independent experiments per condition. (H, J) Lookup table indicates the pixel intensities from 0 to 255. **p<0.01, ***p<0.001, ****p<0.0001 by (G, I, K) two-tailed Student t-test or (P) one-sample t-test or (a) one-way ANOVA followed by Bonferroni’s multiple comparisons test. All data are represented as mean ± SEM.

Direct binding of LMO4 to CREB and Jacob raised the question whether all three proteins can assemble in a trimeric complex or whether they compete for the same binding interface. Sequential resonance energy transfer (SRET) *in vivo* indeed revealed the existence of a triple complex (Fig 3L, M). Jacob harbors a binding interface for the interaction with CREB and LMO4 at the N-terminus (Fig 2) whereas LMO4 can only associate through its first LIM1 domain either to Jacob or to CREB (Fig 2, 3). Subsequent competition pull-down experiments confirmed competitive binding between the N-terminus of Jacob and CREB with LMO4, as shown when GST-LMO4 was coupled to the beads and increasing amounts of the N-terminal fragment of Jacob were added in the presence of CREB (Fig 3N-P). In summary, the LIM1 domain of LMO4 mediates the association either with CREB or Jacob, and Jacob is capable to displace LMO4 from the CREB complex, a mechanism that should facilitate CREB shutoff (Fig 3Q).

### Protein phosphatase 1 γ (PP1**γ)** and LMO4 are involved in Jacob-induced CREB shutoff

We next asked whether LMO4-binding to Jacob might be also actively involved in CREB shutoff. PP1 is a phosphatase that can dephosphorylate S133 in CREB. Jacob harbors several PP1 binding motifs, both proteins co-localize following heterologous expression (Fig EV4A) and tagged Jacob co-immunoprecipitate with endogenous PP1γ (Fig EV4B). Pull-down experiments revealed a direct interaction (Fig 4A, Fig EV4C) and heterologous co-immunoprecipitation experiments pointed to two binding interfaces (Fig 4B). We therefore next addressed whether the association with PP1γ is involved in JaCS. To this end we expressed a phosphodeficient mutant of Jacob in the nucleus of hippocampal primary neurons (Karpova et al., 2013) and incubated cultures with the PP1γ inhibitor okadaic acid (OA). Interestingly, we found that treatment with OA prevented JaCS (Fig 4C, D).

**Fig. 4.**
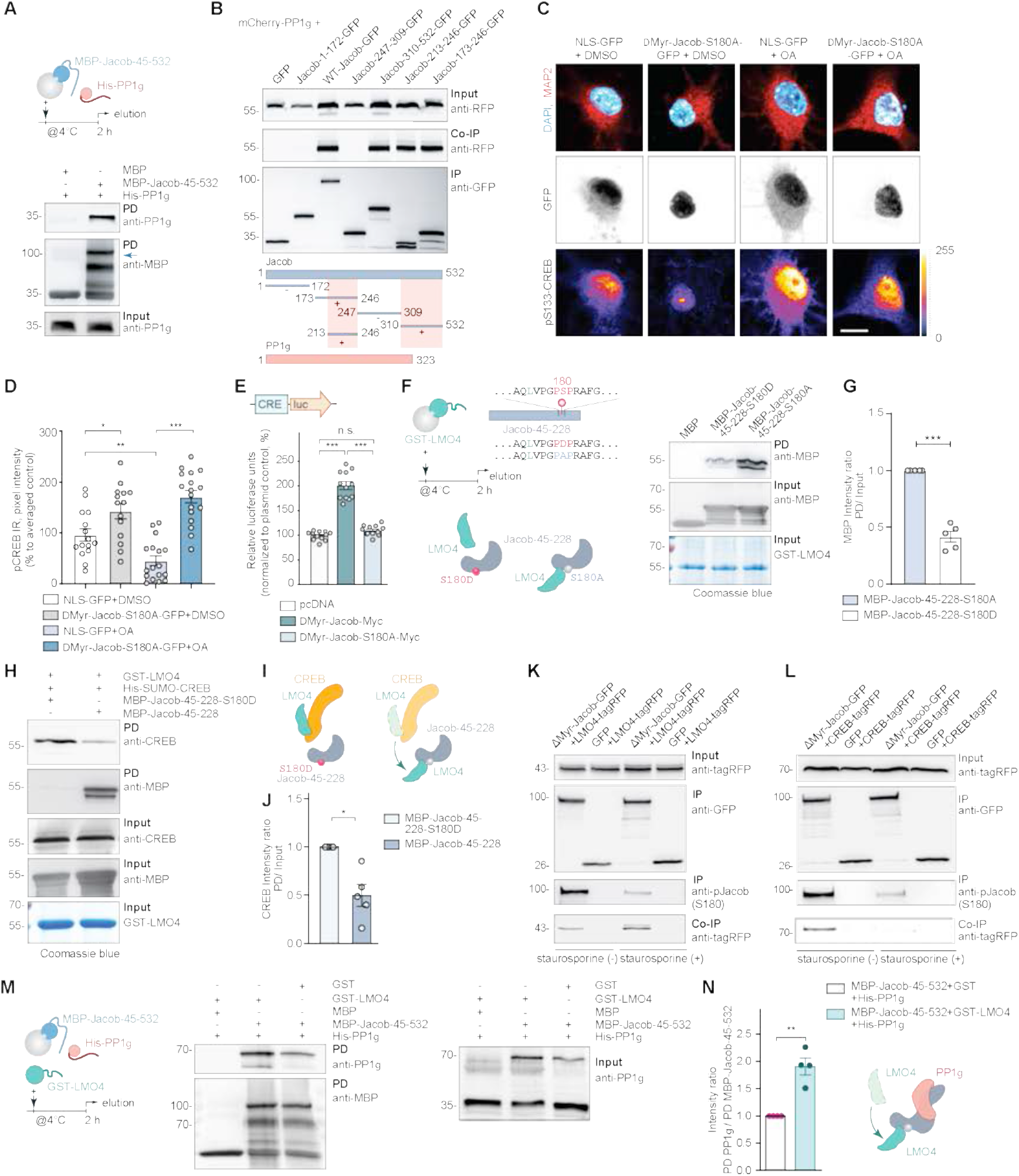
CREB shutoff essentially requires Jacob binding to PP1. **(A)** Pull-down experiments confirmed a direct interaction between recombinant MBP-Jacob-45-532 and His-PP1γ. **(B)** Mapping of the mCherry-PP1γ interaction region within the Jacob sequence revealed binding of a C-terminal fragment (Jacob-310-250-GFP) as well as the N-terminal part (Jacob-173-246-GFP) where the region between 213-246 aa is sufficient for immunoprecipitation. The green boxes in the schematic indicate binding regions. **(C, D)** Treatment of hippocampal primary neurons expressing phosphodeficient Jacob in the nucleus with OA rescues Jacob-induced CREB shutoff. Confocal images of pCREB immunostaining in DIV15 neurons overexpressing ΔMyr-Jacob-S180A-GFP with and without OA treatment. Scale bar: 20 μm. Lookup table indicates the pixel intensities from 0 to 255. N=14-17 nuclei analyzed from two independent cell cultures. **(E)** Overexpression of ΔMyr-Jacob-Myc but not the phospho-deficient mutant (ΔMyr-Jacob-S180A-Myc) positively regulates CREB dependent luciferase expression. N=3 independent experiments. **(F, G)** Phosphodeficient N-terminus of Jacob (MBP-Jacob-45-228-180A) interacts with LMO4 stronger than its phosphomimetic form (MBP-Jacob-45-228-180D). (G) Quantification of MBP immunoreactivity normalized to input. Data are presented from five independent experiments. **(H-J)** Phosphomimetic Jacob mutant (MBP-45-228-180D) does not displace LMO4 from CREB. Recombinant GST-LMO4 was coupled to beads, preincubated with His-SUMO-CREB and subsequently incubated in 1:8 ratio with MBP-Jacob-45-228 or MBP-45-228-180D. (J) Quantification of the CREB band intensity normalized to the input. N=5 independent experiments. **(K)** Treatment with staurosporine decreases Jacob phosphorylation level (S180), but increases its association with LMO4-tagRFP. Immunoblot of HEK293T cells extracts transfected with LMO4-tagRFP and ΔMyr-Jacob-GFP or GFP alone. **(L)** Treatment with staurosporine decreases the association of Jacob with CREB. Immunoblot of HEK293T cells extract transfected with CREB-tagRFP and ΔMyr-Jacob-GFP or GFP as a control. **(M, N)** The association of Jacob with LMO4 enhances its interaction with PP1γ in pull-down assays. (M) PP1γ interacts with Jacob as a dimer (70 kDa) that forms during purification. (N) Bar graph represents quantification of PP1γ immunoreactivity normalized to MBP-Jacob. *p<0.05, **p<0.01, ***p<0.001, ****p<0.0001 by (G, J, N) one-sample t-test or (E) one-way ANOVA followed by Bonferroni’s multiple comparisons test or (D) two-way ANOVA followed by Bonferroni’s multiple comparisons test. All data are presented as mean ± SEM.

Phosphorylation of Jacob at S180 is induced by activation of synaptic NMDAR, whereas nuclear import of non-phosphorylated Jacob is related to extrasynaptic NMDAR activation (Karpova et al., 2013). Accordingly, in a CRE-luciferase activity assay the expression of wild-type but not phosphodeficient Jacob caused increased activity (Fig 4E, Fig EV4D). Furthermore, the association of a phosphodeficient mutant of Jacob to LMO4 was significantly stronger than the corresponding phosphomimetic protein (Fig 4F, G, Fig EV4E, F). Phosphomimetic Jacob, unlike the non-phosphorylated protein, did not displace CREB from LMO4 bound to beads (Fig 4H-J, Fig 3N-Q). To further test the idea that LMO4 binds to non-phosphorylated Jacob more efficiently we applied the protein kinase inhibitor staurosporine and performed heterologous co-immunoprecipitation experiments (Fig 4K, L). We indeed found a stronger association of non-phosphorylated Jacob to LMO4 (Fig 4K, L) and concomitantly a stronger association of S180 phosphorylated Jacob to CREB (Fig 4K, L). Since Jacob directly interacts with PP1γ (Fig 4A) we performed a pull-down assay where we observed that the association between both proteins was much stronger in the presence of LMO4 (Fig 4M, N, Fig EV4E, F). Thus, non-phosphorylated Jacob, entering the nucleus following activation of extrasynaptic NMDAR, will likely displace LMO4 from the CREB complex and the subsequent association with LMO4 will enhance binding to PP1γ, which then ultimately results in CREB shutoff (Fig 4I).

### The small organoarsenic compound Nitarsone selectively blocks binding of Jacob but not of CREB to the LIM1 domain of LMO4

The molecular analysis outlined above allowed us to perform structural modeling of the binding interface between CREB, Jacob and the LIM1 domain of LMO4. To this end we analyzed deposited peptide-bound LMO4 structures. In LMO4:LIM domain-binding protein 1 (Ldb1) (Deane et al., 2004) a peptide of Ldb1 binds to LMO4 by short β-β main chain formations and single hydrophobic side chains protruding into deep pockets in each of the two LIM domains (Fig 5A). LMO4:Ldb1 highlights the relatively weak sequence preferences for peptides binding to either or both LIM domains (Fig 5A). Therefore, we performed a positional scan by computational serial mutation of each peptide residue to any of 20 aa stretches in both Jacob and CREB which let us define a search pattern of only 4 critical residues for potential LMO4 binding regions (Fig EV5A-D). With this approach we found eight potential binding regions in Jacob and five in CREB (Fig EV5B), for which we modeled LIM1:peptide complexes and calculated complex stability. For Jacob we confirmed a peptide including residues 172-186 that is part of the experimentally localized LMO4 binding region. In addition, we found two overlapping peptides within the experimentally determined LMO4 binding region for CREB (Fig EV5D).

**Fig. 5.**
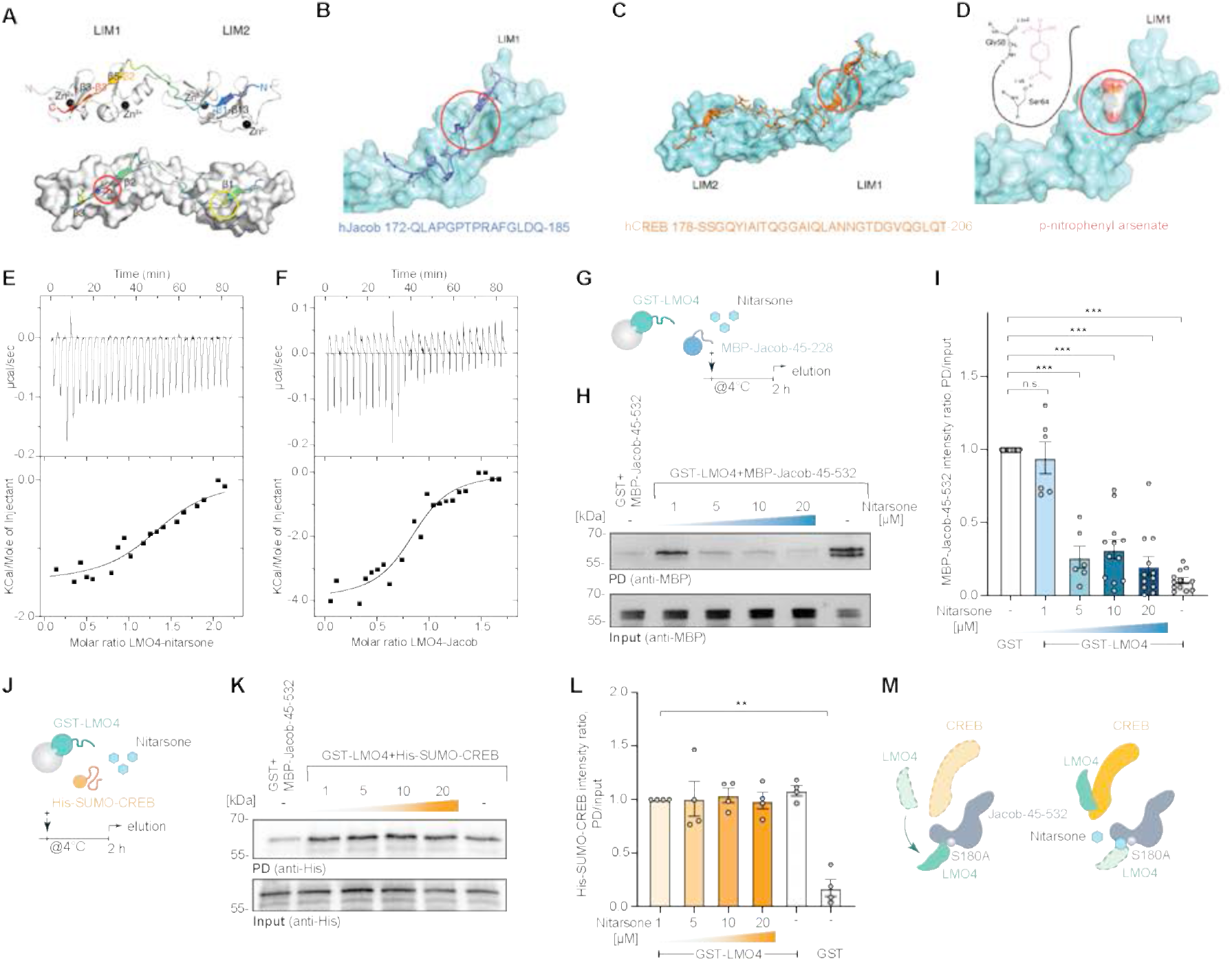
Nitarsone disrupts LIM1-domain binding to Jacob. **(A)** Template structure of an LMO4:peptide complex based on fusion protein LMO4:Ldb1 LID (protein databank (PDB)ID: 1RUT). LIM1-LIM2 tandem domains are folded and stabilized by 4 zinc atoms (black spheres) and bind to a 29 residues long peptide in anti-parallel orientation. The binding occurs mainly via 3 well defined β-strands (β1, β3, β4) interacting with corresponding β-strands of LMO4 (β13, β5, β3). A positional alanine scan highlighted two hydrophobic binding pockets (red & yellow circle, lower panel) as hot spots of that complex allowing only residues Ile (red), Leu, Met, or Val (yellow) to be buried in each of the pockets. The peptide and the residues are rainbow color coded according to the positional alanine scan (ΔΔG (kcal/mol)) from 0% (blue, no side chain effect) to 100% (red, critical conserved residue). **(B)** hJacob residues 172-185 bind to the LIM1 domain. **(C)** hCreb binds to LIM1 and LIM2 similar to Ldb1. **(D)** Nitarsone (p-nitrophenyl arsonic acid) fits to the hydrophobic binding pocket of LIM1 and can form two hydrogen bonds. **(E, F)** ITC analysis of LMO4 (e) nitarsone or (f) Jacob interaction. ITC thermograms for sequential dilutions. Upper panel presents raw data, with heat pulses illustrating exothermic binding. Lower panel depicts binding curve of integrated heat measurements with the best fit using one site binding model. **(G-I)** Nitarsone disrupts binding of Jacob to LMO4 in a concentration-dependent manner. (G) Bacterially expressed GST-LMO4 was immobilized on beads and pre-incubated with increasing concentrations of Nitarsone, and subsequently with MBP-Jacob-45-532. (H) MBP immunoreactivity normalized to input. N=6. (I) Representative immunoblot probed with anti-MBP antibody of input and pull-down with GST as a control. **(J-M)** Nitarsone does not disrupt binding of LMO4 to CREB. (J) Representative immunoblot probed with anti-His of input and pull-down with GST as a control. (K) Bacterially expressed GST-LMO4 was immobilized on beads and was pre-incubated with growing concentrations of Nitarsone, and subsequently with His-Sumo-CREB. (L) His immunoreactivity normalized to input. N=4. (I, L) **p<0.01, ***p<0.001 by one-sample t-test.

We next searched for small molecules that might selectively prevent binding of Jacob and not of CREB to LIM1 of LMO4. Here, we used ZINCPharmer, a tool that allows to define donor and acceptor atoms within the hydrophobic binding pocket (Fig 5B) and the β-strand (β3, Fig 5C) of LIM1 (Koes & Camacho, 2012). Several hits contained a p-nitrophenyl group (e.g. 2-[[1-(4-nitrophenyl)ethyl]amino]ethan-1-ol (ZINC37177221)) fitting to the hydrophobic binding pocket. We therefore next searched the Drugbank database (Wishart et al., 2006) for purchasable drugs containing this group and identified p-nitrophenylarsonic acid (Nitarsone) as promising candidate since Nitarsone fitted into the hydrophobic binding pocket of LIM1 as evidenced by AutoDock Vina (Eberhardt et al., 2021) (Fig 5D).

To prove efficacy, specificity and affinity of Nitarsone binding to the LIM1 domain we next purified recombinant proteins expressed in bacteria (Fig EV5E). Isothermal titration calorimetry revealed a single binding site in LMO4 and a K_D_ of 0.77 µM (Fig 5E), which is roughly matching the K_D_ of 0.37 µM for binding of Jacob to LMO4 (Fig 5F). These results prompted us to test the prediction that Nitarsone will only block binding of LMO4 to Jacob but not to CREB. In GST-pulldown experiments we could indeed show that Nitarsone completely abolished binding of Jacob to LMO4 when applied in 5x times molar excess (Fig 5G-I; Fig EV5E). In these experiments we immobilized 500 nM GST-LMO4, saturated binding with 1 µM Jacob 45-228 and then applied 5 µM Nitarsone. In contrast, even at a concentration of 20 µM, i.e. in 20x times molar excess, the substance did not displace CREB from LMO4 (Fig 5J-M, Fig EV5E).

### Nitarsone application rescues Aβ-induced CREB shutoff as well as synapse loss and synaptic dysfunction

We next found that bath application of 10 µM Nitarsone prevented acute Aβ-induced CREB shutoff in hippocampal primary neurons (Fig EV6A-C). The drug was either applied 30 min prior or 2 h after application of 500 nM Aβ for 2 h. In both conditions we found a rescue of pCREB levels following Nitarsone administration (Fig EV6A-C). Co-application of an even lower dose of 5 µM Nitarsone also rescued CREB shutoff induced by application of 500 nM Aβ oligomers for 48 h (Fig 6A, B).

**Fig. 6.**
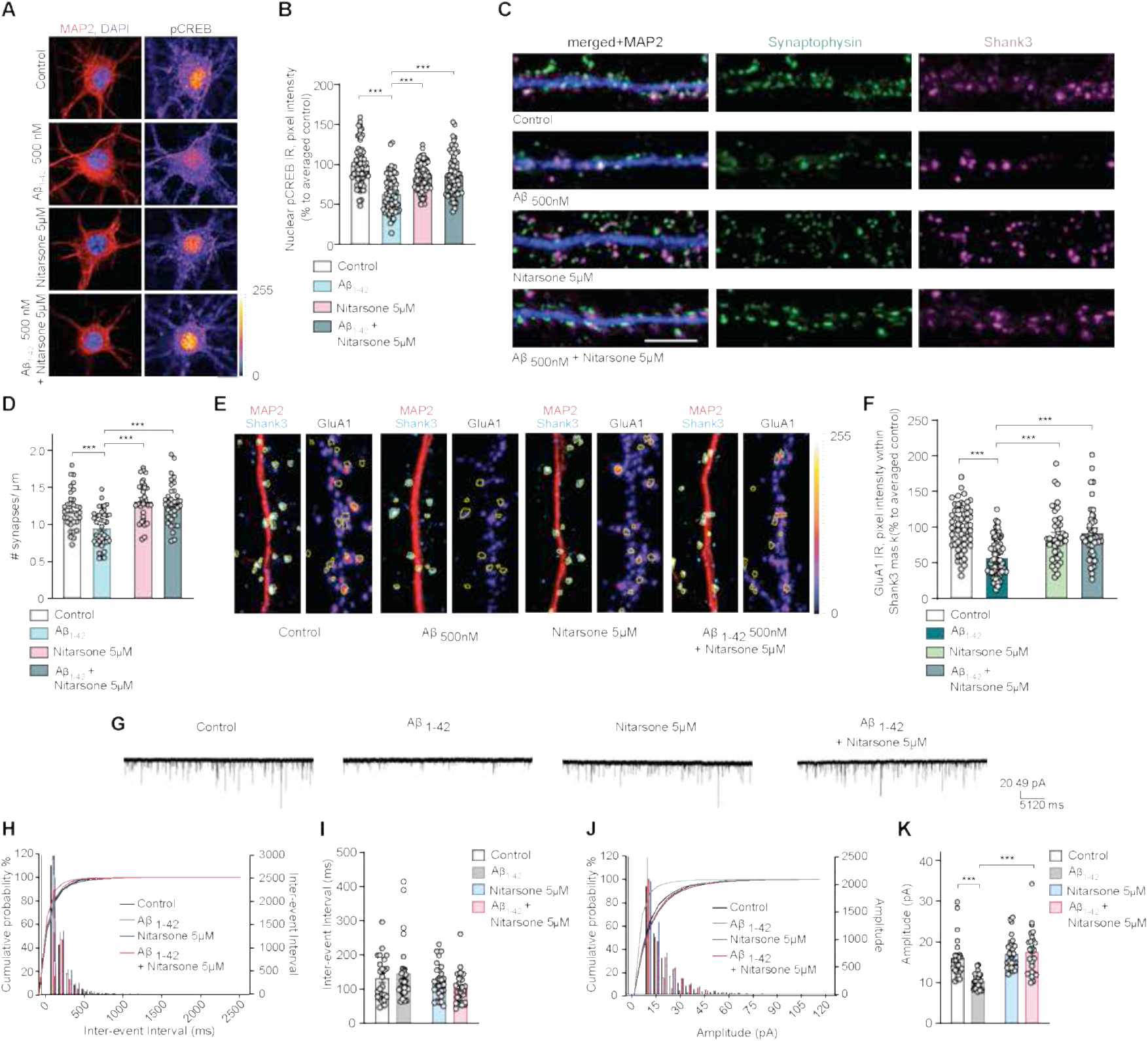
Nitarsone treatment rescues Aβ_1-42_–induced synaptic dysfunction. **(A, B)** Co-application of Nitarsone prevents Aβ-induced CREB shutoff. DIV16 hippocampal cultures were treated with 5 µM Nitarsone, 500 nM Aβ_1-42_, 5 µM Nitarsone with 500 nM Aβ_1-42_ or vehicle control for 48h and stained for pCREB, MAP2, and DAPI. (A) Representative confocal images. Scale bar: 10 µm. (B) Nuclear pCREB immunoreactivity normalized to control. N=63-67 nuclei from three independent cultures. **(C, D)** Treatment with Nitarsone rescues Aβ_1-42_–induced synaptic loss. DIV16 hippocampal cultures were incubated with 5 µM Nitarsone, 500 nM Aβ_1-42_, 5 µM Nitarsone with 500 nM Aβ_1-42_ or vehicle control for 48h and stained for Shank3, Synaptophysin, and MAP2. (C) Representative confocal images of dendritic segments. Scale bar: 5 µm. (D) Number of synaptic puncta per 1 µm. N=33-38 dendritic segments from four independent cultures. **(E, F)** Treatment with Nitarsone rescues Aβ_1-42_–induced decrease of synaptic GluR1-immunoreactivity within Shank3. DIV16 dissociated, hippocampal cultures were treated with 5 µM Nitarsone, 500 nM Aβ_1-42_, 5 µM Nitarsone with 500 nM Aβ_1-42_ or vehicle control for 48h and stained for Shank3, surface GluR1, and MAP2. (E) Representative confocal images of dendritic segments. Scale bar: 5 µm. (F) GluR1-immunoreactivity within Shank3 signal. N=39-61 of dendritic segments from four independent cell cultures. **(G-K)** Nitarsone administration rescues mEPSC amplitude. (G) Analog traces of mEPSCs recorded in DIV16 hippocampal neurons treatd with 500 nM Aβ_1-42_, 5 µM Nitarsone, 5 µM Nitarsone with 500 nM Aβ_1-42_ or vehicle control for 48h. (H, J) Cumulative probability plots of (I) inter-event interval or (K) amplitude. Quantification of (I) inter-event-interval and (K) amplitude. N=24-28 neurons from four independent cell cultures. (A, E) Lookup table indicates pixel intensities from 0 to 255. ***p<0.001 by (B, D, F, I, K) two-way ANOVA followed by Tukey’s multiple comparisons test. All data are represented as mean ± SEM.

We therefore next assessed Aβ-induced synapse loss in dissociated hippocampal neurons that were kept for 48 h in the presence of 5 µM Nitarsone (Fig 6C, D). Aβ induced a 30% reduction of synaptophysin/Shank3 puncta in these cultures (Fig 6C, D) and this synapse loss was completely prevented by Nitarsone application (Fig 6C, D). The concomitant downscaling of synaptic surface expression of GluA1 AMPA-receptors was also significantly attenuated in the presence of Nitarsone (Fig 6E, F). In control experiments these neurons exhibited normal up-scaling of GluA1 AMPA-receptors in response to silencing of neuronal activity with 1 µM TTX application (Fig EV6D). Whole-cell patch-clamp experiments showed that reduced surface expression of GluA1 following Aβ treatment was accompanied by reduced miniature excitatory postsynaptic current (mEPSC) amplitude but not frequency (Fig 6G-K). Co-application of 5 µM Nitarsone preserved mEPSC amplitude while administration of the drug alone had no effect on both measures (Fig 6G-K).

### *In vivo* administration of Nitarsone prevents early synaptic dysfunction and cognitive deficits in two transgenic AD mouse lines

We next administered Nitarsone *in vivo* in two transgenic AD mouse lines with amyloid pathology, TBA2.1 and 5xFAD mice. 5xFAD mice express human APP and PSEN1 transgenes with a total of five AD-linked mutations (Oakley et al., 2006). These mice display less rapid spread of amyloid pathology than TBA2.1 mice, with visible plaques accompanied by gliosis at four months of age with accompanying synaptic dysfunction and cognitive impairment. We administered Nitarsone orally with forced feeding and a defined daily dose of 50 mg/kg that was based on a conservative NOEL (no observed effect level) from several toxicology studies and the rationale to achieve an effective dose in brain tissues (see methods and Fig 7A for experimental details).

**Fig. 7.**
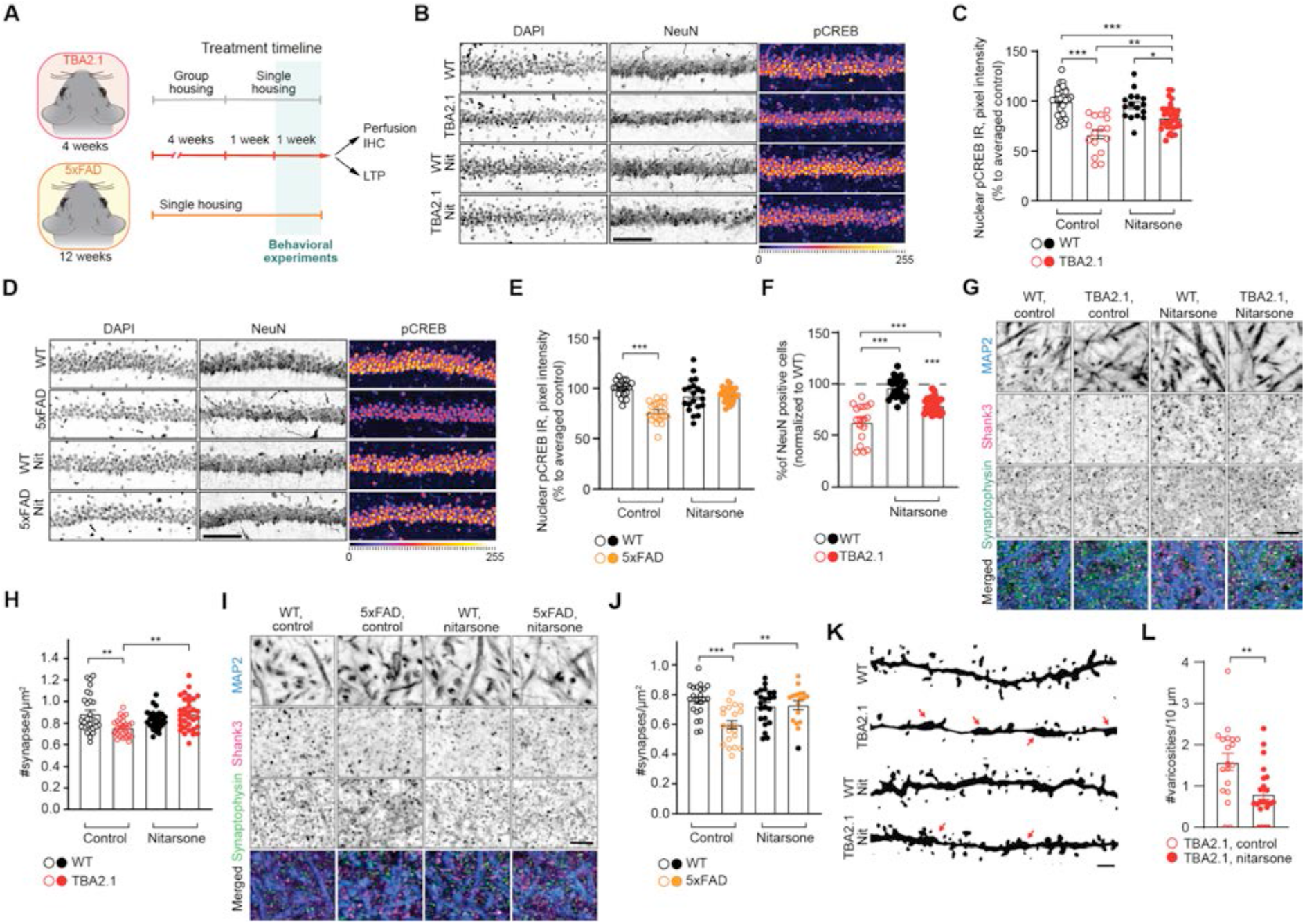
Oral administration of Nitarsone rescues CREB shutoff and synapse loss in TBA2.1 and 5xFAD mice. **(A)** Scheme representing the timeline of treatment with Nitarsone of TBA2.1 and 5xFAD mice. **(B, C)** Nitarsone rescues the reduction of pCREB immunoreactivity in NeuN positive cells in the CA1 region of TBA2.1 mice. (B) Representative confocal images of cryosections from 11 weeks old mice stained for NeuN, DAPI, and pCREB. Scale bar: 10 µm (C) Bar plot of pCREB nuclear staining intensity. N=21-34 hippocampal sections from 6-9 animals. **(D, E)** Nitarsone rescues the reduction of pCREB immunoreactivity in NeuN positive cells in the CA1 region of 5xFAD mice. (D) Representative confocal images of CA1 cryosections from 18 weeks old mice stained for NeuN, DAPI, and pCREB. Scale bar: 10 µm. (E) Cumulative frequency distribution of pCREB nuclear staining intensity. N=22-27 hippocampal sections from 6-7 animals. **(F)** Nitarsone reduces neuronal loss in TBA2.1 animals. The average number of NeuN-positive cells normalized to WT treated with vehicle. N=18-31 CA1 images analyzed from 6-9 animals per genotype. **(G, H)** Nitarsone prevents synapse loss in SLM of CA1 of TBA2.1 mice. (G) Representative confocal images of SLM from 11 weeks old mice stained for MAP2, Shank3, and Synaptophysin. Scale bar: 5 µm. (H) Number of synaptic puncta per ROI. N=27-33 ROIs from 6-9 animals **(I, J)** Nitarsone prevents synapse loss in SLM of CA1 of 5xFAD mice. (I) Representative confocal images of SLM from 18 weeks old mice stained for MAP2, Shank3, and Synaptophysin. Scale bar: 5 µm. (J) Number of synaptic puncta per ROI. N=17-22 ROIs from 6-7 animals. **(K, L)** Nitarsone reduces number of varicosities in SLM of CA1 of TBA2.1 mice. (K) Representative, confocal images of dendrites filled with biocytin stained from 13 weeks old mice. Arrows indicate varicosities. Scale bar: 1 µm. (L) Number of dendritic swellings per 10 µm. N=19-22 dendrites from 2 animals per genotype. *p<0.05, **p<0.01, ***p<0.001 by (c, e, f, h, j) two-way ANOVA followed by Bonferroni’s multiple comparisons test or (l) two-tailed Student t-test. All data are represented as mean ± SEM.

This regime had no effect on the body weight of treated as compared to control animals (Fig EV7A, B) and on amyloid load in both mouse lines (Fig EV7C-F). However, Nitarsone administration effectively prevented CREB shutoff in both transgenic AD mouse lines in the dorsal hippocampal CA1 region following 6 weeks of treatment (Fig 7B-E, Fig EV7G-J). In addition, early neuronal cell loss was reduced in TBA2.1 mice in comparison to vehicle-treated littermates (Fig 7F, EV7K, L) while we could not detect neuronal cell loss in CA1 of 5xFAD mice at 19 weeks of age (Fig EV7M, N). Most important, we found clearly reduced synapse loss in the stratum lacunosum moleculare in mice treated with Nitarsone in both animal models of amyloid pathology (Fig 7G-J). This is the first hippocampal region affected by amyloid-pathology in many AD animal models (Kerchner et al., 2012, Su et al., 2018). Synapse loss within this region was accompanied in TBA2.1 mice by dendritic varicosities (Fig 7K, L), another morphological feature of AD pathology (Jin et al., 2011, Lee et al., 2022). Mice treated with Nitarsone displayed significantly less dendritic varicosities (Fig 7K, L).

In accord with these findings we observed early synaptic dysfunction in acute hippocampal slices, as evidenced by deficits in late-phase long-term potentiation (LTP), in TBA2.1 and 5xFAD mice at postnatal week 11 and 19, respectively (Fig 8A-D). The reduced fEPSP slope in the last 30 minutes of recordings was rescued in slices of Nitarsone-fed mice in both AD lines (Fig 8A-D, EV7O, P), indicating a rescue of synaptic plasticity that is relevant for learning and memory (Fig 8A-D, Fig EV7O, P).

**Fig. 8.**
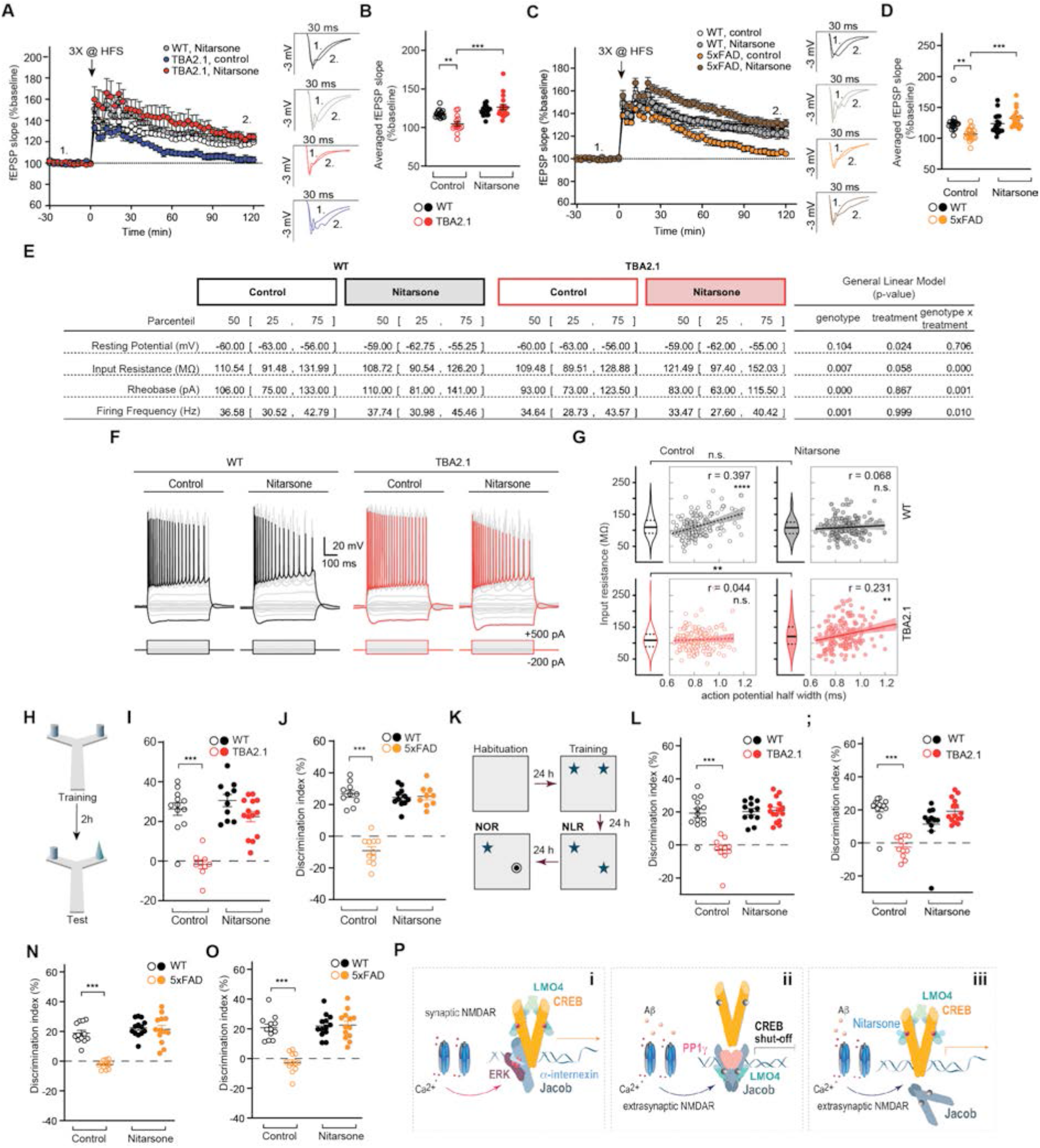
Oral administration of Nitarsone rescues AD-related phenotypes in TBA2.1 and 5xFAD mice. **(A)** Nitarsone rescues late CA1-LTP impairment in TBA2.1 mice. Insets show representative fEPSPs analog traces at indicated time points: 1 = baseline, 2 = late LTP. **(B)** Averaged fEPSP slopes recorded during the last 30 min. N=14-18 slices from 5-6 mice. **(C)** Nitarsone rescues late CA1-LTP impairment in 5xFAD mice. Insets show representative fEPSPs analog traces at indicated time points: 1 = baseline, 2 = late LTP. **(D)** Averaged fEPSP slopes recorded during the last 30 min. N=17-18 slices from 6 mice. **(E-G)** (E) Nitarsone treatment changed basic electrophysiological properties of CA1 pyramidal neurons in TBA2.1 mice. (F) Step current injection evoked responses of CA1 pyramidal cells from wild-type (WT) and TBA2.1 mice in control conditions and following Nitarsone-treatment. Grey and red traces display responses upon step current injections (see methods, black (WT) and red (TBA2.1) traces represent -200 and 500 pA current injection). (G) Nitarsone-treatment recovers the positive correlation of input resistance and action potential half width that is observed in control conditions. Consistent with a higher availability of CREB, CA1 pyramidal neurons showed a treatment-induced increase in input resistance in TBA2.1 mice (p=0.0036, post-hoc Dunn’s comparison). N=156-166 CA1 pyramidal neurons analyzed from 6 animals per genotype and treatment. **(H-J)** Nitarsone rescues short-term memory impairment in Y-maze object recognition task in (I) TBA2.1 N=9-14 and (J) 5xFAD mice. N=9-11 **(K-M)** Nitarsone rescues the impairment in novel location recognition in (K) TBA2.1 N=11-15 and (M) 5xFAD mice. N=12-13 **(N, O)** Nitarsone rescues the impairment in novel object recognition task in (N) TBA2.1 N=11-15 and (o) 5xFAD mice. N=12-13 **(P)** Jacob couples the origin of NMDAR signals to CREB activity. (i) Synaptic NMDAR signaling leads to nuclear translocation of a trimeric complex consisting of pJacob, pERK1/2, and α-internexin, which promote CREB phosphorylation. (ii) Activation of NMDARs at extrasynaptic sites by Aβ leads to nuclear import non-phosphorylated Jacob, displacement of LMO4 from CREB, binding to PP1, and subsequent CREB shutoff. (iii) Nitarsone specifically blocks Jacob-binding to LMO4 and thereby prevents CREB inactivation. **p<0.01, ***p<0.001 by (b, d, I, j, l-o) two-way ANOVA followed by Bonferroni’s multiple comparisons test or (g) Pearson correlation coefficient. All data are represented as mean ± SEM.

High CREB levels have been shown to increase excitability of neurons in the lateral amygdala and the hippocampus (Zhou et al., 2009, Yu et al., 2017). Therefore, we hypothesized that Nitarsone by preventing JaCS in TBA2.1 mice should increase neuronal excitability in hippocampal CA1 neurons. Using patch-clamp recordings we found a mild impact of genotype (transgene versus wild-type) on neuronal excitability in step current protocols (Fig 8F) probably reflecting the early stage of AD pathology in these mice. Nitarsone treatment during this early period increased the input resistance (Fig 8E, F), and reduced the rheobase (Fig 8E), and rescued the positive correlation of input resistance and action potential half width (Fig 8G) (Helmstaedter et al., 2009) that was observed in wild-type mice in control conditions (Fig 8E-G). The latter observation is particularly interesting because following Nitarsone treatment neurons with high input resistance and high half-width within the CA1 neuronal population could serve to specifically overcome deficits in synaptic transmission and the induction of synaptic plasticity (LTP).

We therefore next determined whether treatment with Nitarsone also rescues cognitive decline in TBA2.1 and 5xFAD mice (Oakley et al., 2006, Alexandru et al., 2011). To evaluate short-term memory, we used the Y-maze object recognition task (Creighton et al., 2019), which minimizes contextual cues, with an interval of 3h between training and test. Object recognition was impaired in TBA2.1 and 5xFAD mice treated with vehicle, when compared to littermate controls (Fig 8H-J). Conversely, transgenic TBA2.1 and 5xFAD mice fed with Nitarsone displayed improved discrimination performance in comparison to vehicle treated animals (Fig 8H-J). Human AD patients display impairments in object recognition, which essentially relies on proper synaptic function of CA1 neurons (Didic et al., 2013). Accordingly, TBA2.1 and 5xFAD mice showed deficits in novel object location and novel object recognition memory (Fig 8K-M, Fig EV7Q-T). Treatment with Nitarsone also rescued memory in a novel location recognition as well as in a novel object recognition task with a cognitive performance comparable to vehicle-treated littermate controls (Fig 8N, O). We did not observe major differences in distance travelled as well as the number of interactions with the objects (i.e. preference index) during the training (Fig EV7Q-T). Collectively, these data provide evidence that restoring synaptic plasticity and normal excitability with Nitarsone improve hippocampus-dependent learning and memory despite the presence of manifest amyloid pathology.

## DISCUSSION

Several lines of evidence suggest that the CA1 region, in both human patients and mouse AD models, is among the first to exhibit deficits in CREB activation, synaptic function and neuronal excitability (Wang & Bibb, 2011, Yiu et al., 2011, Kerchner et al., 2012, Su et al., 2018, Wirths & Zampar, 2020). CREB might be a nodal point in the AD transcriptome network, given the central role of this transcription factor in the regulation of gene expression that is essential for synaptic plasticity, intrinsic excitability and memory formation (Barco et al., 2002, Carlezon et al., 2005, Wang & Bibb, 2011, Alberini & Kandel, 2014, Teich et al., 2015). CRE-driven gene expression determines excitability of CA1 neurons (Barco et al., 2002, Lopez de Armentia et al., 2007, Alberini & Kandel, 2014), it reduces the threshold for LTP-induction in the Schaffer collateral pathway (Barco et al., 2002, Lopez de Armentia et al., 2007) and is instrumental for a switch from LTP to LTD that is characteristic for synaptic dysfunction in AD (Kim et al., 2001, Barco et al., 2002, Lopez de Armentia et al., 2007, Saura & Cardinaux, 2017). Despite this central role of CREB, research on amyloid pathology was largely focused on local signaling events that acutely elicit decay of synaptic function, largely ignoring the fact that molecular mechanisms underlying inactivation of CREB are likely to be first in the sequelae of events that cause synapse pathology. In consequence, such mechanisms remained elusive in AD.

Here we revealed a molecular mechanism implying Aβ-induced extrasynaptic NMDAR activation and nuclear import of Jacob for the induction of CREB shutoff (JaCS). Molecular modeling and screening for small chemical molecules subsequently led to the discovery that Nitarsone blocks binding of Jacob to the LIM1 domain. Application of Nitarsone *in vitro* and *in vivo* proved the relevance of the deciphered molecular mechanism for Aβ-induced synaptic pathology, CREB shutoff and the progression of synaptic and cognitive dysfunction at a very early stage of AD. Taken together, the data support the idea that macromolecular protein transport to the nucleus has a pathophysiological role in amyloid pathology. To our knowledge no other molecular mechanism for long-lasting transcriptional inactivation of CREB in neurons has been described yet and it is tempting to speculate that this mechanism will also contribute to early synaptic dysfunction elicited by similar mechanisms in other slowly progressing neurodegenerative diseases. Finally, a key finding of the present work is that the Jacob pathway is druggable and that the intervention with Nitarsone even holds promise for the treatment of synaptic dysfunction in humans.

### JaCS in AD

A key aspect in this regard is the balance between activation of extrasynaptic or synaptic NMDAR and the subsequent differential phosphorylation of S180 in Jacob. We provided evidence that Jacob directly associates with the bZIP domain of CREB irrespective of S180 phosphorylation and that this binding docks a signalosome to CREB that differs in its molecular composition depending upon the origin of NMDAR activation. We could show that binding of Jacob to LMO4 or, as shown previously, α-internexin (Karpova et al., 2013) determines whether the protein associates with the CREB phosphatase PP1γ or the MAP-kinase ERK1/2, and binding to either of these adaptors is decisive whether Jacob induces inactivation of CREB or enhanced CREB-dependent gene transcription (Karpova et al., 2013) / Fig 4). LMO4 is reportedly a transcriptional co-activator of CREB (Kashani et al., 2006) and our data suggest that LMO4 might hinder dephosphorylation of S133, stabilize the CREB dimer (Fig 5) and thereby could act as transcriptional enhancer. In the presence of amyloid pathology Jacob likely displaces LMO4 from the CREB complex (Fig 3N-Q), and we suppose that this contributes to long-lasting CREB dephosphorylation. Thus, enhanced binding of Jacob to PP1γ and displacement of LMO4, renders the association to LMO4 a key event for JaCS.

### JaCS contributes to early synaptic dysfunction in AD

Collectively the present study suggests that long-distance protein transport from extrasynaptic NMDAR to the nucleus is an important mechanism for disease progression at an early stage in AD. We hypothesize that this stage follows the initial hyperexcitability that has been described in transgenic mice with Aβ pathology and AD patients (Busche et al., 2012, Lam et al., 2017, Li & Selkoe, 2020). Recent work suggests that this hyperexcitability is at least in part caused by the suppression of glutamate reuptake (Zott et al., 2019), which in turn might cause sustained activation of extrasynaptic NMDAR in response to increased ambient glutamate levels. We propose that JaCS kicks in when extrasynaptic NMDAR activation is continuous and synergistically driven by ambient glutamate and oligomeric Aβ (Rönicke et al., 2011). In addition, we propose that nuclear import of Jacob might be the initial trigger for decay of synaptic function that is induced by altered gene transcription. Along these lines we found in previous work that the earliest morphological phenotype following nuclear accumulation of non-phosphorylated Jacob is the stripping of synaptic contacts (Dieterich et al., 2008, Karpova et al., 2013). Our observation of a recovery of a positive correlation between input resistance and action potential half width following Nitarsone treatment in TBA2.1 mice (Fig 8G) is particularly interesting with respect to the rescue of LTP. Both parameters lead to a more effective input to output conversion of pyramidal neurons by increasing depolarization upon (synaptic) input and a more efficient transmitter release, respectively. The observed effects compensate for a reduction in synaptic strength and will facilitate the induction of synaptic plasticity. It is likely, that this effect strongly contributes to Nitarsone-mediated rescue of synaptic plasticity observed in this study.

### Preserving synaptic function in AD by targeting the interaction of Jacob to LMO4

We could prove the relevance of CREB and of the proposed mechanism for Aβ-induced decay of synaptic function with a small chemical compound that was selected based on structural modeling of the most crucial binding interface that is involved in CREB shutoff. Thus, Nitarsone selectively interrupts the interaction of Jacob, but not of CREB with the LIM1 domain of LMO4 and competes with a 15 amino acid short peptide in Jacob that binds to LIM1. Moreover, we identified two peptides within the LMO4 binding region of CREB. Structural modeling predicts that CREB binds with both regions to a LIM domain tandem of LMO4 with two-times higher binding energy than Jacob (Fig EV5A-D). The second LIM2 domain has a weaker binding site for the CREB peptides than LIM1 as already shown for other proteins, CtIP and Lbd1 (Deane et al., 2004, Stokes et al., 2013). In concordance, we were not able to show binding of CREB to the isolated LIM2 domain, however, binding to the LIM1 of the first peptide might induce and facilitate binding of the second to the LIM2 domain. Of note, according to our model CREB-binding will not interfere with self-association of the LIM2 domain of LMO4 that has been reported previously (Deane et al., 2004).

### The therapeutic potential of Nitarsone

The most relevant therapeutic window in AD is right at the beginning of synaptic dysfunction, and in light of these findings Jacob is an attractive target for interventions to rescue or even restore synaptic plasticity in the early symptomatic phase. Thus, the intervention with Nitarsone might open up effective and selective therapeutic avenues that directly target altered NMDAR-to-nucleus communication without affecting NMDAR function at the plasma membrane, which has several detrimental side effects.

Nitarsone has been in use in poultry farming as feed additive to prevent histomoniasis and to improve food utilization until 2015, when the compound was withdrawn from the market as a precaution by the FDA (FDA, 2015). The identified health risk, however, was very low since it was estimated that life-long consumption of turkey meat might result in increased lifetime risk of developing or dying from cancer of 0.00031% (Nachman Keeve et al., 2017). Nitarsone is the oxidized form of arsanilic acid, an organic arsenic compound, considered to be less harmful than inorganic arsenic like arsenic trioxide (ATO) (Fowler et al., 2022). However, a transformation in carcinogenic arsenic might be possible (Nachman et al., 2017) by anaerobic gut microbiota (Chen & Rosen, 2016) and chemical modification of the compound to prevent this transformation might be difficult given its almost perfect fit to the binding pocket. On the other hand, arsenic itself has a long tradition in folk and veterinary medicine. It was used for many years to treat syphilis and other disease states (Iland & Seymour, 2013) and even though human arsenic methyltransferases in the liver convert ATO to cytotoxic arsenic (Maimaitiyiming et al., 2020), ATO has become the standard treatment of acute promyelocytic leukemia (Lo-Coco et al., 2013, de Almeida et al., 2021).

In light of these arguments and given that AD is a lethal neurodegenerative disease, one might consider Nitarsone a therapeutic option to attenuate early synaptic dysfunction and cognitive decline and thereby to slow down disease progression. A clear limitation of the present study is, however, that interventions that have beneficial effects in transgenic AD mouse models might not be suitable for several reasons in humans. As a matter of fact, AD mouse models have limitations and do not resemble all aspects of the human disease. Along these lines despite strong evidence that Aβ-driven signaling underlies the pathological mechanisms of AD, in recent years doubts have been casted whether other pathways might contribute to the disease progression and cognitive decline (Haass & Selkoe, 2022). The failure of clinical trials with anti-amyloid therapies to show robust beneficial outcomes in AD patients on cognition has raised concerns that lowering amyloid load is sufficient to stop disease progression (Haass & Selkoe, 2022). We speculate that in light of the complexity of mechanisms in the so-called ‘cellular phase’ of AD (Haass & Selkoe, 2022) and the notion that dissolving β-amyloid plaques with antibodies might have detrimental side effects due to release of oligomeric Aβ species, a combinatorial intervention with anti-Aβ antibodies (like Aducanumab) and Nitarsone might be a more effective means to slow down early synaptic failure in AD. This hypothesis can be tested first in future work in transgenic AD mouse models.

## MATERIALS AND METHODS

### Human subjects

The temporal cortex (area 22) biospecimens (Table S1) were provided by the Brain Banking Centre Leipzig of the German Brain-Net, operated by the Paul Flechsig Institute of Brain Research (Leipzig University). The diagnosis and staging of Alzheimer’s disease cases was based on the presence of neurofibrillary tangles(Braak & Braak, 1991) and neuritic plaques in the hippocampal formation and neocortical areas as outlined by the Consortium to establish a registry for Alzheimer’s disease (CERAD; (Mirra et al., 1991)) and met the criteria of the National Institute on Aging on the likelihood of dementia (The National Institute on Aging, 1997). For the study of human biospecimens informed consent forms were obtained and autopsies were approved (GZ 01GI9999-01GI0299).

### Animals

Animals were maintained in the animal facility of the Leibniz Institute for Neurobiology, Magdeburg. Jacob/Nsmf knockout (homozygous labelled as −/−) animals were characterized previously (Spilker et al., 2016). TBA2.1 (homozygous labelled as TBA2.1) mice (Alexandru et al., 2011) were a kind gift from ProBiodrug. To generate double transgenic animals (animals homozygous for both mutations labelled as TBA2.1, −/−) Jacob/*Nsmf* heterozygous mice were crossed with heterozygous TBA2.1 mice. The 5xFAD mice (Oakley et al., 2006) were purchased from Jackson Laboratories. All lines had a C57BL/6J background. Animal experiments were carried out in accordance with the European Communities Council Directive (2010/63/EU) and approved by local authorities of Sachsen-Anhalt/Germany (reference number 42502-2-1501 LIN, 203.m-42502-2-1112LIN, 203.m-42502-2-1280LIN). Mice were house din individually ventilated cages (Green line system, Tecniplast) under controlled environmental conditions (22 ± 2°C, 55% ± 10% humidity, 12 h, light–dark cycle, with lights on at 06:00 a.m.), with free access to food and water. Unless indicated otherwise, the animals were housed in groups up to 5 mice per cage. All animals were genotyped prior and after the experiment. The exact information concerning sex, age, and treatments are described in method details.

### Murine organotypic hippocampal slice culture (OHSC)

OHSC were prepared according to previously published work (Grochowska et al., 2017). Slices were obtained from P7-P9 mice of both sexes (Jacob/*Nsmf* knockout or WT littermates as a control). Animals were decapitated, brains removed, and hippocampi dissected under a binocular. 350-400 µm thick Perpendicular slices were cut using a McIlwain tissue chopper (Mickle Laboratory Engineering). Slices were cultured on millicell membranes (3 slices per membrane, Merck Milipore) in 6 well-plates in 1 ml of medium 50% minimal essential medium (Gibco), 25% heat inactivated horse serum (Gibco), 25 mM glucose, 2 mM glutamine, 25 mM HEPES, 1xB27 (Gibco), penicillin/streptomycin (100 U/ml). Cultures were grown at the 37°C, 5% CO2, 95% humidity. Every 3rd day the 700 µl of the medium was exchanged.

### Primary hippocampal cultures

Hippocampal and cortical cultures were prepared from Wistar rat embryos (E18) of mixed sex as described previously (Spilker et al., 2016). Briefly, dissected were digested for 15 min with trypsin at 37°C. Neurons were plated on plastic 12-well dishes (Greiner) on glass coverslips coated with poly-L-lysine (Sigma-Aldrich) at a density of 60 000 cells/well in DMEM medium (Gibco, Thermo Fisher Scientific) supplemented with 10% FCS, 1x penicillin/streptomycin, and 2 mM glutamine. After 1h incubation (at the 37°C, 5% CO2, 95% humidity) cells were kept in BrainPhys medium supplemented with 1%SM1 (Stemmcell Technologies), 0.5 mM Glutamine (Gibco) at 37°C, 5% CO2 and 95% humidity.

### Cell lines

HEK293T cells were maintained DMEM medium (Gibco, Thermo Fisher Scientific) supplemented with 10% FCS, 1x penicillin/streptomycin, and 2 mM glutamine at the 37°C, 5% CO2, 95% humidity.

### Antibodies, chemicals, recombinant proteins, software

The antibodies, chemicals, kits, recombinant proteins, peptides, and software can be found in Table S2.

### Structural Modelling

Structures of LMO4:peptide complexes were modeled using coordinates of LMO4:Ldb1 complex (PDBId: 1RUT) using Swiss-PDB Viewer v4.1 (Guex & Peitsch, 1997, Johansson et al., 2012). Positional refinement and calculation of free binding energy ΔΔG of LMO4:peptide complexes were performed by FoldX v5 (Schymkowitz et al., 2005). Donor and acceptor atoms of LMO4 LI1:Jacob peptide complexes were identified using ZINCPharmer and ZINC15 (11/20) (Koes & Camacho, 2012). Nitarsone (4-Nitrophenylarsonic acid) has been geometrically optimized and generated in PDB format by Avogadro v1.2 (Hanwell et al., 2012) and used as ligand for LMO4 LIM1 by molecular docking program AutoDock Vina v1.1.2 (Eberhardt et al., 2021). Secondary structures of full-length hJacob and hCreb were predicted with PsiPred v1.1.2 (McGuffin et al., 2000). RaptorX (12/20) (Källberg et al., 2012) was used to predict the structure of Jacob C-terminus. Structures were visualized using Open-source PyMol v2.5 (pymol.org).

### Yeast-Two-Hybrid screening

The Yeast-Two-Hybrid screening for Jacob interaction partners was performed using a pretransformed human brain cDNA Library in pACT2 (Matchmaker-GAL4 Two-Hybrid II; Clontech) as described previously (Helmuth et al., 2001; Dieterich et al., 2008). The yeast strains AH109 and Y187 as a partner (both from Clontech) were used.

Jacob-LMO4 interaction was reconfirmed using fusion vectors (bait vector pGBKT7, prey vector pGADT7) using MATCHMAKER Two-Hybrid System 3 (Takara Bio Europe/Clontech, France). Co-transformed yeasts were assayed for growth on quadruple drop-out medium (SD/–Ade/–His/–Leu/–Trp) and additionally for LacZ reporter activation accordingly to the manufacturer’s instructions.

### Recombinant protein production

GST-tagged recombinant proteins (LMO4) were produced and purified as described previously (Dieterich et al., 2008, Karpova et al., 2013). For protein production E. coli BL21 (DE3) strain was used. Following induction with with 0.3 mM isopropyl-beta-D-thiogalactoside (IPTG) at 18°C cells were pelleted by centrifugation at 6.000 x g for 15 min and purified from the soluble fraction by glutathione-Sepharose <u>chromatography</u> (elution buffer: 50 mM Tris-HCl, 10 mM reduced glutathione, pH 8.0). For MBP-tagged recombinant proteins (Jacob fragments (in aa): 45-228, 1-116, 117-228, 262-532, 262-532, 45-228, 45-228-S180D (phosphomimetic mutant), 45-228-S180A (phosphodeficient mutant)) lysis was done in 20 mM Tris buffer (pH 7.4), 200 mM NaCl, 1 mM DTT and 1 mM EDTA. Protein was eluted with lysis buffer+10 mM maltose. His-SUMO-tagged recombinant proteins (CREB, CREB fragments (in aa): 1-88, 1-165, 1-293, 89-314, 89-165, 166-341, 166-293, 294-341) were purified from native conditions in which lysis buffer, wash, and elution buffer all have 50 mM NaH2PO4 (pH 8.0) and 300 mM NaCl with various concentrations of imidazole (10, 20 and 250 mM respectively). All Protease inhibitor (Complete, Roche) was used in all mentioned buffers. MBP-tagged PP1ɣ was obtained from Creative BioMart (Cat. #PPP1CC-84). The purity of the protein was checked on SDS-PAGE gels stained and Coomassie blue staining.

### Isothermal titration calorimetry (ITC)

LMO4-Nitarsone (98%, ABCR Gute Chemie) binding affinity was measured using VP-ITC calorimeter (MicroCal) and data were analysed by MicroCal LLC ITC software (MicroCal). Both purified GST-LMO4 protein and Nitarsone were prepared in 25 mM Tris buffer (pH 7.4) containing 50 mM NaCl. 20 µM GST-LMO4 or buffer control (to calculate heat of dilution) was titrated against 180 µM Nitarsone. LMO4 and Jacob binding affinity was measured by analysing binding isotherms for titration of 5 µM Jacob and 40 µM LMO4. To estimate the influence of Nitarsone on Jacob-LMO4 interaction, LMO4 protein was saturated with Nitarsone, and subsequently the Jacob protein was injected. Typically for an ITC experiment 30 injections of 10 µl each were made at 180 sec intervals. Heat change was determined by integration of the obtained peak of differential power by the instrument. Different parameters like binding enthalpy (ΔH), dissociation constant (Kd), and stoichiometry were calculated.

### Pull-down assays

Pull-down assays performed as described previously (Dieterich et al., 2008, Karpova et al., 2013). Briefly, protein amounts ranging from 1-10 µg along with the equivalent amount of the control tag protein was bound on the respective beads and was incubated with 5% BSA for 1 h at room temperature (RT). 200 ng-10 µg of the second recombinant protein was incubated with the resin bound protein in 1ml TBS buffer for 1h either at room temperature or 3 h at 4°C. After three washing steps with TBS buffer containing 0.2-0.5 % Triton X100, the complex was eluted in 2X SDS sample buffer (250 mM Tris-HCl, pH 6.8, 1% (w/v) SDS, 40% (v/v) Glycerol 20% (v/v) β-mercaptoethanol, 0,004 % Bromophenol Blue).

For competition pull-down assay 20µg of GST-LMO4 and equimolar amounts of GST control protein were bound to glutathione beads followed by 5% BSA blocking and washing with 50 mM Tris-Cl, pH 7.5, 150 mM NaCl. Different combinations of recombinant proteins (i.e., HIS-SUMO-CREB; 1:1 of His-SUMO-CREB and MBP-Jacob-45-228; 1:4 of His-SUMO-CREB and MBP Jacob-45-228; and 1:8 of His-SUMO-CREB and MBP-Jacob-45-228) were incubated with GST-LMO4 and GST (control protein) immobilized on Protino Glutathione Agarose 4B beads (Macherey-Nagel). Probes were eluted with 25µl of 2x SDS sample buffer after incubation and washing steps (washing buffer: 50 mM Tris pH 7.4, 500 mM NaCl, 0.1 % Triton X100 and protease inhibitor without EDTA).

For pull-down assays with Nitarsone 10-20 µg of GST-LMO4 or GST control protein were immobilised on glutathione beads followed by overnight incubation with 1-20 µM of Nitarsone solution at 4°C. Equimolar MBP-Jacob-45-228 were added to the solution and incubated rotating for 2 h and washing was performed with 20 mM Tris-Cl (pH 7.4), 150 mM NaCl, 0.2 % TritonX-100 containing phosphatase and protease inhibitors. Complex was eluted using 2X SDS sample buffer.

The exact scheme of the pull-down experiments is included in the main figures.

### Heterologous co-immunoprecipitation

The constructs (LMO fragments (in aa) tagged with tRFP: 1-80 or 81- with GFP, or Jacob fragments (in aa): tagged with GFP: 45-172, 173, 246; mcherry-PP1ɣ (Liu et al., 2010) with GFP or Jacob or Jacob fragments (in aa) tagged with GFP: 1-172, 247-309, 310-532, 213-246, 173-246) were heterologously expressed in HEK293T cells. Cells were harvested in cold TBS buffer containing protease inhibitors (Roche) and phosphatase inhibitors (Roche), in a 1g/10ml ratio 48 h after transfection. The pelleted cells were lysed in cold RIPA buffer (50 mM Tris-HCl (pH 8.0), 150 mM NaCl, 1% NP40, 0,5% doxycycline (DOC), 0,1%, sodium dodecyl sulfate, (SDS), protease and phosphatase inhibitors, pH 7.4) for 1,5 h in 4°C, rotating. The lysates were subsequently centrifuged at 20 000 g for 20 min. The supernatant was incubated with 25 µl of µMACS Anti-GFP MicroBeads (Miltenyi Biotec) for 40 min in 4°C, rotating. Beads were collected with the use of µMACS magnetic column and washed twice with 400 µl of RIPA buffer and 300 µl of 20 mM Tris-HCl (pH 7.5). 60 µl of hot 2x sample buffer (250 mM Tris-HCl, pH 6.8, 1% (w/v) SDS, 40% (v/v) Glycerol 20% (v/v) bmercaptoethanol, 0,004 % Bromophenol Blue) was used for protein elution.

The phospho-dependent association of overexpressed nuclear Jacob was assessed by heterologous co-immunoprecipitation from HEK293T cells pre-treated with 20 nM staurosporine (Cell Signaling Technology, Cat.: #9953S) for 3 h.

### Characterization of Jacob antibodies

For initial characterization of anti-panJacob and pJacob (S180) antibodies for the detection of human protein HEK293T cells were transfected with the plasmid expressing hJacob-SPOT. Overexpressed protein was immunoprecipitated from the cell lysate using tag specific nano-trap (SPOT-Trap-MA, Chromotek) and analysed by WB.

### Immunoblotting of brain protein extracts

Human brain samples were homogenized in a buffer containing 0.32 M sucrose, 5 mM HEPES, protease inhibitors (Sigma) and phosphatase inhibitors (Roche), in a 1g/10ml ratio. The homogenates were centrifuged at 1000g for 10 min at 4°C. The pellets were used for immunoblotting. For immunoblotting of murine samples (CA1), dissected hippocampi were homogenized with TRIS buffered saline (25 mM Tris-HCl, 150mM NaCl, pH 7.4 buffer in presence of protease (Complete, Roche) and phosphatase inhibitors (PhosSTOP, Roche). Band intensities were quantified with Fiji/ImageJ (Schindelin et al., 2012).

### Quantitative real-time PCR (qPCR)

Hippocampal homogenization, RNA extraction and cDNA preparation were described previously (Spilker et al., 2016). *Bdnf exon IV* (F primer: 5′- GCAGCTGCCTTGATGTTTAC-3’, R primer: 5′-CCGTGGACGTTTGCTTCTTTC-3′) and *β-actin* (reference gene, F primer: 5′-AACGCAGCTCAGTAACAGTCC-3’, R primer: 5′-GTACCACCATGTACCCAGGC-3’) cDNA were amplified using the iScript RT-PCR iQ SYBR Green Supermix (BIORAD, Cat.:# 170882) in a qPCR detection system (LightCycler LC480, Roche). The sequences are indicated in Table S2. The relative expression levels were analysed using the 2-ΔΔCt method with normalization relative to *β-actin*.

### Cell based co-recruitment assay

HEK293T cells were transfected with the following constructs: Jacob-1-228 tagged with GFP with tRFP or LMO4 or LMO4 fragments (in aa): 1-165, 1-80, 81-165; PP1ɣ tagged with GFP(Trinkle-Mulcahy et al., 2001) and mCherry(Liu et al., 2010) or ΔMyr-Jacob tagged with tRFP. On the following day cells were fixed with 4% PFA, permeabilized with 0,1% TX-100 in 1xPBS for 10 min, stained with DAPI and mounted. Z-stack with 300 nm step size was taken with 512×512 pixels format using a Leica TCS SP8-STED system. Maximal intensity images from three optical sections were generated for representative images.

### FRET and SRET assays

For FRET experiments, HEK293T cells were transiently co-transfected with MaxPEI, (Polysciences, Cat.: #23966) with different combinations of two constructs of interest tagged with donor (GFP: LMO4, CREB, Jacob, Jacob fragments (in aa): 1-228, 262-532) or acceptor (tagRFP: LMO4, LMO4 fragments (in aa): 1-80, 81-165, CREB, CREB fragments (in aa): 1-165, 166-341), with constant concentration of cDNA of donor and increasing concentration of acceptor. FRET was performed as described previously (Carriba et al., 2008). For SRET measurements, transfected cells were resuspended in Hank’s balanced salt solution (HBSS) supplemented with 10 mM glucose (Sigma-Aldrich). Triple-transfected (CREB-Rluc:Jacob-GFP:LMO4-tagRFP) cell suspension was used to perform the three measurements. (i) The LMO4-tagRFP expression level was assessed by the tagRFP fluorescence intensity. (ii) The CREB-Rluc expression level was estimated by CREB-Rluc luminescence determined 10 min after addition of coelenterazine-H (5 µM). (iii) For SRET, the cells were incubated with 5 µM of DeepBlueC (Molecular Probes). The measurements were done with Mithras LB940 (Berthold Technologies) equipped with detection filters at 400 nm and 590 nm. Net SRET was defined as [(long-wavelength emission)/(short-wavelength emission)]-Cf, where Cf corresponds to [(long-wavelength emission)/(short-wavelength emission)] for cells expressing CREB-Rluc:Jacob-GFP:LMO4-tRFP.

### Luciferase assay

HEK293T cell line stably expressing luciferase under the CRE promoter (vector pGL4.29[luc2P/CRE/Hygro], Promega) were transfected for 24h and lysed with the lysis buffer (25 mM Tris-phosphate, 2 mM DTT, 2 mM 1,2-diaminocyclohexane-N,N,Ń,Ń-tetraacetic acid, 10% glycerol, 1% TX-100, pH 7.8). The luciferase activity was measured with the Dual-Luciferase Reporter Assay System (Promega) on a (FLUOstar Omega, BMG Labtech).

### Animal perfusion and immunohistochemistry

Animals were anesthetized with ketamine/xylazine (Medistar) and transcardially perfused with 0.9% saline followed by 4% PFA. The brains were post-fixed in 4% PFA in PBS overnight, followed by immersion in 0.5 M sucrose for 24h and then 1M sucrose for another 1 to 2 days or until the brains were sunken down. The brains were snap-frozen and cut into 35 µm thick coronal cryosections (Leica CM3050S, Leica-Microsystems). The sections were blocked in a buffer (0.3% TX-100, 10% NGS in PBS) for 30 min at RT, and incubated with primary antibodies diluted in the blocking solution, overnight at 4°C. Secondary antibodies were applied for 2h at RT. Sections were counterstained with DAPI and mounted in Mowiol 4-88 (Merck Chemicals). For the detection of Aβ, the heat-based antigen retrieval method was used. For anti-Aβ staining, following permeabilization, brain sections were immersed into 10 µM sodium citrate solution (Fluka, pH=9) for 30 min at 80°C. Imaging of nuclear pCREB/CREB immunoreactivities in cryosections was performed using a Leica TCS SP5 system (Leica-Microsystems). Images were acquired sequentially with a HCX ApoL20x/1.0 water objective, optical zoom 4. Sections were imaged with constant laser/detector settings along the z-axis with 400 nm step in a 512×512 pixel format.

### OHSC stimulation and immunocytochemistry

For Aβ-induced CREB shutoff Aβ_1-42_ oligomeric solution was prepared as described in Grochowska et al., (2017). Briefly, the Aβ_1-42_ peptide film (Anaspec) was re-suspended in 2 µl DMSO (Sigma-Aldrich), sonicated for 5 min, diluted with F12 medium (Gibco) to final concentration of 50 µM, sonicated for 10 min, and left for oligomerization at 4°C, on. 7 DIV OHSC were treated with 1 µM of Aβ_1-42_ oligomers for 1h, fixed for 1h at RT in 4% PFA/4% sucrose. After fixation the slices were washed, permeabilized with 0.4% TX-100 and treated with 50 mM NH_4_Cl for 30min. Subsequently, the samples were blocked for 1h at RT in 10% normal goat serum (NGS) in PBS. Next, the slices were incubated for 72h with primary antibody followed by incubation with the secondary antibody for 24h. Following counterstaining with 4’,6-diamidino-2-phenylindole (DAPI; Vectashield/Biozol) slices were mounted in Mowiol 4-88 (Merck Chemicals).

### Primary cultures transfection and stimulation

Hippocampal neuronal cultures were transfected with plasmid cDNA using Lipofectamine2000 (Thermo Fisher Scientific) following the manufacturer’s instructions. For LMO4 knockdown cells were expressing shRNA (5’- AGATCGGTTTCACTACATC -3’) for 5 days; for Jacob knockdown 4 days. For CREB shutoff experiments hippocampal neurons were transfected at DIV15 with NLS or ΔMyr-Jacob or ΔMyr-Jacob-L175A-V176A tagged with GFP or GFP and fixed after 24h. For the experiment with okadaic acid (OA, Tocris) cells were treated for 20 min with 2 µM OA 1 day post transfection. The target sequences for LMO4 and NLS-Jacob knockdown (Spilker et al., 2016) as well as scrambled controls are indicated in Table S2. For Aβ-induced CREB shutoff experiments Aβ oligomers (Anaspec) were prepared as described previously (see previous section and (Grochowska et al., 2017)). Transfected neurons were treated with 500 nM Aβ_1-42_ or Aβ_3(pE)-42_. Nitarsone was diluted in distilled water. The concentration and duration of treatments are indicated in the figure legends.

### Immunostaining of primary neurons

Primary neuronal cultures were fixed with 4% PFA/4% sucrose solution for 10 min at RT, washed with PBS, and permeabilized with 0.2% TX-100 in PBS for 10 min. Then, cells were incubated for 1 h in blocking solution (2% glycine, 0.2% gelatin, 2% BSA, and 50 mM NH_4_Cl (pH 7.4)) and primary antibodies were applied overnight at 4°C. Next, the coverslips were incubated with secondary antibodies diluted 1:500. Coverslips were mounted with Mowiol 4-88 (Merck Chemicals). For detection of Jacob-CREB and Jacob-LMO4 co-localization in STED imaging, a heat-based antigen retrieval protocol was used.

For surface expression of AMPA receptors dissociated hippocampal neurons were incubated for 10 min at RT with anti-GluA1 antibody diluted in Tyrodés buffer (128 mM NaCl, 5 mM KCl, 1 mM MgCl2, 2 mM CaCl2, 4.2 mM NaHCo3, 15 mM HEPES, 20 mM glucose, pH 7.2-7.4), rinsed, and fixed for 10 min at RT with 4% PFA-sucrose, and subsequently stained with other antibodies as described above.

### STED microscopy

STED images were obtained using a Leica TCS SP8 STED 3X equipped with pulsed White Light Laser (WLL) and diode 405 nm for excitation and pulsed depletion laser 775 nm and Leica HC ApoCS2 100x/1.40 oil objective. MAP2 and DAPI were acquired in confocal mode. The pixel size for all images was 20-23 nm (XY-plane). The Z-step was 150 nm. STED and confocal images were deconvolved using Deconvolution wizard (Huygens Professional, SVI), with optimized iteration method. Intensity profiles were created in Fiji/ImageJ.

### Neuronal nuclei isolation and flow cytometry

Nuclei were isolated with a Nuclei Isolation Kit (Sigma Aldrich, Cat.: #NUC101) according to the manufacturer’s protocol. Nuclei were fixed with ice-cold methanol for 10 min. 0.5% Triton for permeabilization and 10% normal donkey’s serum for blocking were used followed by 1h RT incubation with respective primary and, subsequently, secondary antibodies (see Table S2). Nuclei were counterstained with Hoechst33342 (1:500, ThermoFisher Scientific, Cat.:# H3570). Samples were measured with a BD LSR II flow cytometer (BD Biosciences) and analysed with FlowJo (LLC).

### Whole-cell patch-clamp recording in slices

Hippocampal coronal brain slices (300 µm) were obtained from TBA2.1 mice, as described before(Justus et al., 2017) in four groups of mice: wild type littermates and TBA2.1 mice treated with sucrose; wild type littermates and TBA2.1 mice following Nitarsone treatment. Slices were kept at room temperature for up to 8 hours in ACSF solution of the following composition in: 125 mM NaCl, 3 mM KCl, 1.25 mM NaH2PO4, 26 mM NaHCO3, 2.6 mM CaCl2, 1.3 mM MgCl2, 15 mM glucose (supplied with 95% O_2_, 5% CO_2_).

Whole cell patch-clamp recordings were performed in the CA1 area of the hippocampus in ACSF solution using borosilicate glass (GB150F-8P, Science Products) with pipette resistance in the range 3-6 MΩ. The internal solution contained: 140 Mm K-gluconate, 7 Mm KCl, 5 Mm HEPES-acid, 0.5 Mm MgCl2, 5 Mm phosphocreatine, 0.16 Mm EGTA and was implemented with Biocytin (Sigma-Aldrich). The signals were acquired with either an BVC-700A amplifier (setup 1, Dagan) or an ELC-03XS (setup 2, NPI) amplifier at 50kHz via a LIH 8+8 interface (HEKA) and Igor Pro software (Version 8.0.4.2, WaveMetrics). The mean series resistance was 21.3 ± 0.2 MΩ (mean ± SEM). Recordings were performed at the temperature of 33°C. Membrane potential was not corrected for the liquid junction potential. No difference in membrane potential was observed between the experimental groups. For the investigation of the electrophysiological properties neurons were held at a membrane potential of -65 mV by current injection. First, a step current injection protocol was performed, consisting of 500 ms step pulses with of −200 pA, −100 pA, −50pA, −30 pA, −20 pA, −10 pA, +10 pA, +20 pA, +30 pA, +50 pA, +100 pA, +200 pA, +300 pA, +400 pA and +500 pA. For estimation of rheobase 500 ms square pulse currents were injected with increasing amplitudes in steps of 2 pA until an action potential was generated.

The input resistance (IR) was calculated using -30 pA to +30 pA step-wise current injections as the slope of the relationship of the injected current and the steady state voltage response(Justus et al., 2017). Recordings were analyzed using Python (Version 5.1.5) and Igor Pro (V. 8.0.4.2, WaveMetrics). Statistical analysis was conducted using Prism 8 for Windows (Prism version for Windows, GraphPad Software) and IBM SPSS Statistic (Version 21).

### SPECT-imaging of cerebral blood flow

Mice (three months of age, both sexes) were intravenously injected with 99mTcHMPAO via chronically implanted catheters in the right external jugular vein. Jugular vein catheter implantation and preparation of the 99mTcHMPAO injection solution were done as described in Kolodziej et al. (2014). For tracer injection, the catheter was extended by a polyethylene (PE) tube (BioMedical Instruments, 60 cm, prefilled with 150 µl 0.9%NaCl) and connected to a syringe containing 130-170MBq/330 µl of freshly prepared 99mTc-HMPAO-solution. Injections were made at flow rates of 33 µl/min during periods of 15 min. During this time, the animals were awake and freely moving. After injection, animals were anaesthetized (4-1.5% isoflurane, 800 ml/min O2) and transferred to the SPECT/CT-scanner. The injected 99mTc activity was calculated by determining the amounts of 99mTc that had remained in the syringe and in the extension tube by using a radionuclide calibrator (Aktivimeter Isomed 2010, Nuklear-Medizin-Technik Dresden GmbH). Co-registered head SPECT/CT-scans were made using a four-head NanoSPECT/CT (Mediso). CT scans were made at 45kVp, 177µA, with 180 projections, 500 ms per projection and 96 µm spatial resolutions, reconstructed (InVivoScope 1.43) at isotropic voxel-sizes of 100 µm. SPECT scans were made using ten-pinhole mouse brain apertures with 1.0 mm pinhole diameters, providing a nominal resolution of <0.7 mm. Twenty-four projections were acquired during a scan time of one hour. Axial FOV was 20.9 mm. Photopeaks were set to the default values for 99mTc (140keV +/− 5%). SPECT images were reconstructed (HiSPECT, v 1.4.1876, SCIVIS) at an isotropic voxel output size of 250 µm. The co-registered SPECT/CT-images were aligned to a reference MR (MPI Tool software, v6.36, ATV; Dorr-Steadman-Ullmann-Richards-Qiu-Egan (40 micron, DSURQE)) and SPECT brain data-sets were global mean normalized using ImageJ. Further data processing and calculation of group means and differences and statistical analyses were done using ImageJ and Matlab (R2017b). Results were illustrated with Osirix (v. 5.8.1) (Rosset et al., 2004).

### Nitarsone feeding regime

Nitarsone (98%, ABCR Gute Chemie) was prepared in aliquots of 15 mg/850 µl (350 µl dH_2_O, 500 µl MediGel Sucralose, ClearH_2_O) to be used within five days. Mice were treated with Nitarsone (50 mg/kg body weight) or vehicle (sucrose solution) once a day per os for a total of 6 to 7 weeks. For TBA2.1 mice (both sexes), treatment started at the age of 4 weeks. The animals were group housed and force fed for 4 weeks. Afterwards they were single housed and allowed to choose voluntary feeding (Nitarsone or vehicle mixed into a gel paste, DietGel boost Hazelnut, ClearH2O) over continued force-feeding. Treatment in 5xFAD male mice started at the age of 12 weeks. Here, animals were single housed right away and given a choice between voluntary and forced feeding. Behavioral experiments were conducted for both lines during the 6^th^ week of Nitarsone treatment. During the course of the 7^th^ week of treatment, animals were either subjected to perfusion to obtain their brains for subsequent immunohistochemistry or they were culled to obtained native brains for electrophysiological recordings.

### Open-field test

Locomotor activity was assessed using the Open-field test, performed in a square arena (45×45×45cm) made of black Plexiglas and dimly illuminated. The animals were placed in the center of the arena, and were left to explore it for 10 min. The sessions were video recorded, and the distance travelled in the maze as well as the speed was tracked with ANY-maze Software 7.0 (Stoelting Co). Data were analyzed in 1 min bins.

### Novel object recognition and location

The hippocampus-dependent memory was assessed using a novel object location and novel object recognition tasks. Assays were performed as described elsewhere (Andres-Alonso et al., 2019). Briefly, training and testing were done in an open field square arena made out of plastic (50 cm x 50 cm). The mice were habituated in the empty arena for 20 min. Subsequently, 2 identical objects were introduced to the maze and the animals could explore the objects in 2 training session, 20 min each. 2 h afterwards, in the memory test phase, one object was replaced with an unfamiliar, unknown object (novel object recognition). Afterwards, the location of one object was changed (novel object recognition). During all sessions, the amount of time the animals explored the objects was recorded, and a discrimination index was calculated using the formula DI=[(T_new_-T_familiar_)/(T_new_+T_familiar_)]*100. The arena was cleaned with 5% ethanol before and after each animal was tested.

### Y-maze Object Recognition

Y-maze Object Recognition short-term memory task was performed according to Creighton et al. (2019). During training session, two identical objects (3D printed rectangle) were placed at the end of arms B and C and the animal was left to explore both objects for 10 min. In order to assess short-term memory, retrieval performance was tested 3 h after training with one of the familiar objects replaced by a novel one (wooden cube). The mice were placed in the arena, and had 5min to explore both objects. During both sessions the time animals explored the objects was scored, and a discrimination index was calculated, in the same manner as for the novel object recognition test.

### Image analysis of murine brain sections

All quantifications were done within the distal CA1 region of the hippocampus. Fiji/ImageJ software(Schindelin et al., 2012) was used to calculate maximum intensity projection from five optical sections for each channel. CREB and pCREB immunoreactivity of every single nucleus in a 100µm stretch was measured in arbitrary units of pixel intensity. ROIs were defined based on NeuN and DAPI. The neuronal loss was quantified based on NeuN staining (the number of neurons/CA1 length). For the quantification of microglia and astrocytes the number of cells based on overlaid DAPI and Iba-1 or GFAP signal was quantified within the rectangular ROI. Aβ plaques were counted in the region where the neuronal loss was quantified. For the quantification of synaptic density the openview (Tsuriel et al., 2006) software was used.

### Image analysis of hippocampal cultures

For quantitation of nuclear staining intensity (pCREB or CREB) somatic regions were sequentially scanned using the 63x objective (Leica) in both: Leica TCS SP8-STED and Leica TCS SP5 systems. Image format was set either to 1024×1024 or 512×512 with optical zoom 4. Average intensity images from three optical sections (z-step 300 nm) were generated and intensity measurements were performed within the ROIs defined by DAPI. The values were normalized to the mean intensity of the control group. Unprocessed images were analysed using Fiji/ImageJ software(Schindelin et al., 2012). For visualization of the quantified channel a fire look-up table (LUT) was used.

Synaptic density was quantified in secondary dendrites using Fiji/ImageJ (Schindelin et al., 2012). The number of synaptophysin and Shank3-positive puncta was divided by the length of the dendritic segment. Intensity of the GluA1 channel was measured in ROIs defined by a Shank3 mask as a readout for surface AMPAR expression. All analysis was performed on raw, unprocessed images. For some representative images linear contrast-enhancement (histogram normalization) was applied equally for all groups. The figures were created with Adobe Illustrator or Photoshop software.

### Statistical analysis

All analysis was performed by a researcher blind to experimental groups (treatments, genotypes, transfected constructs). Statistical analysis was performed in Prism 8.0 (GraphPad) with the exception of SPECT data analysis where the Matlab (R2017b) and whole-cell patch-clamp recording where IBM SPSS Statistic (Version 21) was used. To choose the appropriate statistical analysis, a normality test (Shapiro-Wilk test) was performed. The type of statistical test used for each experiment, significance levels and the n numbers are reported in the corresponding figure legends. Merged values are represented as mean ±S.E.M. For qPCR data Ct values were subjected to both Dixon’s and Grubb’s outlier test.

### Data and materials availability

Plasmids generated in this study (Marked as ‘this paper’ in the Supplementary Table 2) will be made available on request after completion of a Materials Transfer Agreement. Antibodies and other reagents are available from their respective sources. All data are available in the main text or the supplementary materials.

## Acknowledgments

The authors gratefully acknowledge the professional technical assistance of M. Marunde, C. Borutzki, S. Hochmuth, K. Böttger, T. Stöter, and O. Kobler. We would like to thank Dr. Schilling and Probiodrug for the generous gift of the TBA2.1 mouse line and Dr. Głów for the help with virus production. We would like to thank Dr. A.V. Failla from the UKE Microscopy Imaging Facility (DFG Research Infrastructure; RI_00489) CNI. This work was supported by grants from the Deutsche Forschungsgemeinschaft (DFG) (Kr 1879/9-1/FOR 2419, FOR 5228 RP6; Kr1879/10-1; CRC 1436 TPA02 and Z01; Research Training Group 2413 SynAGE, TPA3), BMBF ‘Energi’ FKZ: 01GQ1421B, The EU Joint Programme— Neurodegenerative Disease Research (JPND) project STAD (FKZ: 01ED1613) and Leibniz Foundation SAW 2017, 2018, 2019 ‘Neurotranslation’, SyMetAge and ‘SynERCA’ to MRK. People Programme (Marie Curie Actions) of the European Union’s Seventh Framework Programme FP7/2007-2013/under REA grant agreement no. [289581], H2020 Priority Excellent Science, Marie Sklodowska-Curie Actions (MSCA) MC-ITN NPlast to KMG, CS, MRK. DFG CRC 779 TPB8 to AK. U.S. Alzheimer’s Association, grant Ref. No. AARFD-17-503612 to GNB. DAAD/CAPES scholarship, Alexander-von-Humboldt Foundation Fellowship (1756/14-1) and CBBS NeuroNetwork grant to GMG, GRK1167 scholarship to RK.

## Author contributions

Conceptualization: MRK

Methodology: KMG, GMG, RR, RK, LS, HK, PY, AMO, IR, GB, SS, CS, MSW, GNB, AK, SR, LS, HK, SR

Investigation: KMG, GMG, RR, RK, LS, HK, PY, AMO, IR, GB, SS, CS, MSW, GNB, AK, SR, LS, HK, SR

Visualization: AK, KMG

Funding acquisition: AK, MRK

Supervision: AK, MRK

Writing – original draft: KMG, AK, MRK

Writing – review & editing: KMG, GMG, RR, RK, LS, HK, PY, AMO, IR, GB, SS, CS, MSW, GNB, AK, SR, AK, MRK, LS, HK, SR

## TABLES AND THEIR LEGENDS

**Table S1.** Summary of cases used in the study and PM delay of tissue collection for each case.

| Number | Gender | Age | PM delay, hrs | Group |
| --- | --- | --- | --- | --- |
| **1** | m | 93 | 49 | Alzheimer's disease |
| **2** | w | 74 | 14 | Control |
| **3** | w | 86 | 48 | Alzheimer's disease |
| **4** | m | 67 | 29 | Control |
| **6** | m | 50 | 19 | Control |
| **7** | m | 52 | 24 | Control |
| **8** | m | 76 | 32 | Alzheimer's disease |
| **9** | m | 65 | 24 | Control |
| **10** | w | 83 | 48 | Alzheimer's disease |
| **11** | w | 81 | 18 | Alzheimer's disease |
| **12** | w | 77 | 34 | Alzheimer's disease |
| **13** | w | 84 | 36 | Control |
| **14** | m | 82 | 46 | Control |
| **15** | m | 44 | 40 | Control |
| **16** | w | 53 | 28 | Control |
| **17** | w | 72 | 24 | Control |
| **18** | w | 84 | 12 | Control |
| **19** | m | 69 | 40 | Alzheimer's disease |
| **20** | m | 73 | 24 | Alzheimer's disease |
| **21** | w | 64 | 32 | Alzheimer's disease |
| **22** | w | 88 | 48 | Alzheimer's disease |
| **23** | w | 82 | 24 | Alzheimer's disease |
| **24** | m | 80 | 36 | Alzheimer's disease |

**Table S2.**
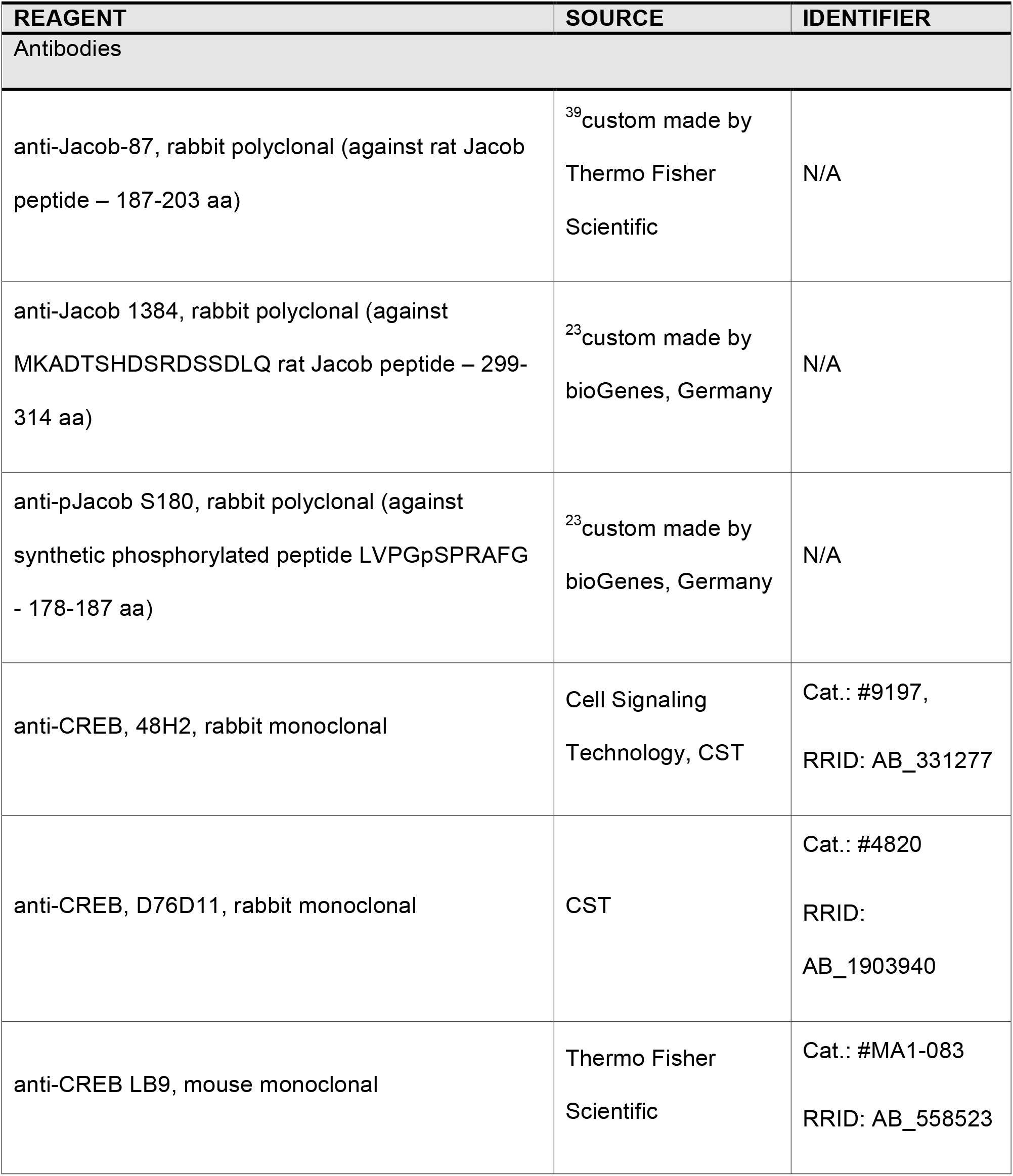
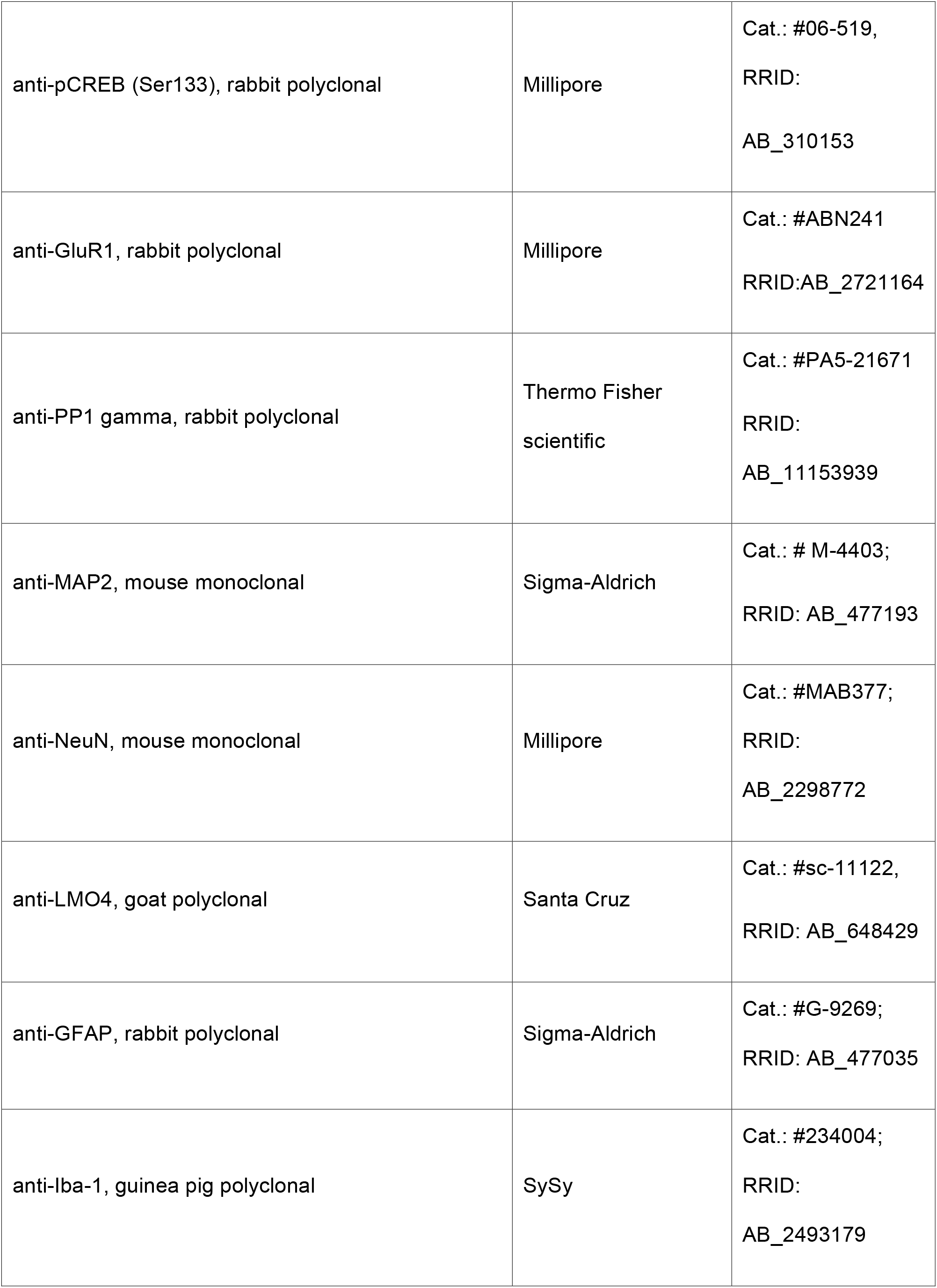
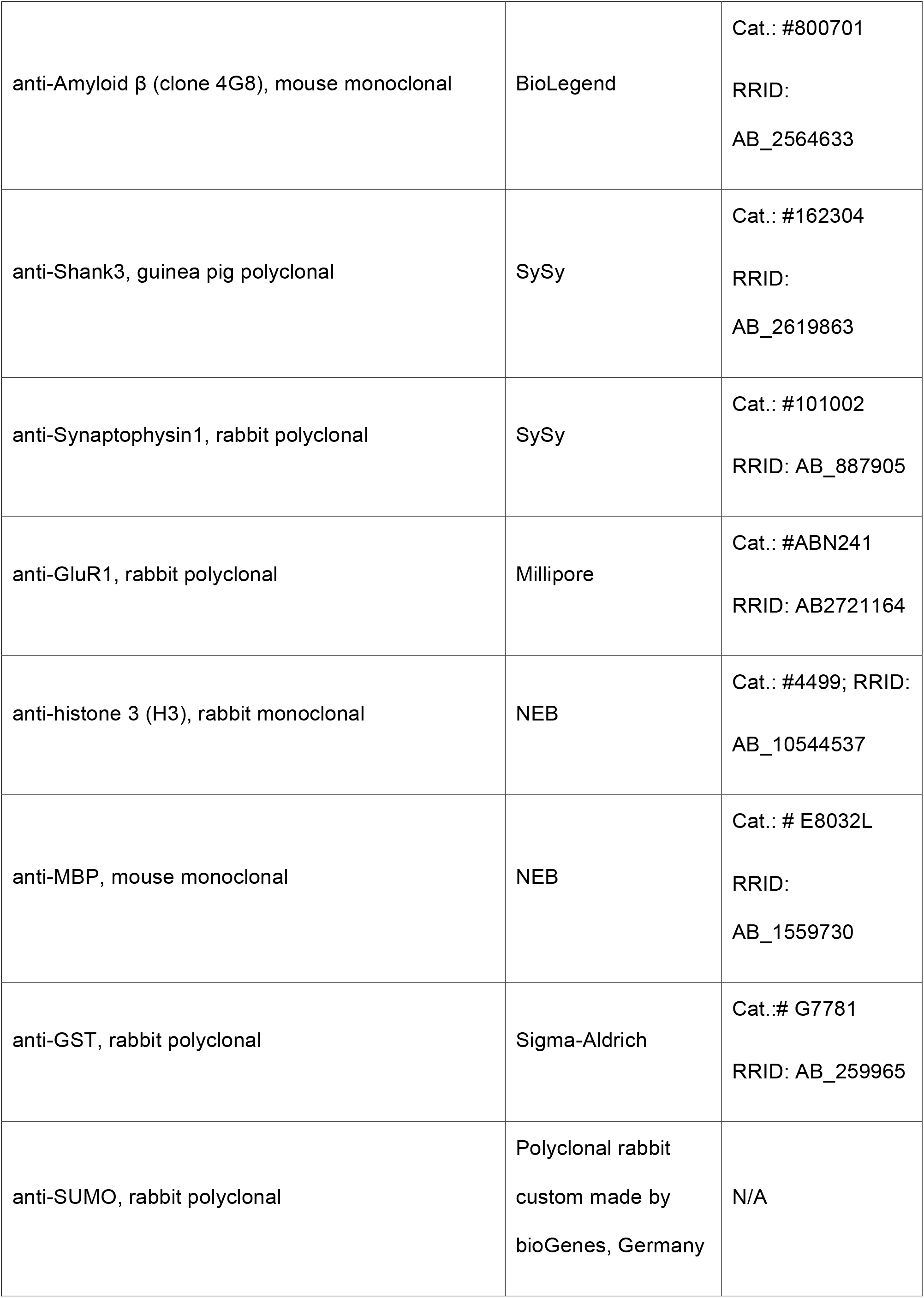
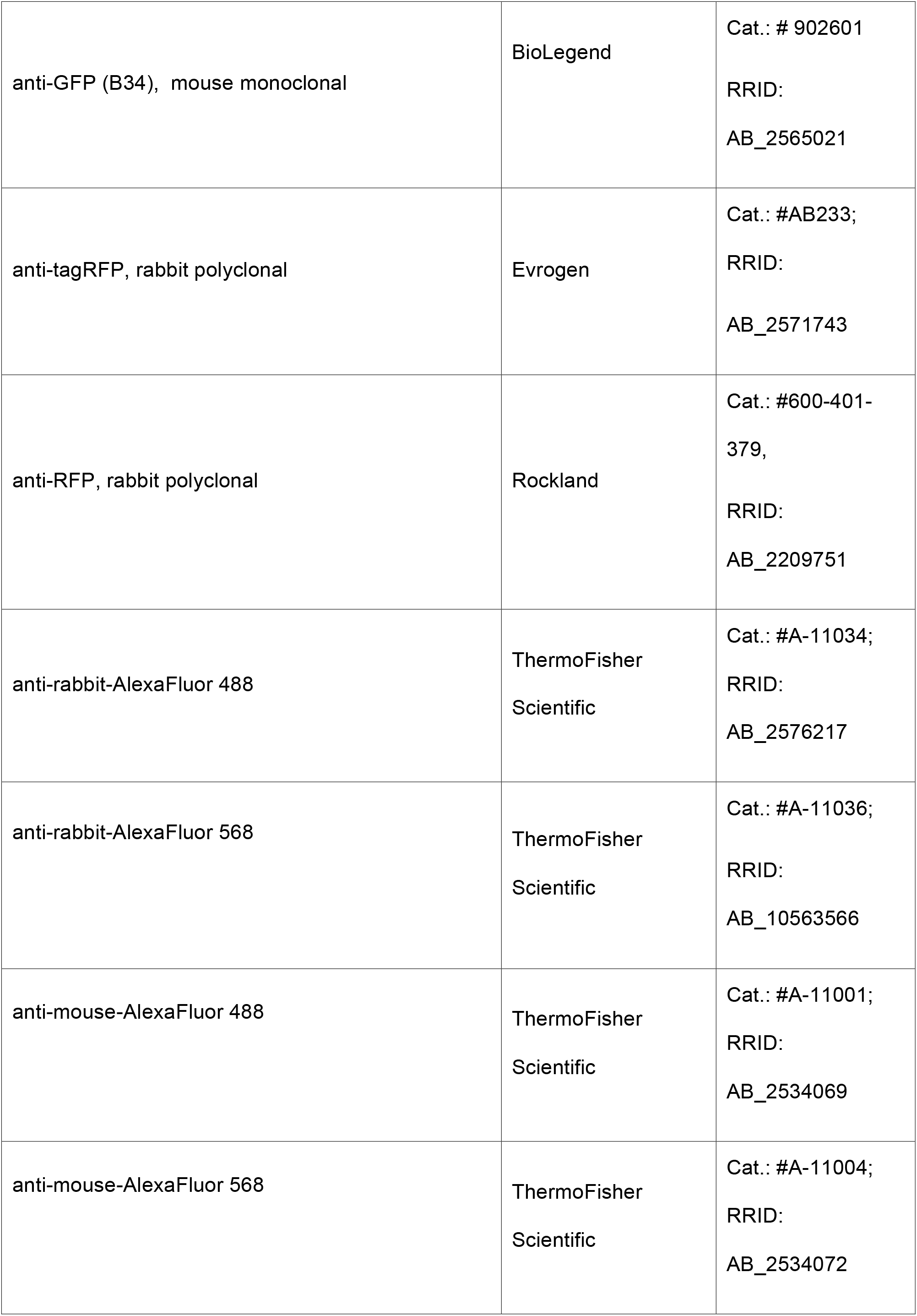
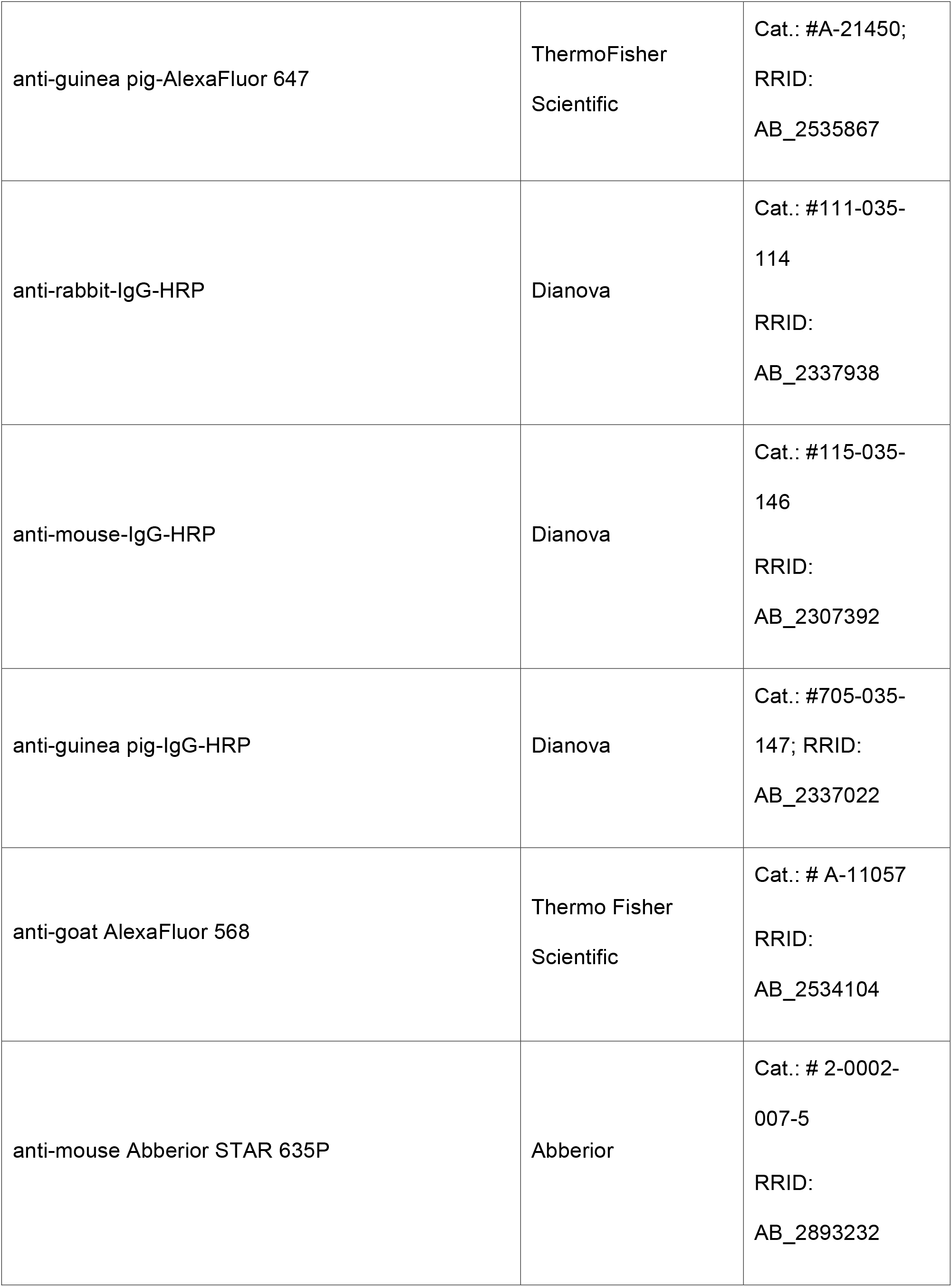
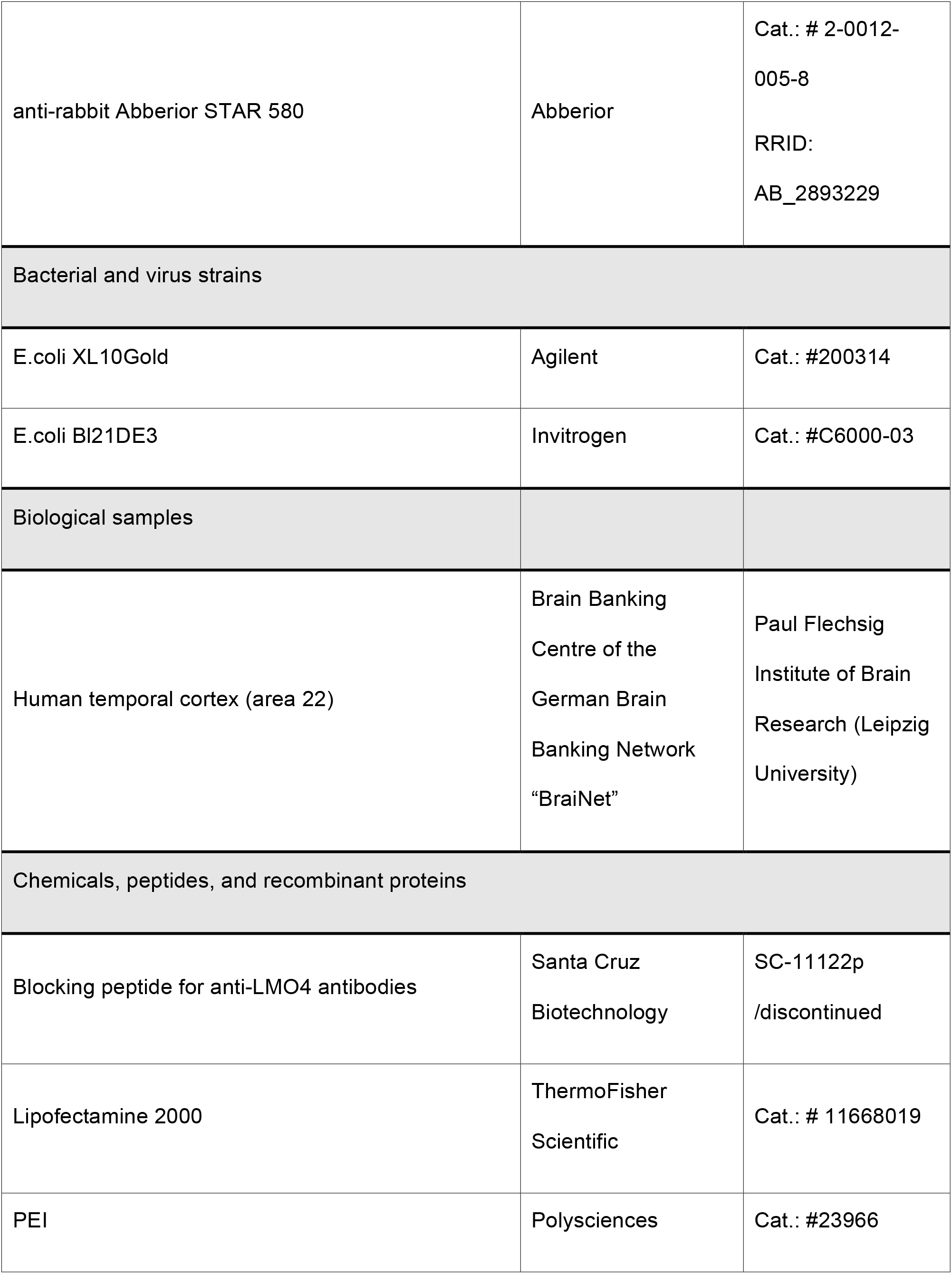
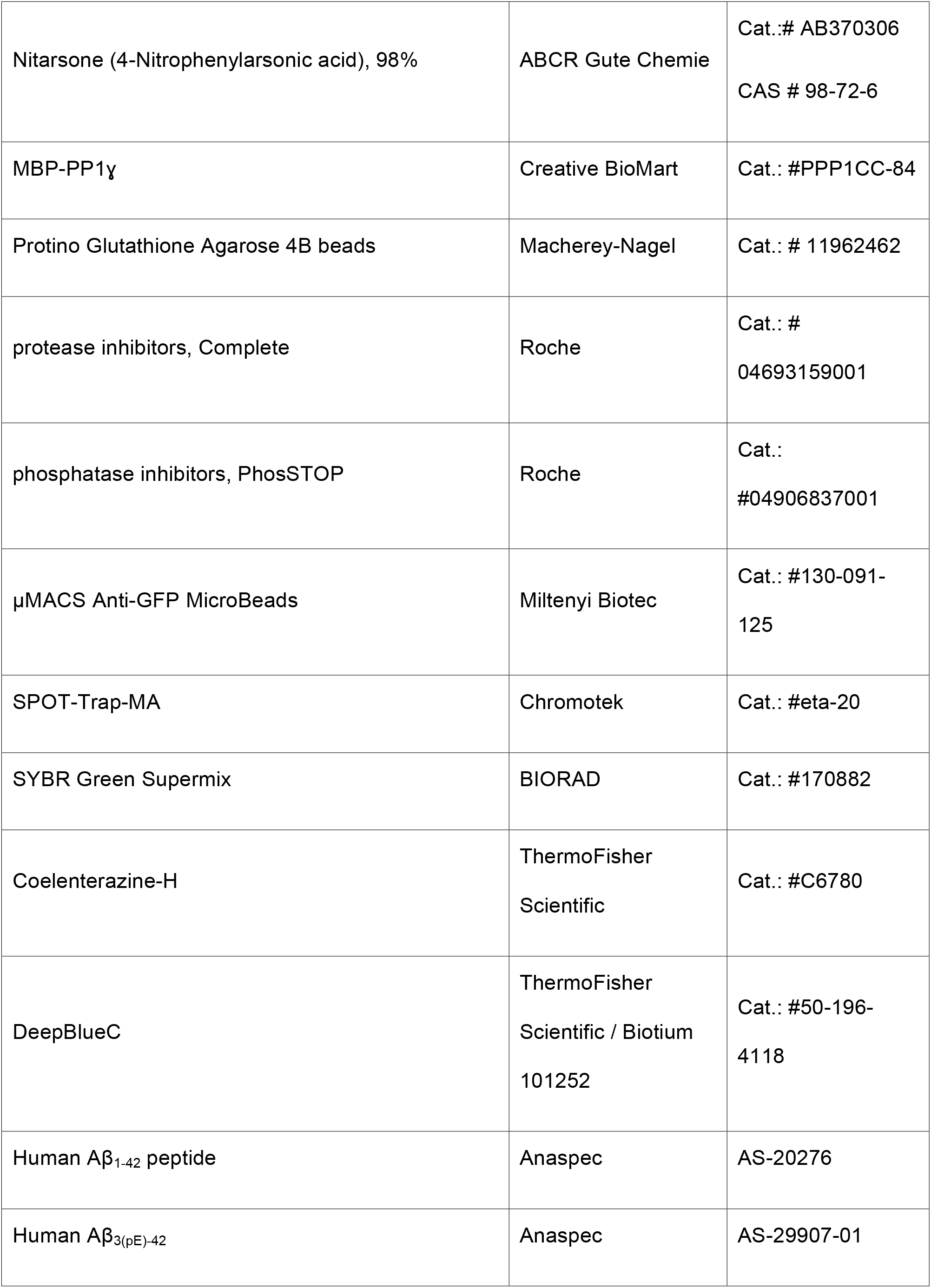
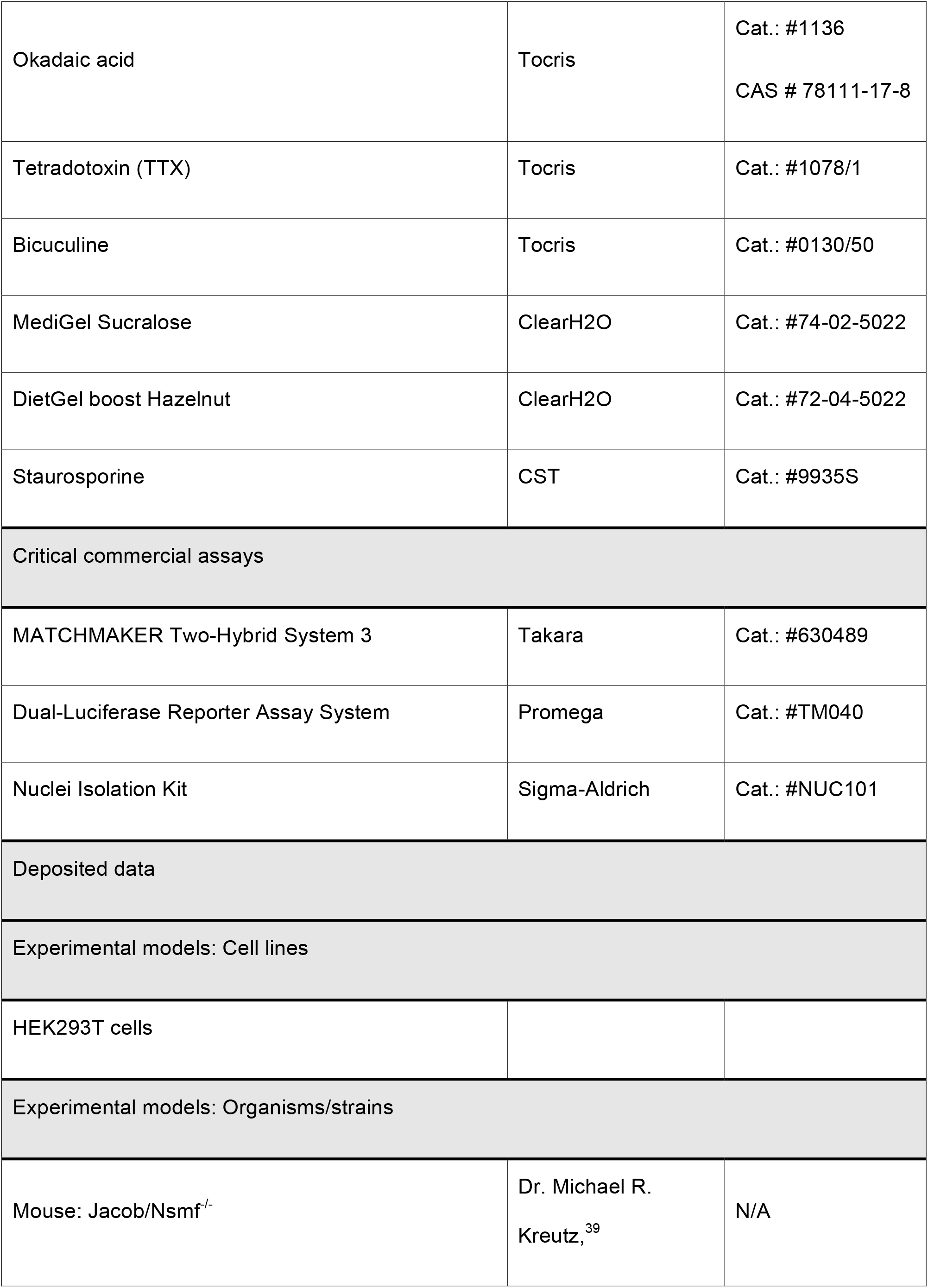
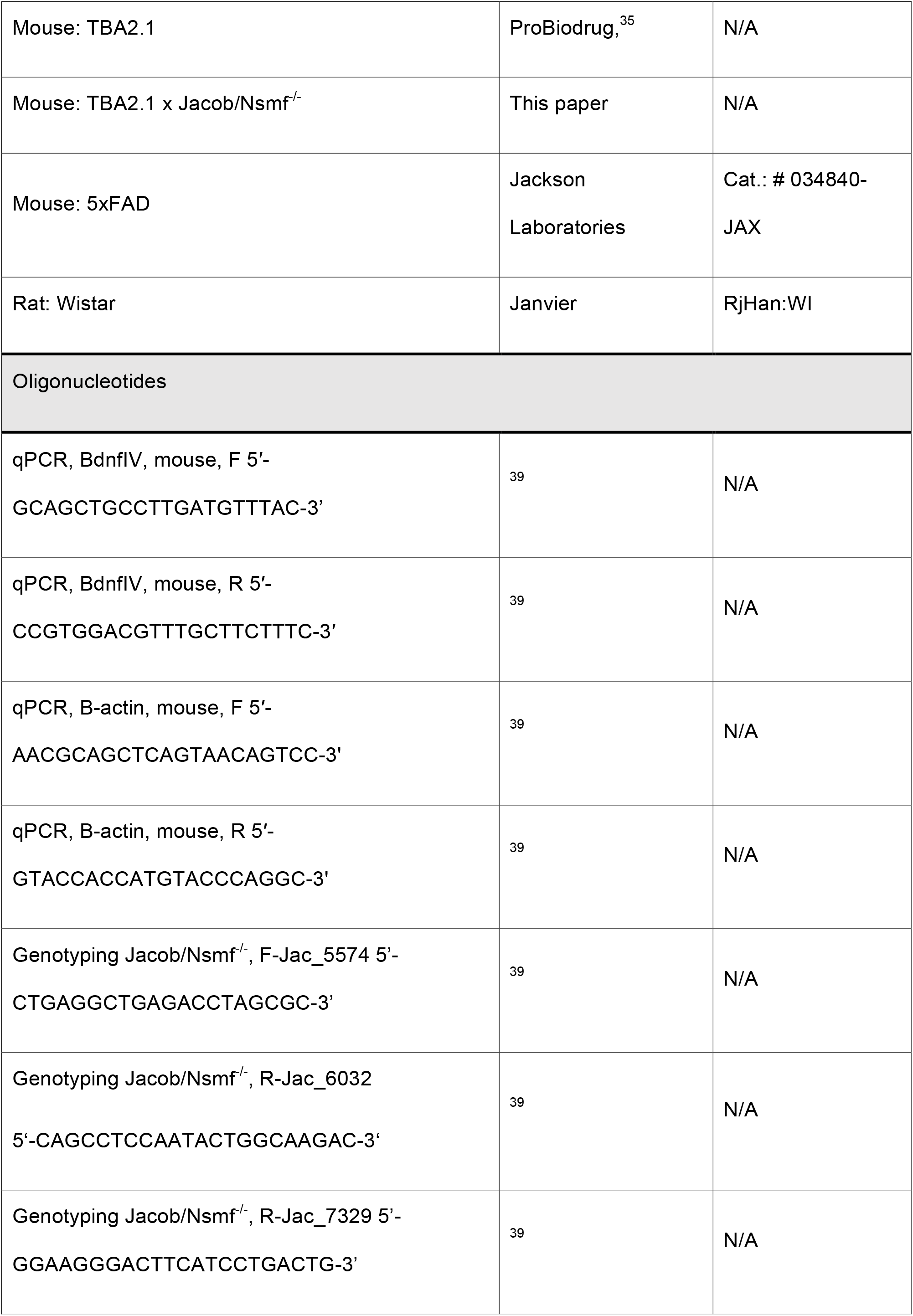
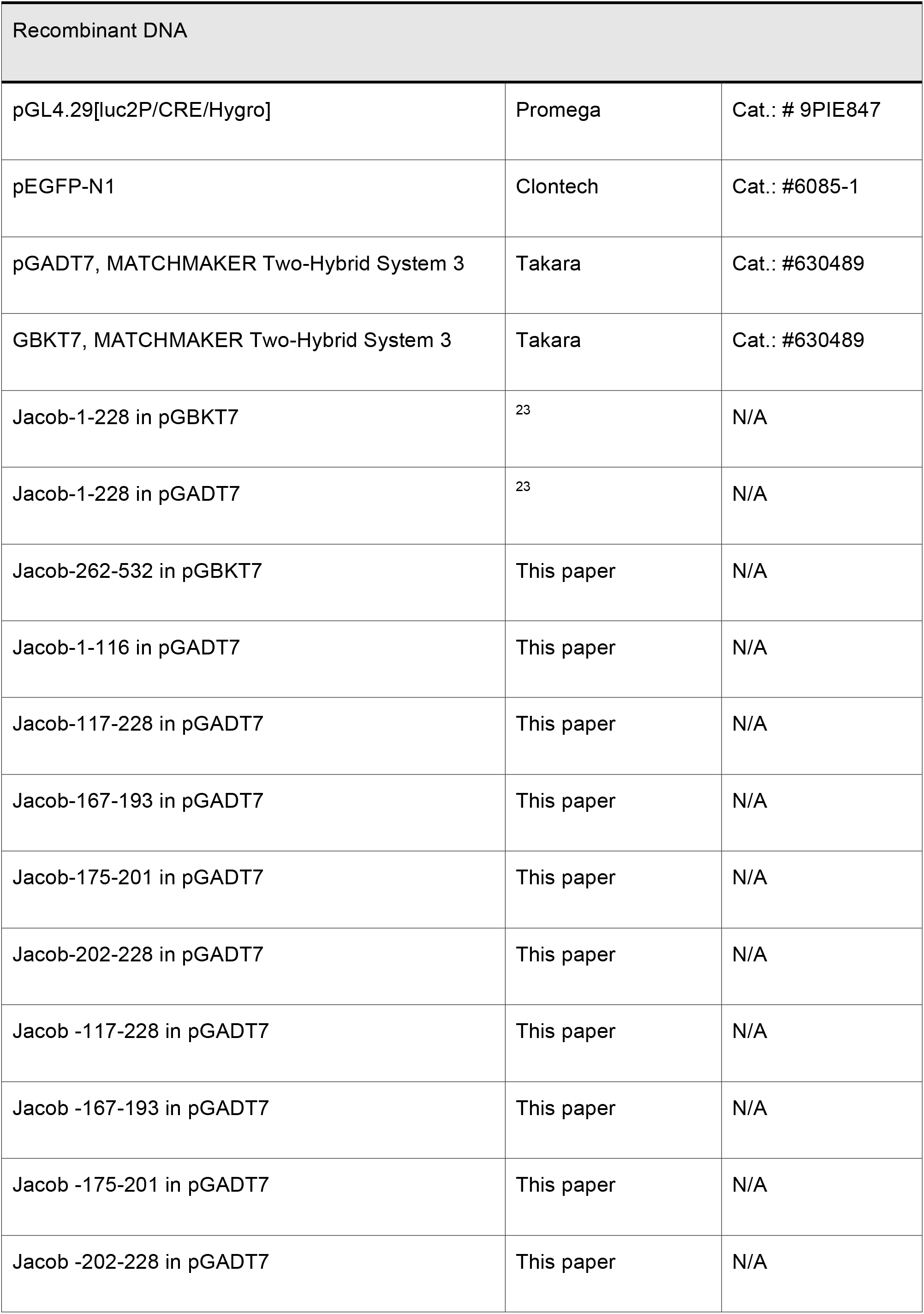
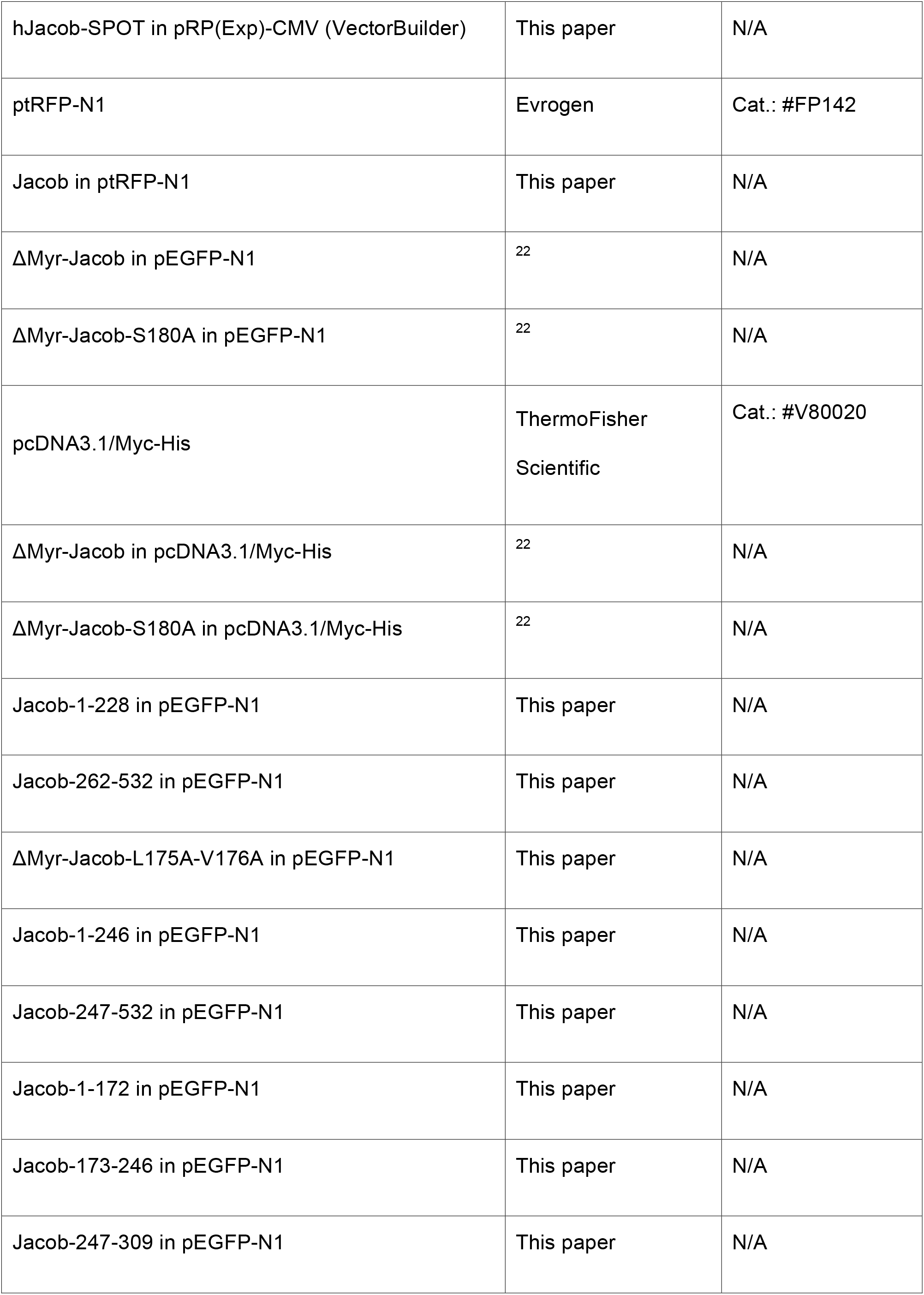
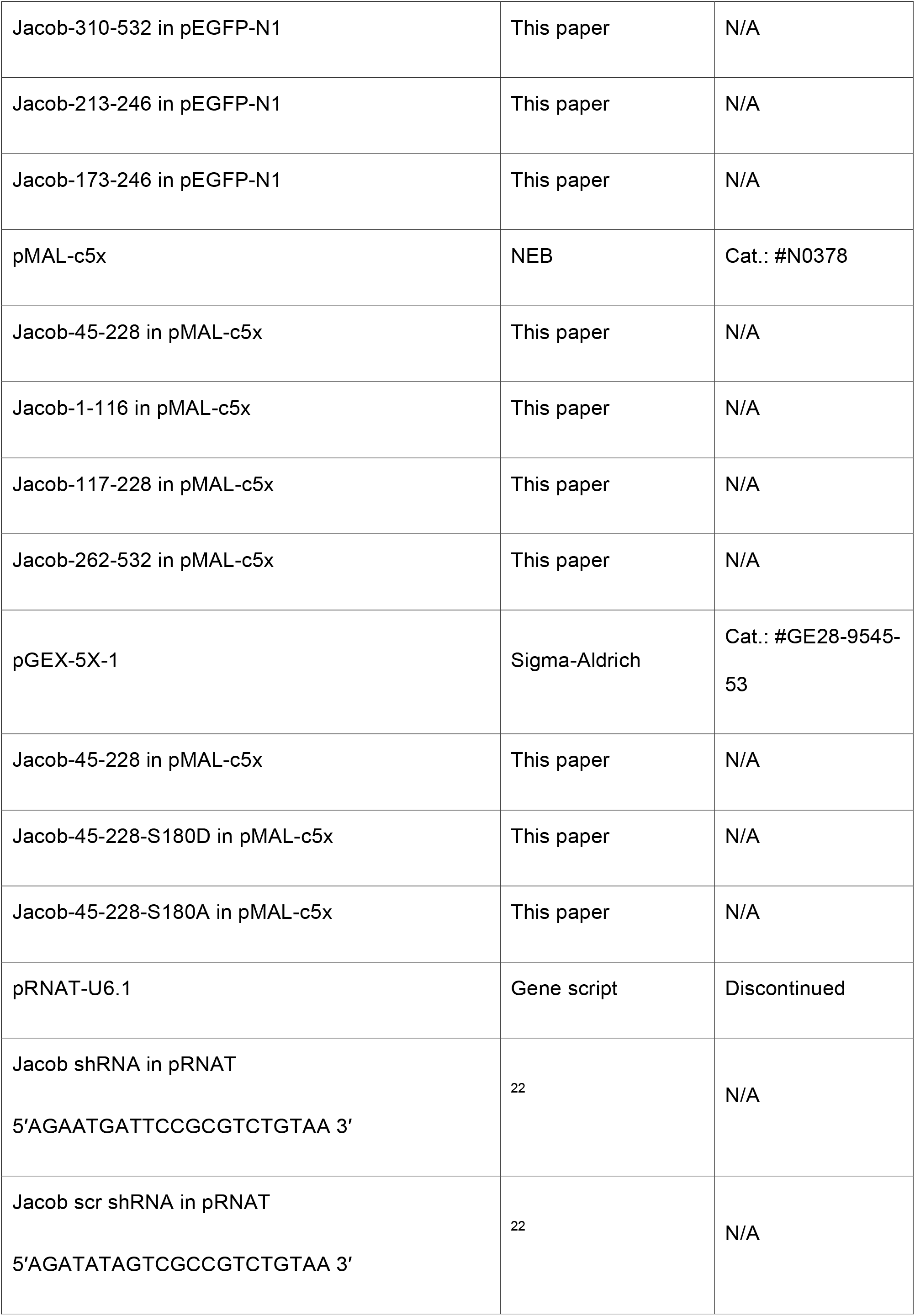
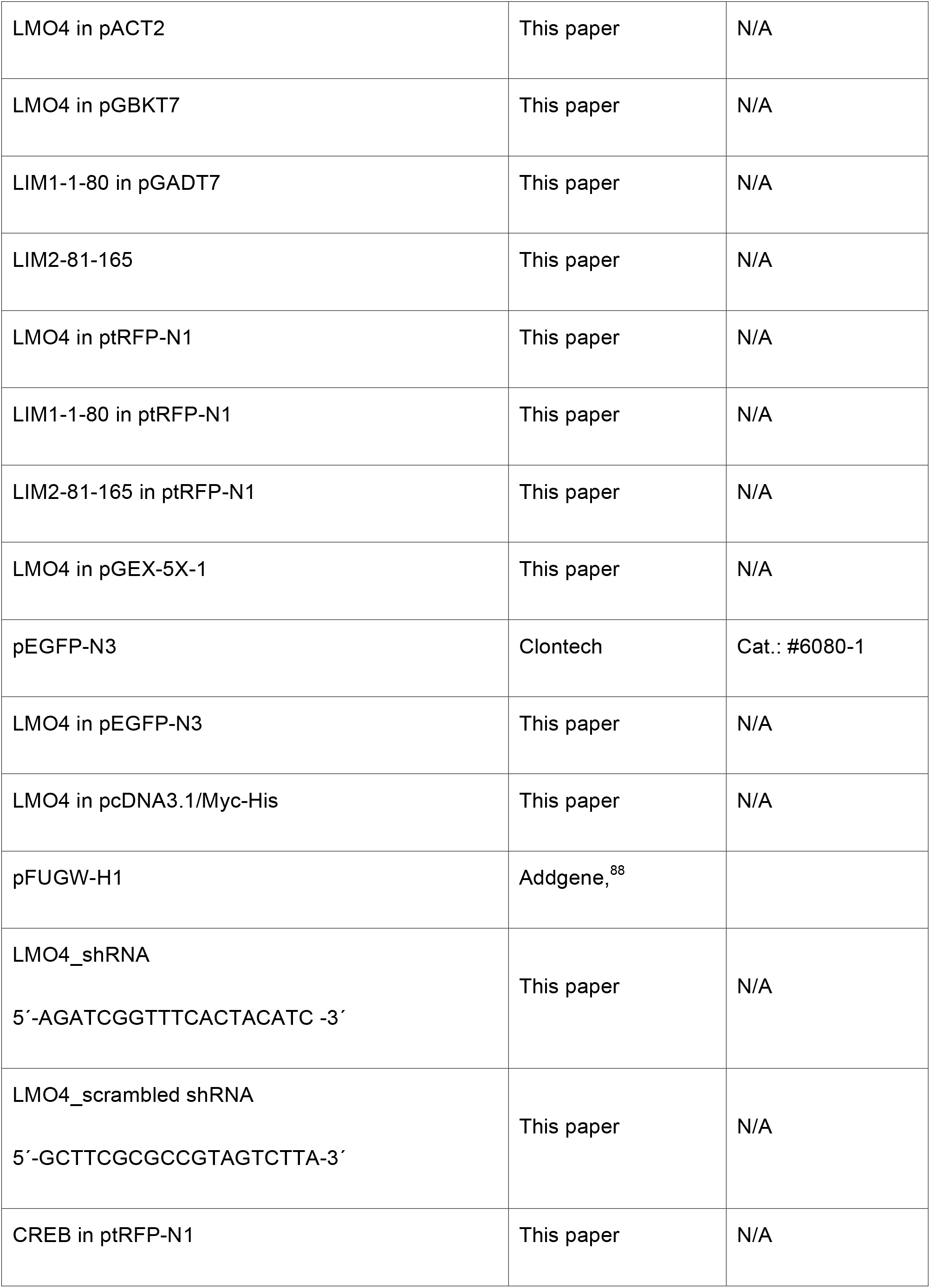
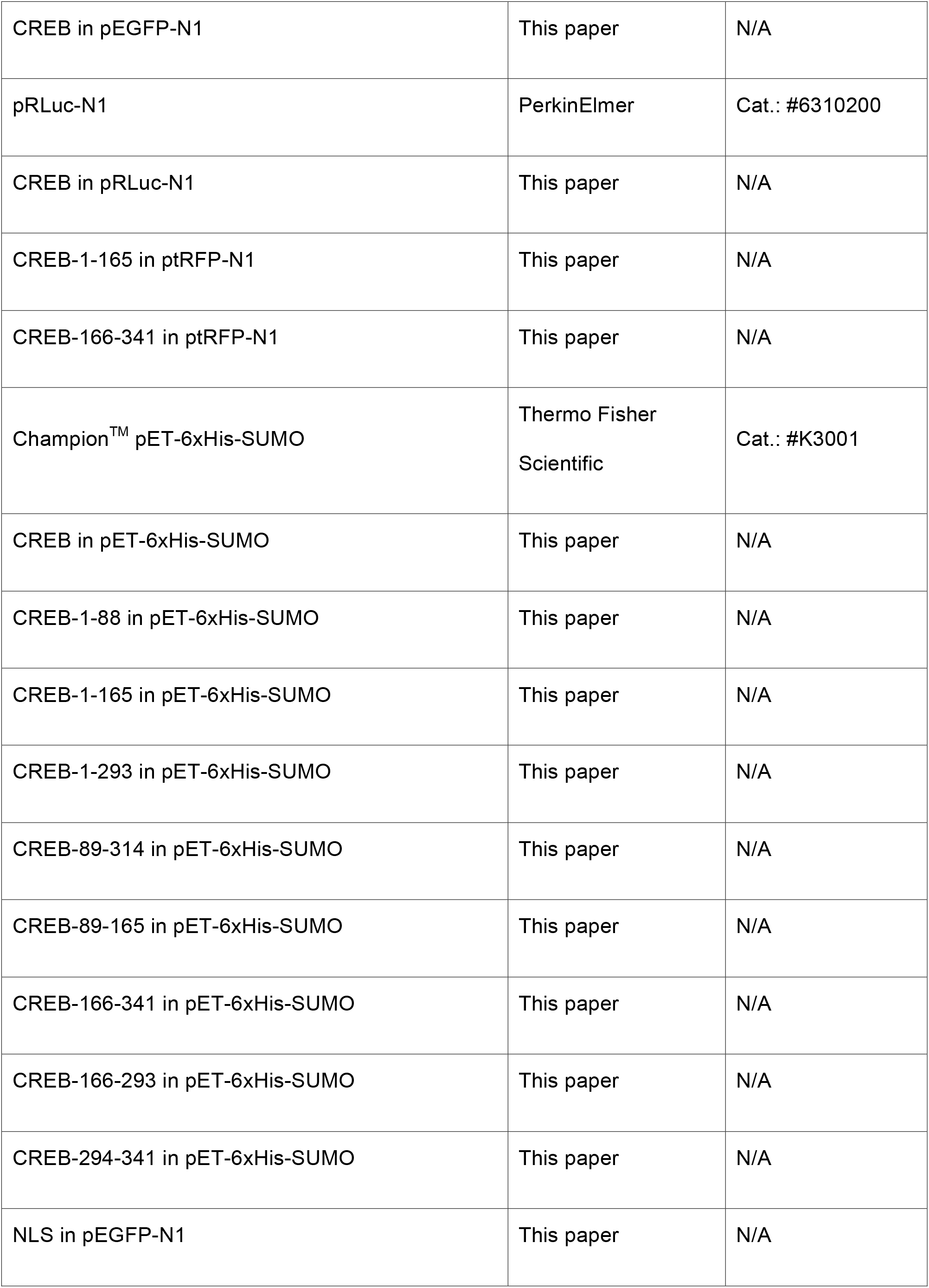
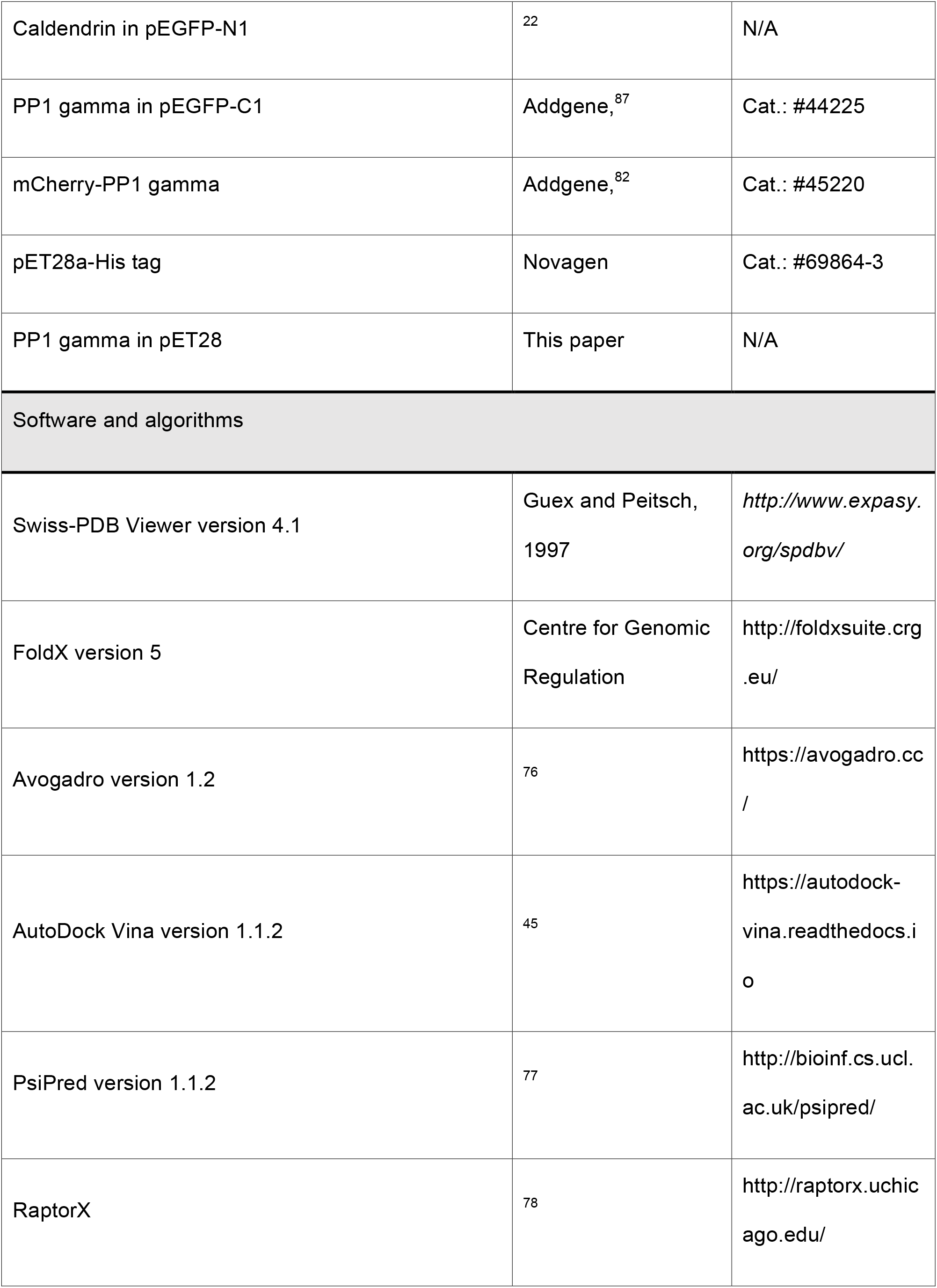
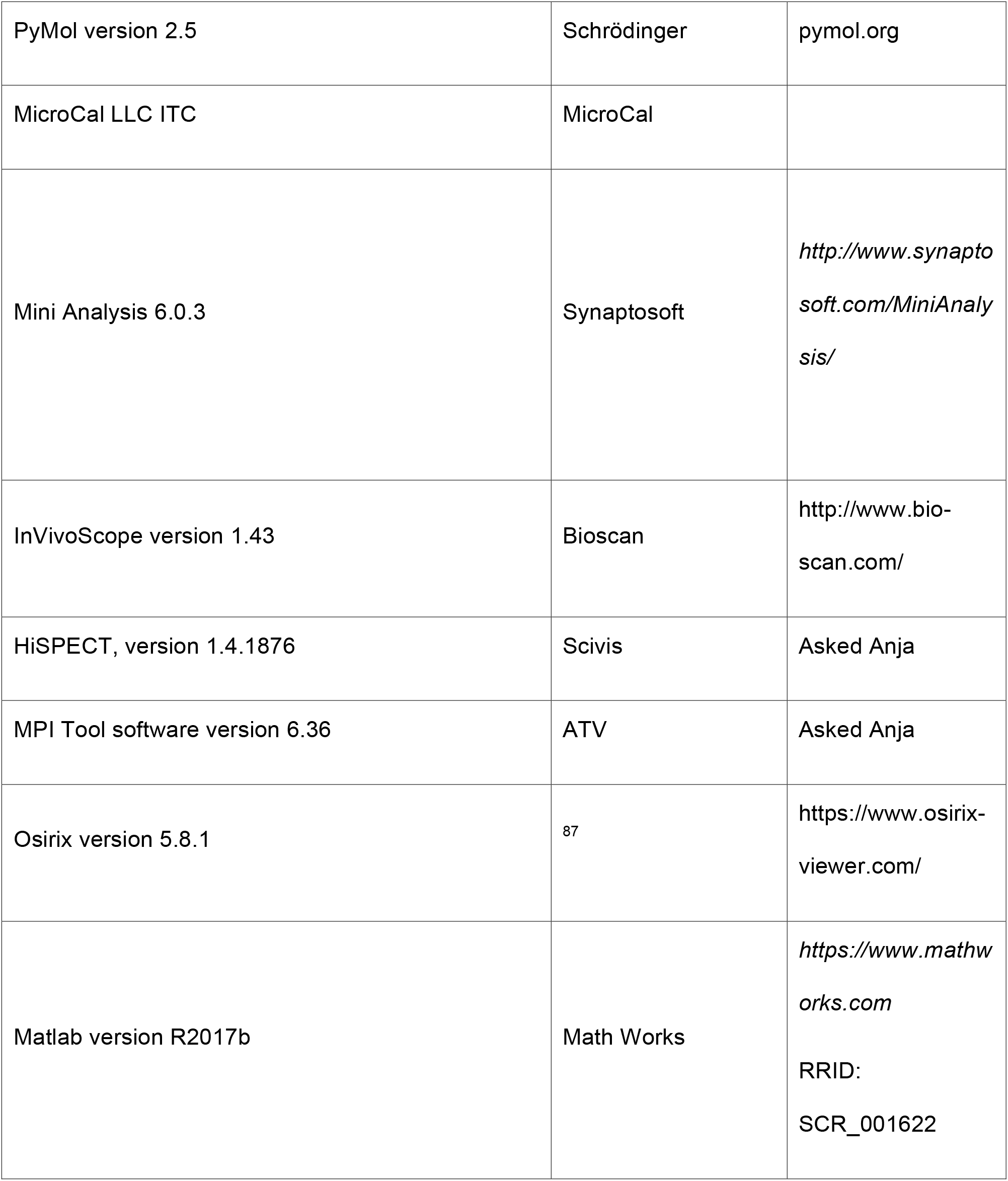
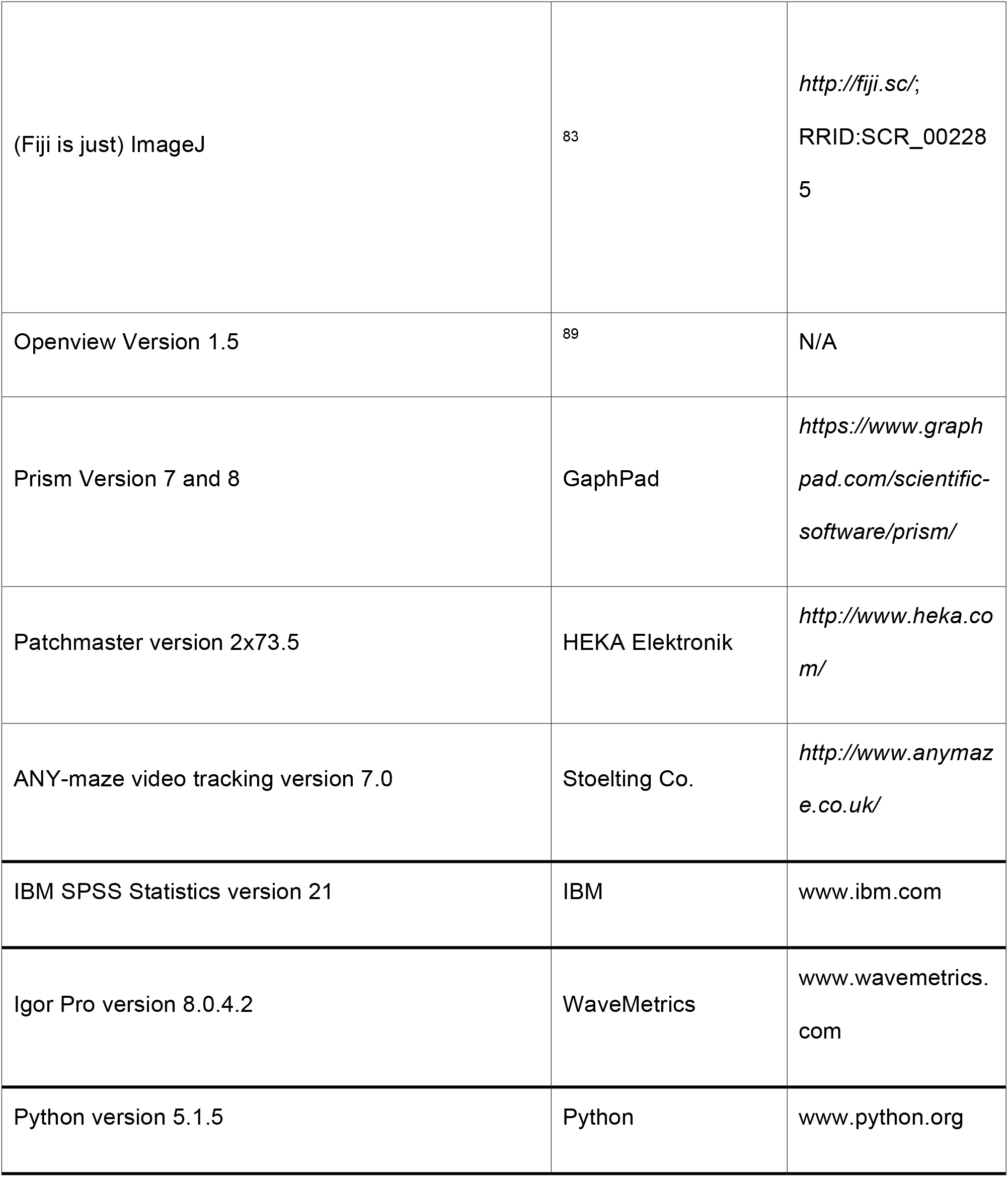
The antibodies, chemicals, kits, recombinant proteins, peptides, and software used in the study.

## EXPANDED VIEW FIGURES

**Fig EV1.**
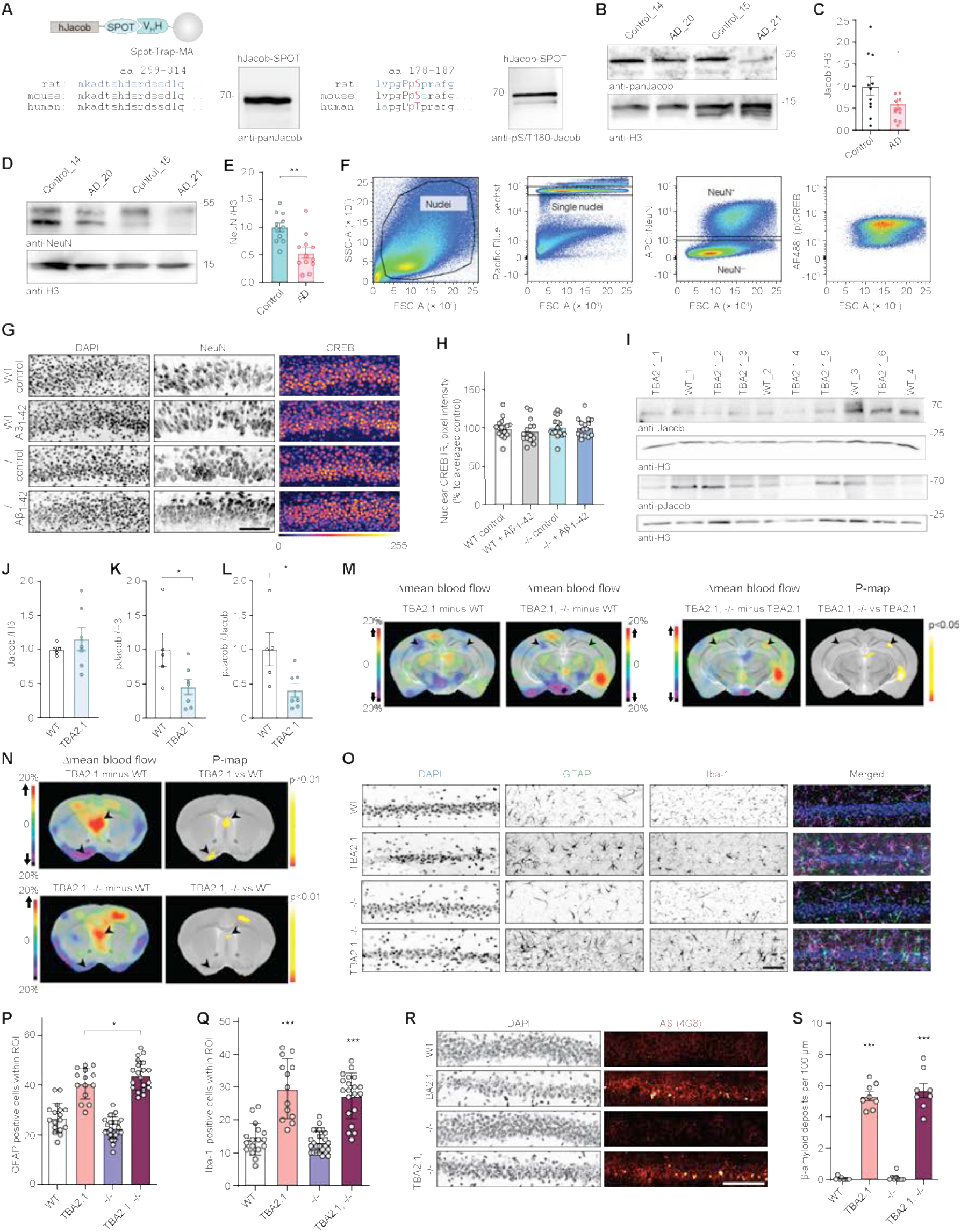
The role of Jacob in CREB shutoff. **(A)** Validation of Jacob and pJacob antibodies for detection of the human protein. Human Jacob fused to a Spot tag was expressed in HEK293T cells for antibody detection. The rat amino acid sequence of Jacob used for generation of pan-Jacob antibodies is highly conserved in human and mouse and the antibody effectively detects human Jacob. **(B, C)** Total Jacob protein levels are not significantly reduced in brain samples from Alzheimer’s disease patients as compared to the control group. (C) Bar plots representing the quantification of Jacob immunoreactivity normalized to H3. N=11-12 protein extracts from different subjects in each group. **(D, E)** Total NeuN protein levels are significantly reduced in brain samples from Alzheimer’s disease patients as compared to the control group. (C) Bar plots representing the quantification of NeuN immunoreactivity normalized to H3. N=11-12 protein extracts from different subjects in each group. **(F)** Scatter plots representing gating strategy used in FACS experiments for neuronal pCREB and CREB immunoreactivity quantification. **(G, H)** Acute (1h) Aβ_1-42_ treatment does not induce changes in pan CREB levels in organotypic hippocampal slices from Jacob (−/−) mice. (G) Representative confocal images of slices immunolabeled against pCREB, co-labeled with NeuN and DAPI. (H) Bar plot of CREB immunoreactivity averaged per slice, N=14-16 slices. **(I)** pJacob level and pJacob/panJacob ratio are decreased in TBA2.1 mouse line compared to WT animals. Representative images of the immunoblot probed with antibodies against pJacob, pan-Jacob, and re-probed with Histone3 (H3). **(J-L)** Bar plots representing the quantification of (J) Jacob, (K) pJacob levels and the (L) pJacob/Jacob ratio normalized to H3. N=5-7 hippocampal protein extracts. **(M)** Significant changes in cerebral blood flow between TBA2.1 and WT, TBA2.1, −/− and WT, and TBA2.1 and TBA2.1, −/− as determined by 99mTc-HMPAO SPECT measurements. Difference images overlay over a reference MR for comparison with TBA2.1 mice as described on panel labeling. Bregma -2.5. Statistically significant differences between TBA2.1 and double transgenic animals were detected in dorsal CA1. N=10 animals. (p<0,05) by two-tailed Student t-test. **(N)** Significant changes in cerebral blood flow between TBA2.1 and WT, TBA2.1, −/− and WT as determined by SPECT measurements. The statistically significant differences between TBA2.1 or double transgenic animal and WT were detected in lateral septal nucleus and the diagonal band nucleus. (p<0,01) by two-tailed Student t-test. **(O-Q)** The quantification of glial cells revealed no major differences between TBA2.1 and double transgenic animals (TBA2.1, −/−). (O) Representative confocal images of distal CA1 sections from 13 weeks old mice stained for GFAP, DAPI and Iba-1. Scale bar: 100 µm. (P) Bar plot representing the number of GFAP positive cells per rectangular region of interest. N=17-24 cryosections from 5-7 animals per genotype. **(Q)** Bar plot representing the number of Iba-1 positive cells per rectangular region of interest. N=17-26 cryosections from 5-7 animals per genotype. **(R, S)** The quantification of Aβ plaques revealed no major differences between TBA2.1 and double transgenic animals (TBA2.1, −/−). (R) Confocal images averaged from two sections of the molecular layer of 13 weeks old mice distal CA1 labelled for amyloid-β (4G8 antibody) and co-stained with DAPI. Scale bar: 100 µm. (S) Bar plot representing the number of amyloid-β positive puncta per 100 µm. N=8, number of cryosections from 2 animals per genotype. *p<0,05, **p<0.01, ***p<0.001, ****p<0.0001 by (J, K, L) two-tailed Student t-test or (N, P, S) two-way ANOVA followed by Bonferroni’s multiple comparisons test. All data are represented as mean ± SEM.

**Fig EV2.**
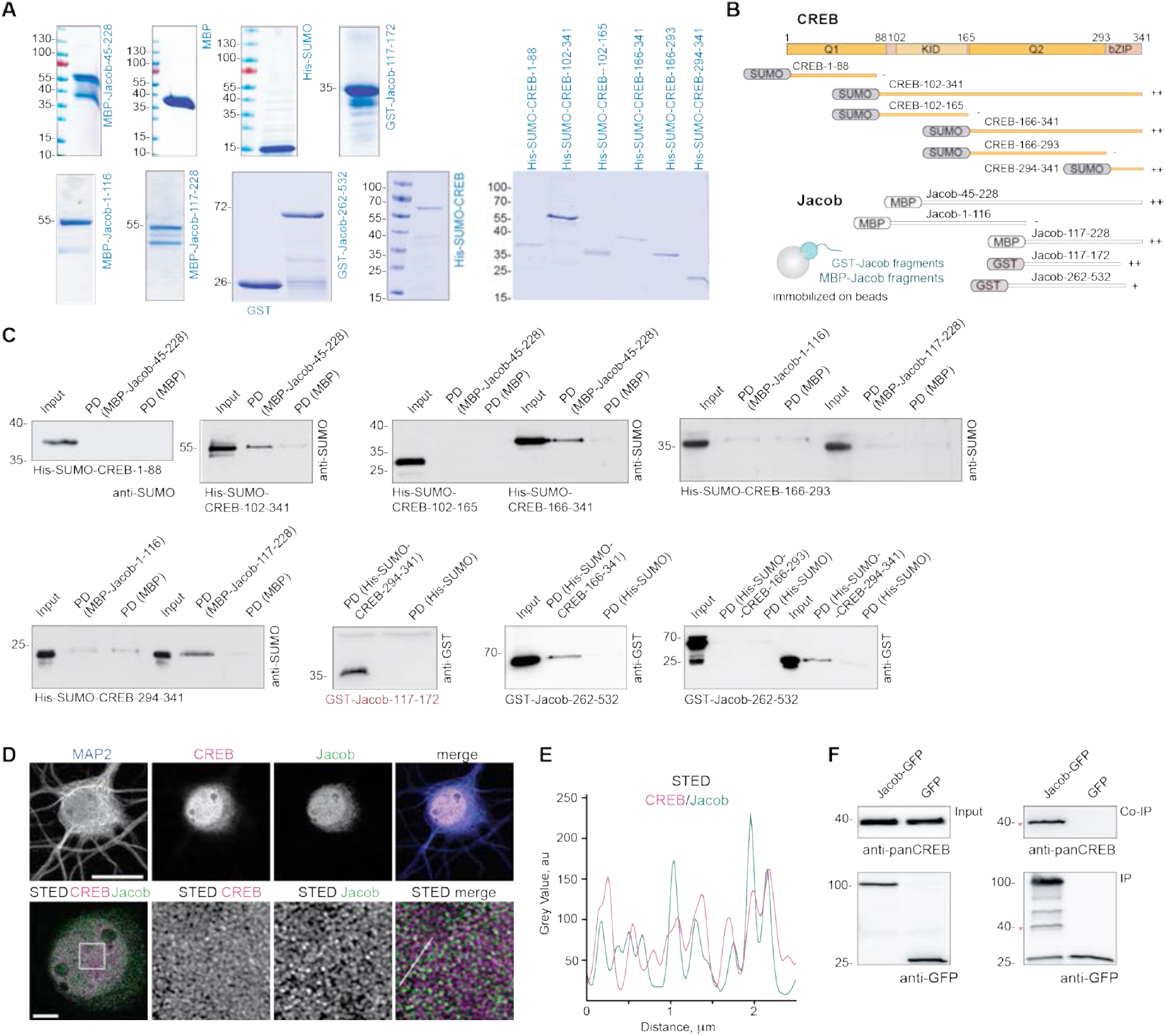
Mapping of the binding interfaces between Jacob and CREB. **(A)** Coomassie blue staining depicting the purity of bacterially produced proteins used for pull-down assays between CREB and Jacob. **(B)** Scheme representing the constructs used for mapping of the interaction sites in Fig. 1D and S1B. **(C)** The N-terminus of Jacob (117-172 aa in red) interacts with the bZIP domain of CREB, but not with the Q1 (1-88 aa), KID (102-165 aa), or Q2 (166-293 aa) domain. The C-terminus of Jacob (262-532 aa) shows weaker binding to the bZIP domain of CREB. Images of immunoblots representing pull-down assays performed with Jacob and CREB protein fragments depicted in the panel B. **(D, E)** Confocal and STED images show an association of CREB with Jacob in the nucleus of DIV16 hippocampal primary neurons. (D) The upper panel represents deconvolved confocal images. Lower panels depict deconvolved STED images. Scale bars: 20 µm and 5 µm respectively. Inserts are denoted by a white square. (E) Line profiles indicate the overlap of relative intensities for CREB and Jacob along a 2,5 µm line. **(F)** Endogenous CREB co-immunoprecipitate with overexpressed Jacob-GFP, but not GFP from HEK293T cell extracts. The asterisk denotes the CREB band from a membrane subsequently re-probed with an anti-GFP antibody.

**Fig EV3.**
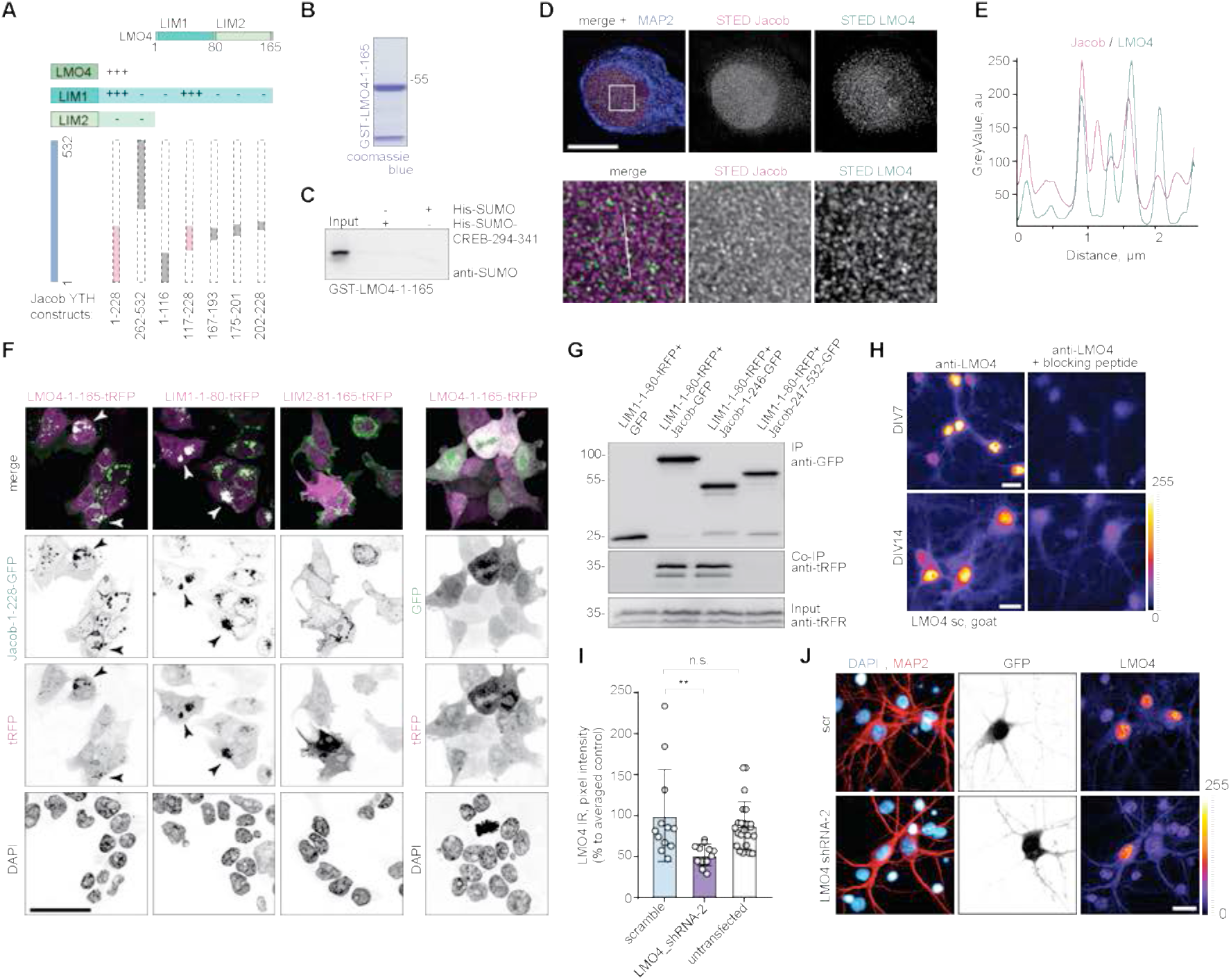
Confirmation of the LMO4-Jacob interaction, LMO4 antibody specificity and quantification of LMO4 knockdown efficiency, confirmation of Jacob-PP1 interaction, protein purity and loading controls for pull down experiments and studies on the interaction and network activity depending upon Jacob S180 phosphorylation. **(A)** Jacob-117-228 interacts with the LIM1 domain of LMO4 in Y2H. (+++) indicates a strong interaction while (–) indicates no interaction. Evaluation was based on the number of colonies growing in triple drop-out media. **(B)** Coomassie blue stained gel showing purity of GST-LMO4 used in the pull-down experiments. **(C)** Pull-down experiments revealed no interaction between His-SUMO-CREB-294-341 with GST-LMO4. **(D, E)** Super-resolution STED imaging revealed association of Jacob with LMO4 in the nucleus (D). DIV16 primary hippocampal neurons stained with antibodies against MAP2, LMO4, pan-Jacob. Upper panel - scale bar:10 µm, lower panel (inserts 5 µm x5 µm – denoted by the white square). (E) Line profiles indicate relative intensities for deconvolved STED channels along a 2.5 µm line. **(F)** Jacob-1-228-GFP co-recruits the LIM1 (1-80), but not LIM2 (81-165) domain of LMO4. Confocal images of HEK293T cells co-transfected either with Jacob-1-228-GFP or GFP together with LMO4 constructs. Arrows indicate co-recruitment. Scale bar: 40 µm. **(G)** LIM1-1-80-tRFP co-immunoprecipitate with Jacob-1-246-GFP, but not Jacob-247-532-GFP from HEK293T cell extracts. **(H)** A goat LMO4 antibody was used to stain LMO4 in hippocampal neurons (DIV 7 and DIV 14) in the presence or absence of the blocking peptide. Scale bar: 15 μm. **(I)** Representative, confocal images of hippocampal neurons transfected with shRNA targeting LMO4 or scrambled control. Reduction of nuclear LMO4 level was confirmed in immunocytochemistry with the goat anti-LMO4 antibody. Scale bar:10 μm. **(J)** Nuclear LMO4 staining intensity was downregulated in neurons expressing shRNA targeting LMO4 mRNA compared to scrambled-transfected (scr shRNA) or non-transfected cells. Data represented as mean ± SEM. n=12-23 nuclei. **p<0,021 by Kruskal-Wallis test followed by Dunn’s multiple comparison test.

**Fig EV4.**
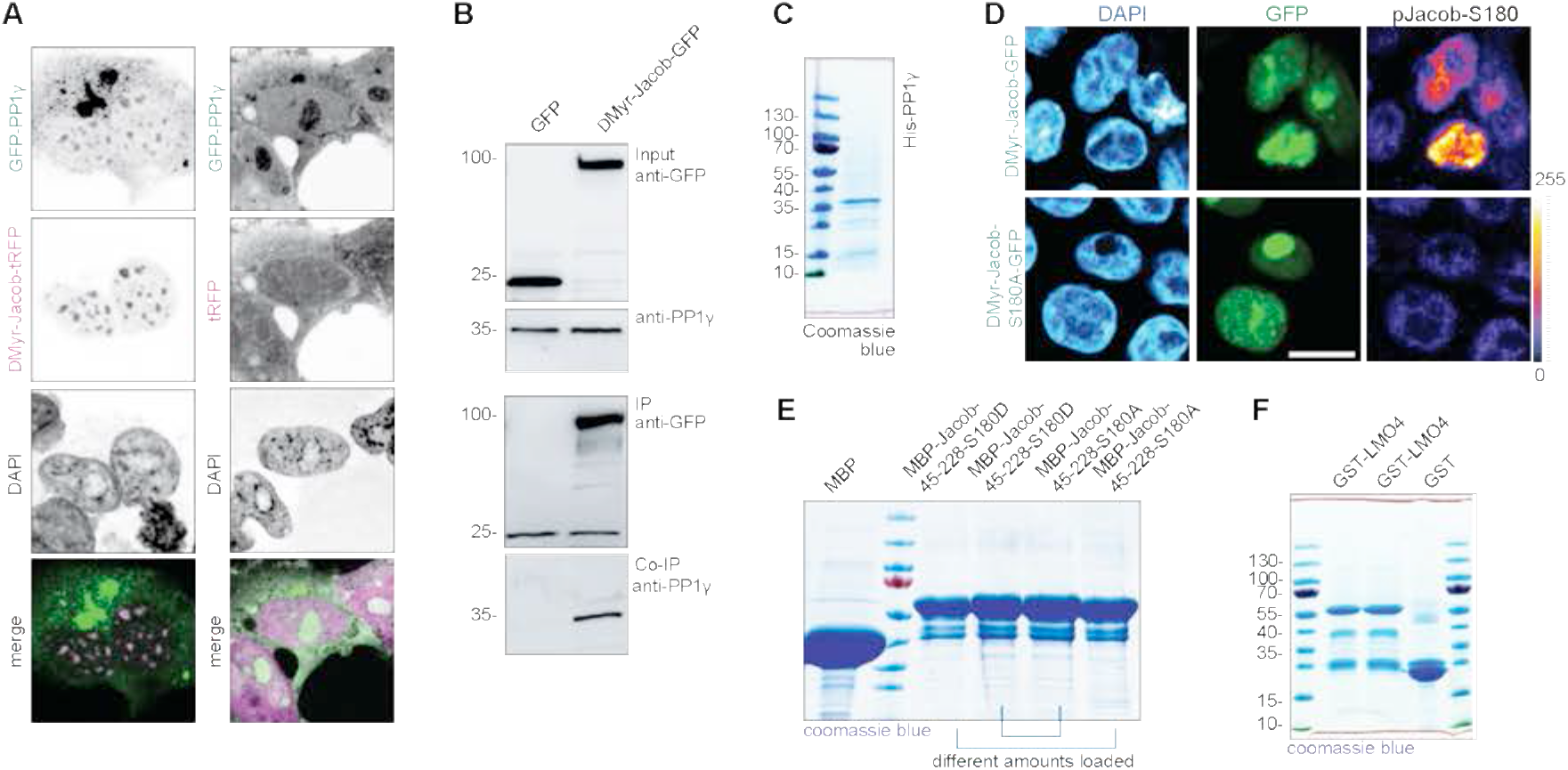
Jacob interacts with PP1γ. **(A)** Confocal images of HEK293T cells overexpressing GFP-PP1γ together with either ΔMyr-Jacob-tagRFP or tRFP control revealed nuclear co-clustering of both proteins. Scale bar: 20 µm. **(B)** ΔMyr-Jacob-GFP but not GFP co-immunoprecipitate with endogenous PP1γ from HEK293T cells extracts. **(C)** Coomassie blue stained gel showing purity of commercially available His-PP1γ. **(D)** Nuclear Jacob (ΔMyr-Jacob-GFP), but not nuclear phosphodeficient mutant (ΔMyr-Jacob-S180A-GFP) expressed in HEK293T cells is phosphorylated. Confocal images of HEK293T cells overexpressing Jacob and immunostained with anti-pS180Jacob antibodies. Scale bar: 10 μm. **(E)** Image of gels stained with coomassie blue showing the purity of bacterially produced Jacob mutants used for pull-down assay. **(F)** Images of gels stained with coomassie blue showing inputs for bacterially produced GST-LMO4 coupled to beads used for pull-down assay with Jacob and PP1γ.

**Fig EV5.**
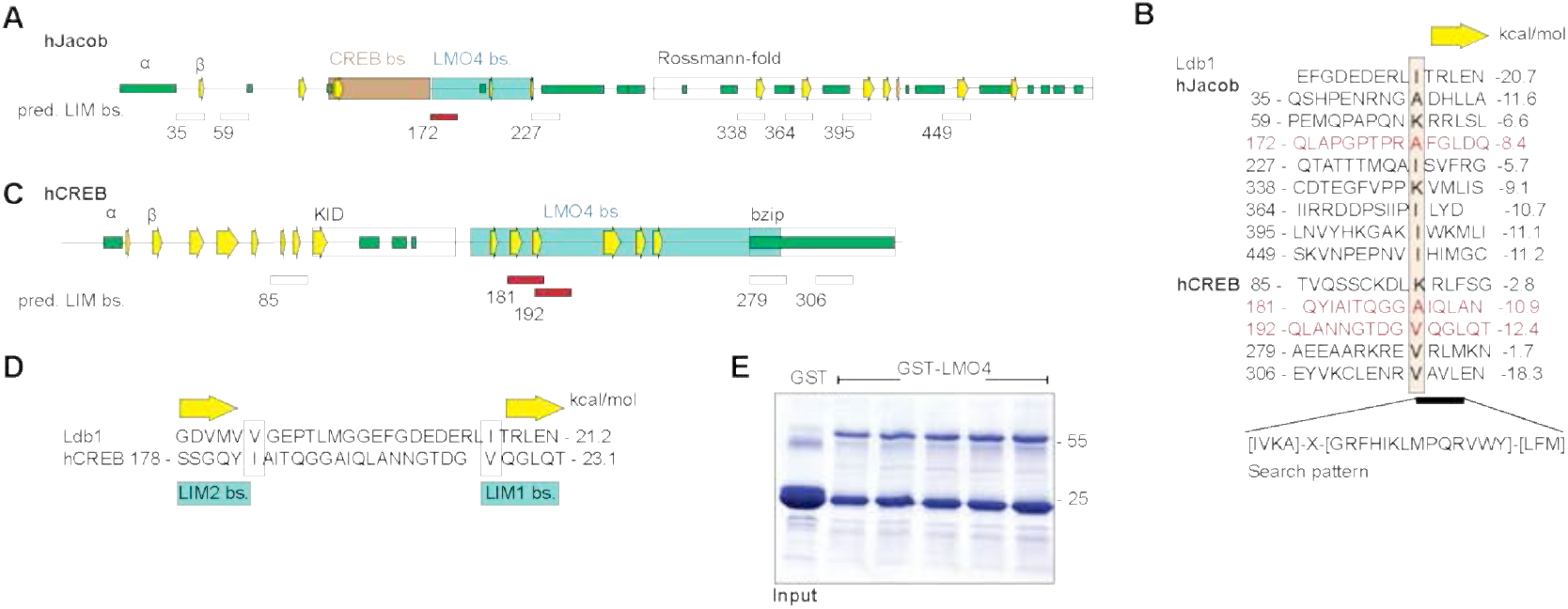
Nitarsone disrupts the Jacob-LMO4 interaction. **(A-D)** Predicted binding sites for LMO4 LIM domains in Jacob and CREB. (A) Schematic structure of human Jacob showing predicted secondary structures (helices, green; β-strands, yellow arrows) and experimentally determined binding regions for CREB (orange) and LMO4 (grey). The C-terminus of Jacob is predicted to have a Rossmann-fold similar to caspases. (B) The LIM1 binding peptide of Ldb1 is aligned to 8 sequences of Jacob that match the search pattern for the conserved hydrophobic residues and the adjacent β-strand. Structures of LIM1:peptides were modelled and free energy ΔΔG were calculated. Only the peptide starting at 172 (red) lies within the LMO4 binding region. In human CREB 5 matching peptides were identified. (C) Schematic structure of human CREB with labeled LMO4 binding region and known KID and bZIP domains. (D) The two peptides starting at 181 and 192 are within the LMO4 binding region and align to Ldb1 peptide where 181 binds to LIM2 and 192 to LIM1. **(E)** Image of gels stained with coomassie blue showing the purity of bacterially produced GST-LMO4 used for pull-down assay.

**Fig EV6.**
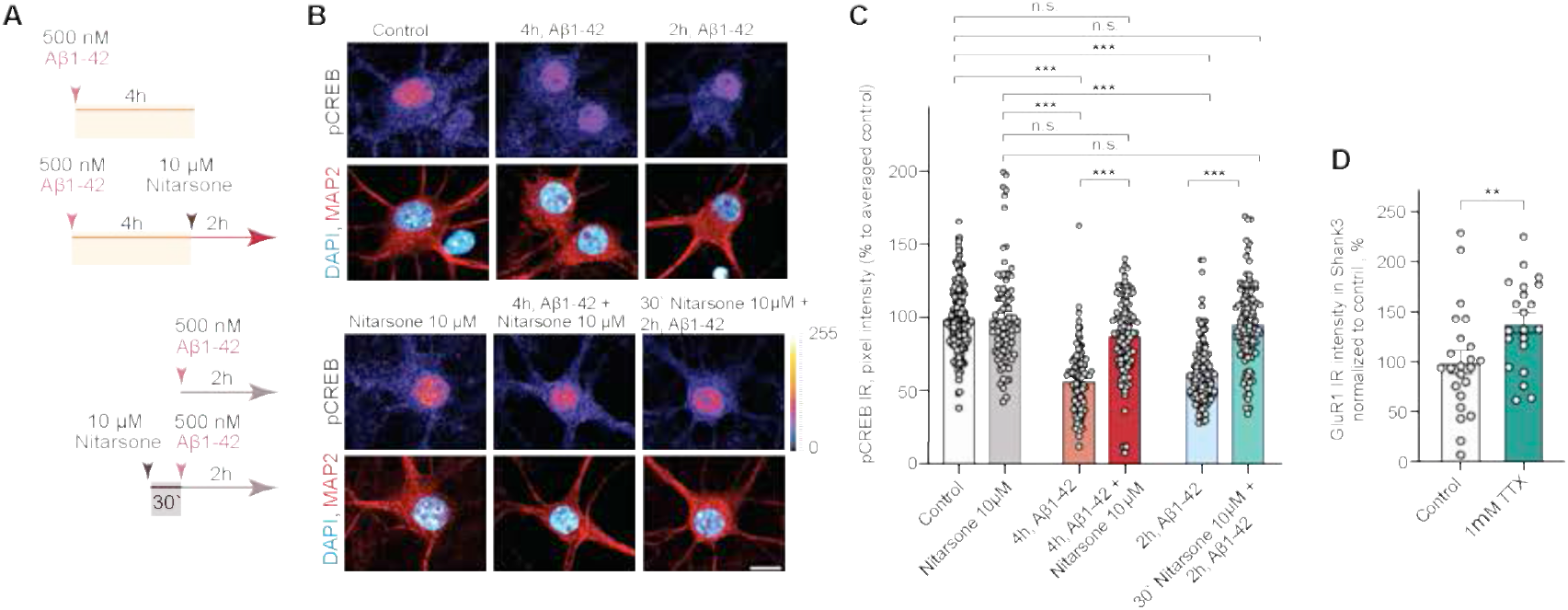
Nitarsone prevents Aβ-induced CREB shutoff in vitro. **(A-C)** Acute treatment with 10 µM Nitarsone rescuses Aβ_1-42_-induced CREB shutoff. (A) Scheme of the experimental design. The dissociated, hippocampal cell cultures at DIV16 were either pre-treated for 30 min with 10 µM Nitarsone and subsequently 2h with 500 nM Aβ_1-42_ or the drug was added 2h after the 500 nM Aβ_1-42_ treatment. The pCREB immunoreactivity was measured in comparison to vehicle control. (B) Bar plot representing nuclear pCREB immunostaining intensity normalized to vehicle control. N=88-101 from 5-7 independent cell cultures. ****p<0,0001 by two-way ANOVA with Sidak’s post hoc test. (C) Representative confocal images of hippocampal neurons. Lookup table indicates the pixel intensities from 0 to 255. Scale bar: 10µm. **(D)** Treatment with 1 µM TTX induced upregulation of GluR1 surface expression. N=21-23 dendritic segments from 3 independent cell cultures. **p<0,01 by two-tailed Student t-test. All data are represented as mean ± SEM.

**Fig EV7.**
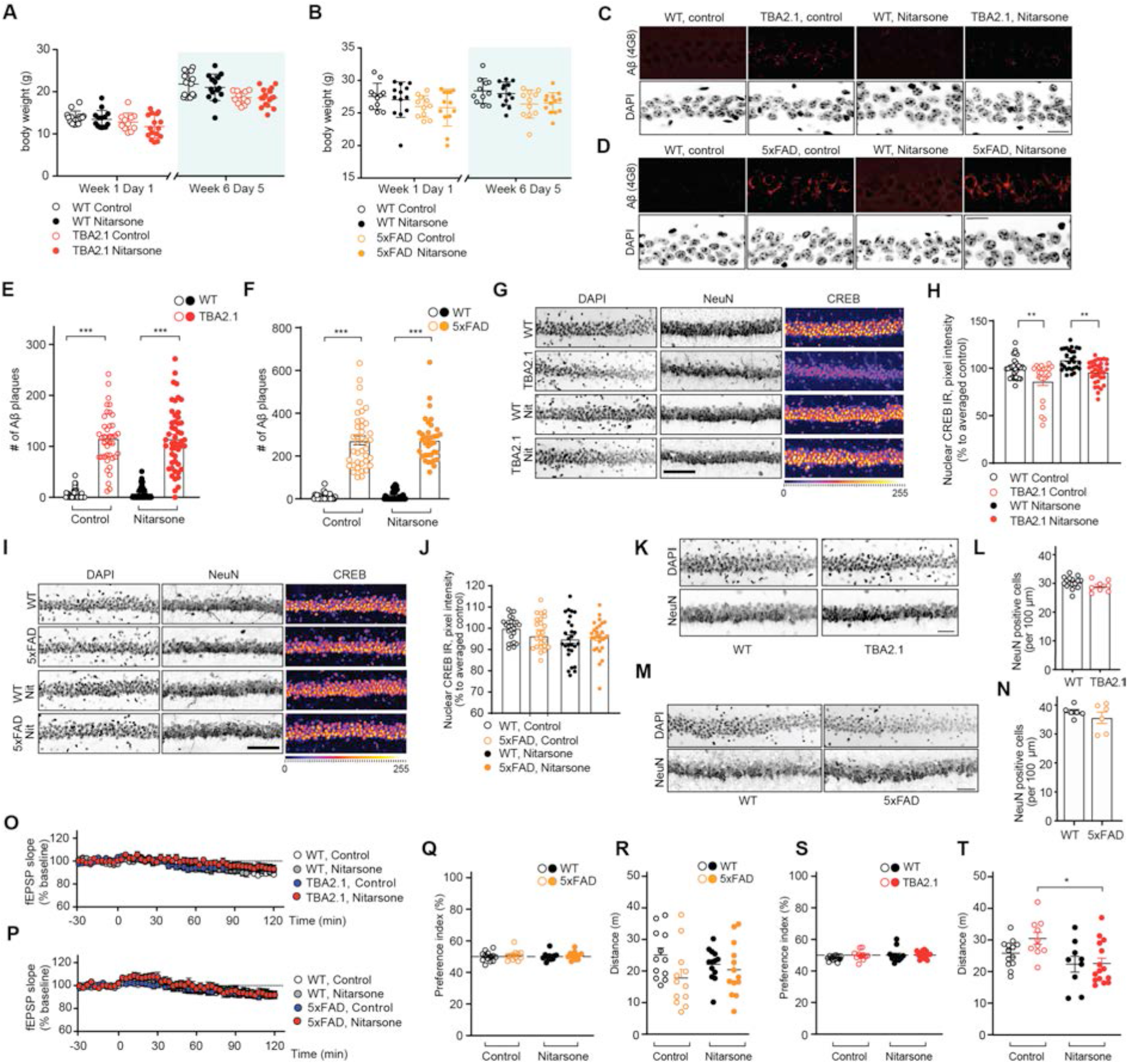
Treatment with Nitarsone rescues AD related phenotypes in vivo. **(A, B)** Nitarsone treatment did not affect body weight of (A) TBA2.1 (N=22-24 animals per group) or (B) 5xFAD mice (N=11-13 animals per group). **(C-F)** Nitarsone treatment does not change amyloid load in (C, E) TBA2.1 and (D, F) 5xFAD mice. (C, D) Confocal images averaged from two sections of the molecular layer of CA1 labelled for amyloid-β (4G8 antibody) and co-stained with DAPI. Scale bar: 100 µm. (E, F) Bar plots representing the number of amyloid-β positive puncta. (E) TBA2.1 N=40-50 CA1 regions 6-9 animals per genotype and (F) 5xFAD N=33-40 CA1 regions 6-7 animals per genotype. **(G, H)** Nitarsone rescues the reduction of CREB immunoreactivity in NeuN positive cells in CA1 of TBA2.1 mice. (G) Representative confocal images of CA1 cryosections from 11 weeks old mice stained for NeuN, DAPI, and CREB. Scale bar: 10 µm. (H) Bar plot of CREB nuclear staining intensity. N=21-34 hippocampal sections from 6-9 animals. **(I, J)** Nitarsone does not affect CREB immunoreactivity in NeuN positive cells in CA1 of TBA2.1 mice. (I) Representative confocal images of CA1 cryosections from 18 weeks old mice stained for NeuN, DAPI, and CREB. Scale bar: 10 µm. (J) Bar plot of CREB nuclear staining intensity. N=28-34 hippocampal sections from 6-7 animals. **(K, L)** TBA2.1 mice do not display neuronal loss at the beginning of the Nitarsone treatment. (K) Representative confocal images of distal CA1 cryosections from 4 weeks old mice stained for NeuN, DAPI, and CREB. Scale bar: 10 µm. (L) Bar graph representing the average number of NeuN-positive cells normalized to WT treated with vehicle. N=8-16 hippocampal sections from 2-3 animals **(M, N)** 5xFAD mice do not display neuronal loss at the end of the Nitarsone treatment. (M) Representative confocal images of distal CA1 cryosections from 19 weeks old mice stained for NeuN, DAPI, and CREB. Scale bar: 10 µm. (N) Bar graph representing the average number of NeuN-positive cells normalized to WT treated with vehicle. N=6 hippocampal sections from 2 animals **(O, P)** Basal synaptic transmission is not affected by bath application of Nitarsone in (O) TBA2.1 and (P) 5xFAD mice. TBA2.1: N=14-18 slices from 5-6 mice and 5xFAD: N=17-18 slices from 6 mice. **(Q, R)** (Q) Nitarsone treatment does not influence preference index and (R) distance travelled during open field arena exploration of TBA2.1 mice. **(S, T)** (S) Nitarsone treatment does not influence preference index and (T) slightly normalizes increased distance travelled during open field arena exploration of 5xFAD mice. *p<0.05, ****p<0.0001 by two-way ANOVA followed by Bonferroni’s multiple comparisons test. All data are represented as mean ± SEM.

